# Vitamin D promotes DNase1L3 to degrade ecDNA and inhibit the malignant progression of hepatocellular carcinoma

**DOI:** 10.1101/2023.08.30.555532

**Authors:** Heng Zhang, Lu-ning Qin, Qing-qing Li, Ting Wu, Lei Zhang, Kai-wen Wang, Shan-bin Cheng, Yue Shi, Yi-qian Feng, Jing-xia Han, Yi-nan Li, Zhi-yang Li, Hui-juan Liu, Tao Sun

**Affiliations:** State Key Laboratory of Medicinal Chemical Biology and College of Pharmacy, Nankai University, Tianjin, China; Tianjin International Joint Academy of Biomedicine, Tianjin, China

**Author notes:** **Correspondence:** Tao Sun, **Email:**. Contributed equally (Heng Zhang, Lu-ning Qin).

**Keywords:** Vitamin D, DNase1L3, ecDNA, HCC, Lipid droplets

## Abstract

Extrachromosomal DNA (ecDNA) is an important carrier of oncogene amplification. However, the degradation mechanism of ecDNA is not well understood. We found that endogenous natural molecular vitamin D (VD) reduces ecDNA and inhibits the progression of hepatocellular carcinoma (HCC). VD reduces ecDNA depending on its transporter GC, which interacts with the endonuclease DNase1L3 and stabilize its protein level. DNase1L3 with its lipophilic region on N-terminus exhibiting an affinity towards lipid droplets (LDs) demonstrates direct degradation effect on ecDNA. Intranuclear LDs are abundantly distributed around ecDNA, DNase1L3 therefore shows an affinity for ecDNA owing to its lipophilic region. VD, as a lipid-soluble molecule, can increase the size of LDs and improve the degradation of DNase1L3 on ecDNA. Consequently, we designed two mRNA-based therapeutics, DNase1L3 and GC-DNase, both of which had an anti-tumor effect on PDX models. The above results showed that treatments targeting ecDNA in cancer are prospective in clinical practice.

## Introduction

The DNA of the vast majority of eukaryotic cells is located on the chromosome of the nucleus. However, as early as the 20th century, scientists discovered that DNA is also present outside of chromosome, such as mitochondrial DNA, plasmids in yeast, and double minutes in cancer cells^1^. The DNA outside of chromosome is circular and named extrachromosomal circular DNA (ecDNA), but its formation and function have only been studied in detail in recent years. In 2017, Paul S. Mischel et al. found that a large number of oncogenes in cancer cells are amplified in the form of ecDNA, which are large enough to carry intact genes ^2^. In 2019, researchers further proved that ecDNA in cancer displays significantly enhanced chromatin accessibility and high transcription levels^3, 4^. In 2020, the research group used pan-cancer whole genome sequencing data for in-depth analysis and found that ecDNA amplification frequently occurs in most of cancer types, and patients whose cancers carried ecDNA had significantly shorter survival^5^. In 2023, Luebeck et al. reported that ecDNA may be present in precancerous tissues, helping to drive non-cancerous tissues to carcinoma, and may serve as a predictive marker for tumors^6^. Thus, ecDNA has important clinical significance.

However, the balancing mechanism by which cells respond to ecDNA, i.e., how ecDNA is degraded within cells, is unclear. The formation of micronuclei has been suggested as a potential mechanism to promote ecDNA removal. In addition, since the presence of ecDNA in cancer cells is positively correlated with tumor immune evasion, the elimination of ecDNA through the action of the immune system is another possible pathway^7^. On the basis of the obvious clinical significance of ecDNA, whether ecDNA can become a target for anti-cancer drugs should be studied. Nevertheless, the drugs that can target ecDNA are currently unknown. It was believed that the unique features of ecDNA provided at least five potential strategies for therapy^8^, but none have yet been developed. Through drug screening, we found that vitamin D (VD) can reduce ecDNA levels in hepatocellular carcinoma (HCC) cells, and further research found that DNase1L3 can directly degrade ecDNA.

VD is an inexpensive and readily available pharmacologically active molecule that, in addition to playing a role in bone metabolism and calcium homeostasis, has a role in cancer treatment, especially in breast, prostate, and colon cancers^9^. Clinical studies have confirmed that the higher serum concentration of VD, the lower the risk of cancer^10^. Moreover, in 2019, the results of a clinical META analysis of 6,537 patients showed that VD supplementation reduced cancer mortality^11^. In terms of HCC, high serum concentration of VD is associated with prolonged survival in HCC^12^. Cellular and animal experiments have confirmed that VD also inhibits breast and prostate cancer, as well as leukemia and lymphoma. However, the molecular mechanism of VD antitumor activity is unclear.

This work studied the anti-tumor effect of VD and the degradation of ecDNA, revealed the pharmacological role of VD from a new perspective, and provided ideas for drug development targeting ecDNA. The study revealed that DNase1L3, whose function is dependent on phase catalysis by microlipid droplets (microLDs) in the nucleus, has the capability to degrade ecDNA, demonstrating characteristics akin to tumor suppressor genes. Moreover, it exhibited promise in the development of biological therapeutics targeting ecDNA. To assess the potential clinical impacts, two mRNA-based drugs, DNase1L3 and fusion protein of GC with the enzymatically active region of DNase1L3 (GC-DNase), were specifically designed and evaluated using patient-derived xenograft (PDX) models in this study.

## Results

### VD reduces ecDNA level in HCC cells

To screen for pharmacologically active molecules capable of decreasing the ecDNA level, we selected some common pharmacologically active molecules that have antitumor effects for metaphase chromosome spread assay and ecDNA was quantified by image analysis with the ecSeg algorithm^13^. These pharmacologically active molecules included 25-hydroxy vitamin D3 (25OHD, the main form of VD in peripheral blood), Vitamin C, Sodium butyrate, Metformin, and natural molecules such as Artemisinin, Puerarin, Baicalein, Myricetin, Hesperetin, Salidroside, Lentinan, Sodium pyruvate, β-Glucan, Apigenin, and Resveratrol. We found that 25OHD, Vitamin C, Sodium butyrate, Metformin, Artemisinin, Puerarin, Baicalein, Myricetin, Hesperetin, Salidroside, Lentinan showed varying degrees of reduction in ecDNA level after treatment, with 25OHD having the most pronounced effect, while Sodium pyruvate, β-Glucan, Apigenin, and Resveratrol have no effect on ecDNA contents (Figure S1A, B). Continually, we proceeded to study the pharmacological effects of VD. Different forms of VD including Vitamin D3 (VD3), 25OHD, and 1,25-dihydroxyvitamin D (1,25(OH)_2_D) were selected to evaluate the specific effect of the active isomer of VD on decreasing the ecDNA levels, as shown in Figure S1C and D, VD3 and 25OHD had an effect on decreasing the ecDNA levels, while 1,25(OH)_2_D revealed almost no effect. Consequently, the follow-up cell experiments were conducted with 25OHD, specifically, 1,25(OH)_2_D was employed as an inactive isomer of VD in reducing ecDNA in HCC, serving as a negative control. Additionally, VD3 was utilized in the animal experiments.

To study the effect of VD on ecDNA, we used the metaphase chromosome spread method to image the chromosomes of HCC cells treated with 25OHD at different concentrations; ecDNA was then quantified using ecSeg. The results indicated 25OHD could reduce the amount of ecDNA in a dose-dependent manner in all three HCC cell lines (Figure 1A, B). Notably, the PLC-PRF-5 cell line exhibited the most significant decrease in ecDNA content after treatment with 25OHD. The PLC-PRF-5 cell line was therefore employed as a model to further study the dose- and time-response relationship of 25OHD reducing ecDNA. As shown in Figure 1C and Figure S1E, 25OHD could reduce ecDNA in PLC-PRF-5 cells on a time-dependent basis, and the ecDNA content almost no longer decreased after 48 h. Therefore, unless otherwise specified, the processing time of 25OHD in subsequent experiments was generally 48 h. Additionally, As shown in Figure 1D and Figure S1F, VD reduced ecDNA in PLC-PRF-5 cells in a concentration-dependent manner. At a concentration of 500 nM, the ecDNA level reached the lowest, and further reduction was negligible, so unless otherwise specified, two concentrations of 100 and 500 nM for cell treatment were selected. We further performed Circle-seq^14^ on PLC-PRF-5 cells treated with 25OHD and the experimental procedure has been shown in Figure 1E. Meanwhile, the matched whole-genome sequencing (WGS) was performed. ecDNA was identified by the combined analysis of Circle-seq data and WGS data^15, 16^. As shown in Figure 1F, the length of ecDNA in the control group ranged from 100 ∼ 10^6^ bp, with most of them distributed in the range of 100 kb ∼ 1 Mb. 1674 ecDNAs were detected in the control group, and the ecDNAs detected after 100 nM and 500 nM 25OHD treatments decreased to 21 and 17, which indicated that 25OHD resulted in a significant decrease in the number of ecDNAs. Moreover, 25OHD not only resulted in a significant decrease in the number of ecDNAs, but also the length of the ecDNAs showed a trend of getting shorter. The function of these downregulated ecDNAs by 25OHD were related to Epithelial-mesenchymal transition (EMT), proliferation, cell cycle, focal adhesion, and other malignant behaviors (Figure 1G). From these oncogenes carried on ecDNA, we selected two typical oncogenes (*BRAF*, *KRAS*) associated with HCC, and *CENPF*, which had poor prognosis (Figure S1G) and higher circle score, serving as representative genes for further study. As shown in Figure 1H, we generated circular maps of the three ecDNAs with different size (*BRAF*, 23.32 Mb, *KRAS*, 4.24 Mb, *CENPF*, 611 kb) by the combined analysis of Circle-seq data and WGS data^15^. Fluorescence in situ hybridization (FISH) in metaphase chromosomes and interphase cells was further conducted to evaluate the amplification of the aforementioned oncogenes and to determine the intranuclear location of the three genes. The three FISH probes targeting the full-length gene sequence in combination with matched chromosomes of centromeric probes were used for the FISH procedure. To set up a positive control, junctions of ecDNA were found from the Circle-seq data (Table. S1) and CRISPR-Cas9 plasmid specifically cutting the junction of *BRAF* was designed. After 25OHD treatment, the FISH signals of all three genes were significantly reduced (Figure 1I, J and Figure S1H-K). Many clustered signals were found outside the chromosomes in the control group, reflecting a high abundance of ecDNA, whereas the signals in the 25OHD group were mainly located on the chromosomes (Figure 1I, J and Figure S1H, J). Consistently, the FISH signals of the three genes were also significantly reduced in interphase PLC-PRF-5 cells after 25OHD treatment (Figure 1I, J and Figure S1I, K), indicating that 25OHD could reduce ecDNA in HCC cells. We then imaged *KRAS* ecDNA in live cells by CRISPR live FISH method (Yi et al., 2022b). As shown in movie S1-2 and Figure S1L, a gradual decrease in the size and magnitude of the signal of the intracellular *KRAS* ecDNA was found after 25OHD treatment. The average signal intensity of *KRAS* ecDNA in the cells of the DMSO control group fluctuated in a small range, while in the 25OHD group it showed a continuous and significant decrease. Notably, 25OHD induced ecDNA degradation at a rapid rate, which was observed in a very short period of time. These data suggest that 25OHD can cause a continuous and steady decrease in ecDNA in living cells. The result was also verified by qPCR of ecDNA junctions and the level of all three oncogenes on ecDNA decreased significantly after treatment of 25OHD (Figure 1K and Figure S1M). The corresponding mRNA (Figure 1L and Figure S1N) and protein levels (Figure S1O) of these oncogenes also showed a downward trend. The above results showed that 25OHD could play a role in reducing ecDNA in HCC cells, as well as decrease the transcription and translation of malignant behavior-related genes including *BRAF*, *KRAS*, and *CENPF*.

**Figure 1.**
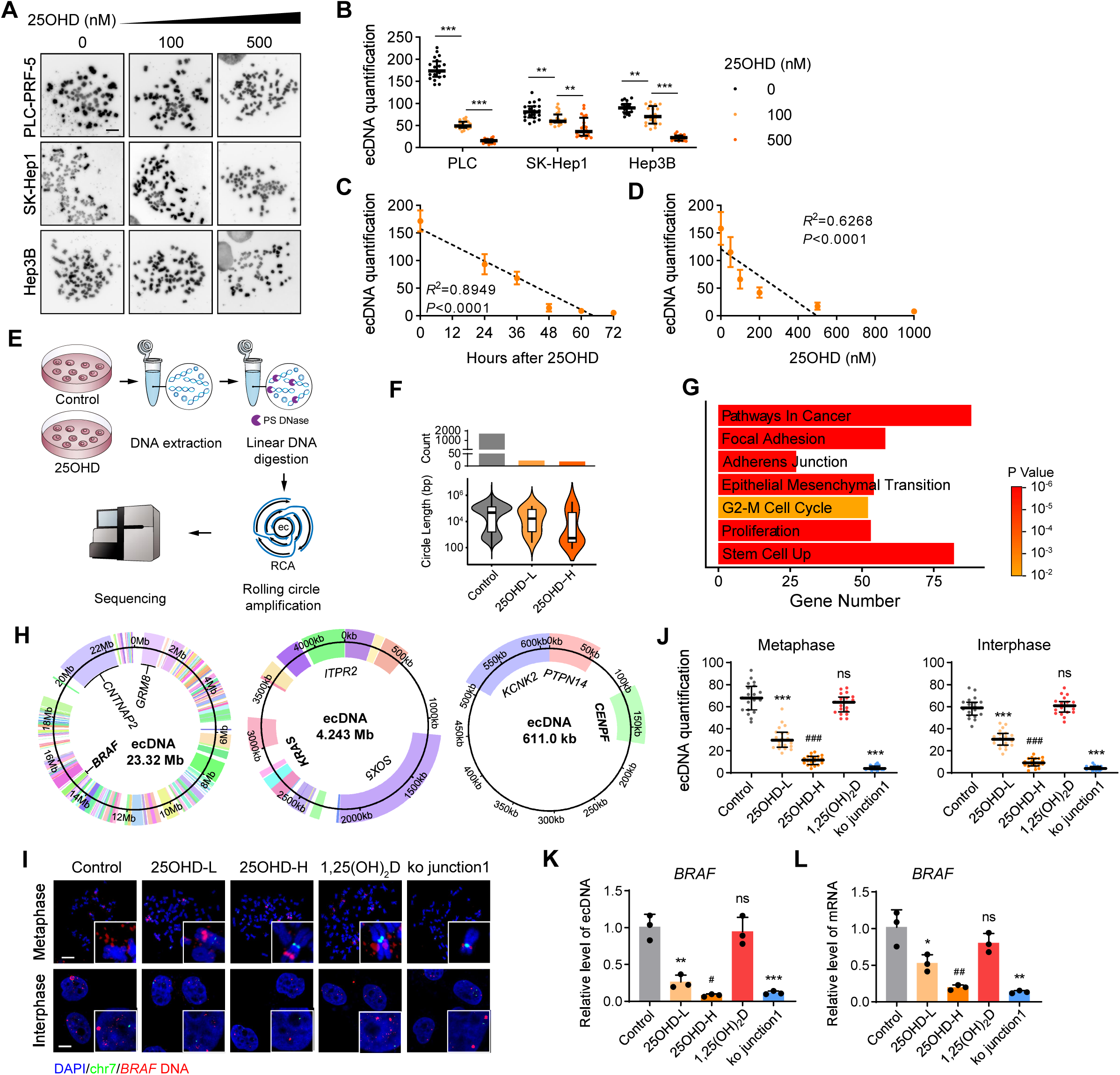
VD reduced ecDNA level in HCC cells. **(A and B)** Representative images (**A**) and quantification (**B**) of metaphase ecDNA by ecSeg in HCC cells after treatment of DMSO or 25OHD (100 nM or 500 nM) for 48 h. Scale bar, 10 µm. 20 metaphase spreads from 3 biologically independent samples were counted. Statistics were calculated on biological replicates with Wilcoxon rank-sum test. \*\**P* < 0.01, \*\*\**P* < 0.001. Error bars show median, upper and lower quartiles. **(C)** Quantification of metaphase ecDNA by ecSeg in HCC cells after treated with DMSO or 25OHD (500 nM) for indicated time points. 20 metaphase spreads from 3 biologically independent samples were counted. Statistics were calculated on biological replicates with simple linear regression. **(D)** Quantification of metaphase ecDNA by ecSeg in HCC cells after treated with different concentrations of 25OHD for 48 h. 20 metaphase spreads from 3 biologically independent samples were counted. Statistics were calculated on biological replicates with simple linear regression. **(E)** Schematic diagram of isolation, purification and sequencing of ecDNA. **(F)** ecDNAs identified from combined analysis of Circle-seq and WGS showed a decrease of number and length. Control DMSO, 25OHD-L (100 nM), 25OHD-H (500 nM). **(G)** Gene annotation and GO analysis of the down-regulated ecDNA in the 25OHD (500 nM, 48 h) treatment group compared to the control group (DMSO). **(H)** The circular ecDNA plots of BRAF, KRAS and CENPF structure. **(I)** Representative *BRAF* FISH images of metaphase ecDNA (top) and interphase ecDNA (bottom) signal in PLC-PRF-5 cells after treatment of DMSO, 25OHD-L (100 nM), 25OHD-H (500 nM), 1,25(OH)_2_D (500 nM) or transfected with plasmid specifically ko ecDNA junction of *BRAF* for 48 h. Scale bar, 10 µm. **(J)** FISH signal quantification by ecSeg. Scale bar, 10 µm. 20 metaphase spreads or interphase cells of FISH images from 3 biologically independent samples were counted. Statistics were calculated on biological replicates with Wilcoxon rank-sum test. \*\*\**P* < 0.001, ns, not significant. Error bars show median, upper and lower quartiles. **(K)** QPCR detected of ecDNA levels of *BRAF* purified and isolated from PLC-PRF-5 cells treated with DMSO, 25OHD-L (100 nM), 25OHD-H (500 nM), 1,25(OH)_2_D (500 nM) or transfected with specifically ko ecDNA junction of *BRAF* plasmid for 48 h. *n*=3, biological replicates. Statistics were calculated on biological replicates with two-tailed unpaired t-tests. \*\*\**P* < 0.001, ns, not significant. Error bars show mean with SD. **(L)** Quantification of mRNA level of *BRAF* by qPCR in PLC-PRF-5 cells. Student’s *t*-test, *n*=3, biological replicates. Statistics were calculated on biological replicates with two-tailed unpaired t-tests. \*\*\**P* < 0.001, ns, not significant. Error bars show mean with SD.

### VD inhibits the malignant progression of HCC through ecDNA

Building on the findings of Figure 1 that the ecDNA reduced by 25OHD was closely related to a series of malignant behaviors of HCC cells, we further verified the effect of 25OHD on inhibiting the malignant progression of HCC *in vitro* and *in vivo*. The cells did not undergo significant apoptosis after 25OHD treatment, as indicated by the results of Annexin V-PI flow cytometry (Figure S2A). We further studied whether 25OHD functions by inhibiting the proliferative capacity or EMT of HCC. The results of scanning electron microscopy (SEM) and E-cadherin/Vimentin immunofluorescence (IF) staining showed an increase in adhesion of HCC cells and an enhancement in E-cadherin expression after 25OHD treatment (Figure 2A). Wound healing assay (Figure 2B and Figure S2B) and Transwell assay (Figure 2C and Figure S2B) revealed that 25OHD could reduce the migration and invasion ability of HCC cells on a dose-dependent basis. Additionally, fluorescent gelatin degradation assay indicated that 25OHD dose-dependently inhibited the ability of degrading the extracellular matrix of HCC cells (Figure 2D). RNA-seq was conducted, to further evaluate the global effect of 25OHD on HCC cells. The results showed that the mRNA levels of 2491 genes were significantly downregulated after 25OHD treatment (Figure 2E). Upon annotation, these downregulated genes were found to be associated with malignant behaviors such as invasion, EMT, and enhanced stemness (Figure 2F). Extrachromosomal circularization and amplification are associated with increased oncogene expression^17^. To demonstrate the correlation between the down-regulatory effect of 25OHD on oncogene mRNA expression levels and ecDNA, we further analyzed Circle-seq and mRNA-seq data. The oncogene expressions identified including *BRAF*, *KRAS*, and *CENPF* using mRNA-seq revealed that 25OHD suppressed the oncogenes coincided with ecDNAs, as confirmed by Circle-seq, which also showed significant down-regulation of these ecDNAs after 25OHD treatment, suggesting that ecDNAs contribute to oncogene expression (Figure 2G). Moreover, with all mRNAs were sorted by its expression (FPKM) in Control (in grey) or 25OHD (in orange) group, and the ecDNA amplified genes that were reduced after 25OHD treatment were marked with dots, the results show that the mRNA expression level of oncogenes amplified on ecDNA are inhibited by 25OHD. Specifically, the gray dots are mainly clustered in the upper position, indicating that the expression of these ecDNA-amplified genes is higher in the control group. The orange dots have a more obvious downward shift in position, suggesting that the expression of these ecDNA-amplified genes is overall suppressed after 25OHD treatment (Figure 2H). Therefore, the downregulation of mRNA in these genes was related to the reduction of ecDNA. We further detected markers of the relevant malignant phenotype in 25OHD-treated HCC cells, western blot results revealed a decrease in the levels of mesenchymal markers such as vimentin, ZEB1 and Twist1, whereas the protein levels of epithelial marker E-cadherin and Occludin were significantly increased (Figure S2C).

**Figure 2.**
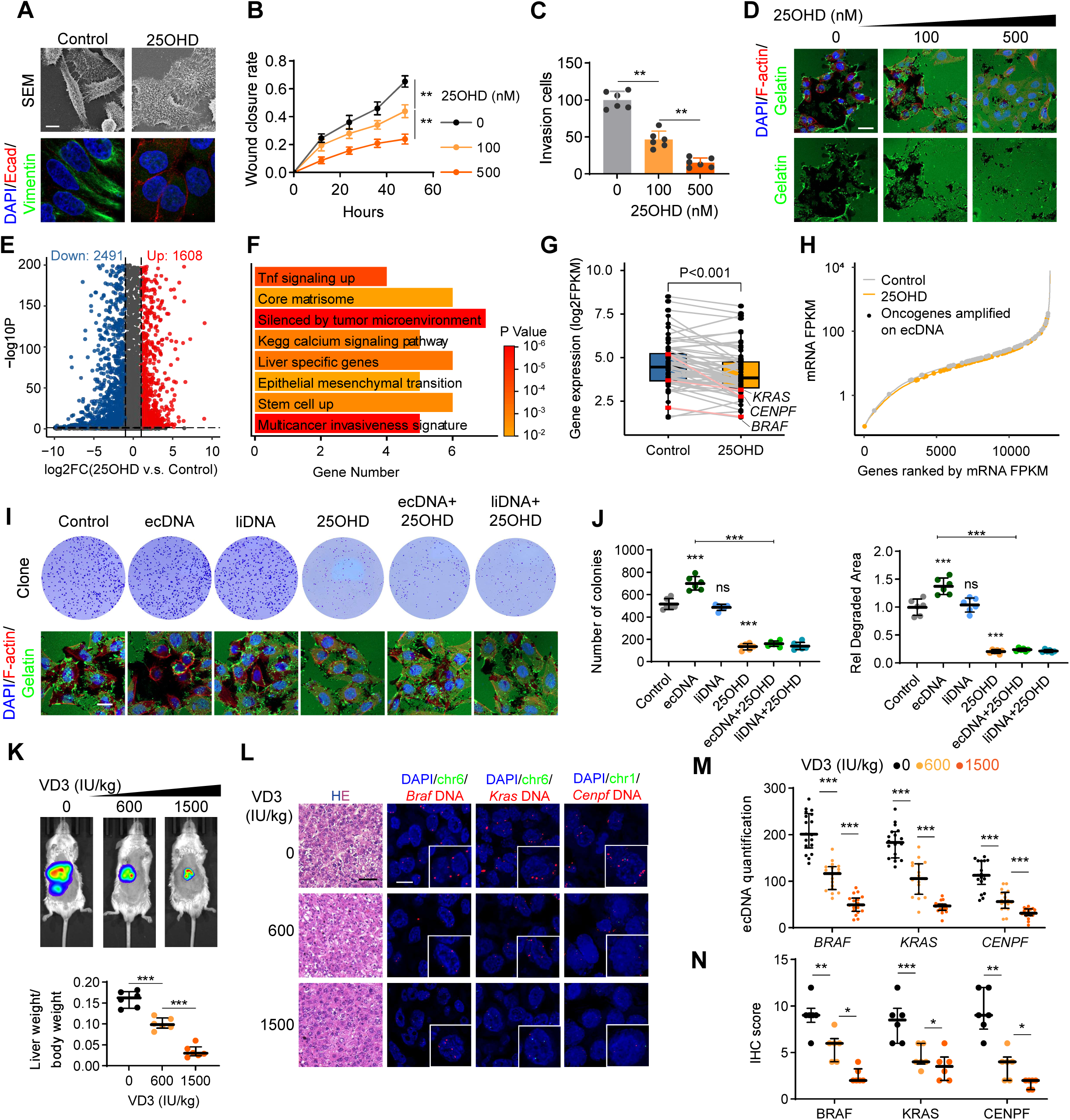
VD inhibits the malignant progression of HCC through ecDNA. **(A)** The morphologic detect with scanning electron microscopy (SEM) and Immunostaining of E-cadherin and vimentin signal in PLC-PRF-5 cells. Scale bar, 10 µm. **(B)** Quantification of wound closure rate in wound healing assay of PLC-PRF-5 cells treated with DMSO or 25OHD (100 nM or 500 nM) for 48 h. *n*=3, biological replicates. Statistics were calculated on biological replicates with two-tailed unpaired *t*-tests, \*\**P* < 0.01. Error bars show mean with SD. **(C)** Quantification of invasion cells in Transwell assay of PLC-PRF-5 cells treated with DMSO or 25OHD (100 nM or 500 nM) for 48 h. *n*=6, biological replicates. Statistics were calculated on biological replicates with two-tailed unpaired *t*-tests, \*\**P* < 0.01. Error bars show mean with SD. **(D)** Fluorescent gelatin degradation and phalloidin/DAPI staining of PLC-PRF-5 cells treated with DMSO or 25OHD (100 nM or 500 nM) for 48 h. Scale bar, 40 µm. **(E)** Volcano plot showing up- and down-regulation of mRNA after 25OHD (500 nM, 48 h) treatment. **(F)** Gene annotation of 25OHD (500 nM, 48 h) downregulated mRNA. **(G)** The expression of genes on VD down regulated ecDNAs in Control group (DMSO, 48 h, blue) compared to the VD (25OHD, 500 nM, 48 h, orange) group. **(H)** All genes were ranked based on their mRNA-seq expression levels (FPKM), with those amplified in extrachromosomal DNA (ecDNA) marked as dots. **(I)** Representative images of colony formation (top) and fluorescent gelatin degradation and phalloidin/DAPI staining (bottom) of PLC-PRF-5 cells with indicated treatments. Control: treated with DMSO, ecDNA: delivery of purified ecDNA, liDNA: delivery of linearizing ecDNA. VD: treated with 500 nM 25OHD, ecDNA+VD: delivery of purified ecDNA and treated with 500 nM 25OHD, liDNA+VD: delivery of linearizing ecDNA and treated with 500 nM 25OHD. For the colony formation assay, the treatments were for 10 days. For fluorescent gelatin degradation assay, the treatments were for 48 h. Scale bar, 40 µm. **(J)** Quantification of the number of colonies (left) and relative degraded area (right) in PLC-PRF-5 cells. *n*=6, biological replicates. Statistics were calculated on biological replicates with two-tailed unpaired *t*-tests, \*\**P* < 0.01, \*\*\**P* < 0.001, ns, not significant. Error bars show mean with SD. **(K)** Representative *in vivo* images (top) of H22-luc-tumor-bearing-BALB/c mice with treatments of Ethanol or VD3 (600 IU/kg or 1500 IU/kg) for 2 months, and liver weight/body weight ratio was quantified (bottom). *n*=6, biological replicates. Statistics were calculated on biological replicates with two-tailed unpaired *t*-tests, \*\*\**P* < 0.001. Error bars show median, upper and lower quartiles. **(L and M)** Representative image of HE staining and DNA FISH for liver tissues in H22-luc-tumor-bearing-BALB/c mice (**L**) and quantification of FISH signal by ecSeg (**M**). Scale bar, 10 µm. Average fluorescence intensity in the nucleus in 18 cells of FISH images from 6 biologically independent samples were measured. Statistics were calculated on biological replicates with Wilcoxon rank-sum test, \*\*\**P* < 0.001. Error bars show median, upper and lower quartiles. **(N)** Quantification of Immunohistochemical staining of BRAF, KRAS, CENPF in liver tissues of H22-luc-tumor-bearing-BALB/c mice. *n*=6, biological replicates. Statistics were calculated on biological replicates with unpaired *t*-test with Welch’s correction, \**P* < 0.05, \*\**P* < 0.01, \*\*\**P* < 0.001. Error bars show median, upper and lower quartiles.

To determine whether the effect of 25OHD on cell function depends on ecDNA, ecDNA was firstly purified from PLC-PRF-5 cells following the published procedure^18^. The DNA was then subjected to linearization by sequential treatment of nickase fnCpf1 and single strand DNA-specific nuclease S1^19^, which was used as linear amplicons (liDNA) for control. Extracted ecDNA was stained with YOYO1 and then imaged by SIM to visualize the integrity of the ecDNA circular structure (Figure S2D), and the sensitivity of ecDNA and liDNA to exonucleases Plasmid-Safe ATP-dependent DNase was verified to further validate the integrity of ecDNA circles and successful linearization of liDNA (Figure S2E). Equal amounts of ecDNA and liDNA were delivered into PLC-PRF-5 cells using extracellular vesicle technologies for cellular delivery of large molecular weight DNA ^20^ (Figure S2F). The increase in ecDNA level shown by metaphase spread assay verified the successful delivery of ecDNA (Figure S2G). As shown in Figure 2I and J, the malignant phenotypes such as proliferation and invasion of PLC-PRF-5 cells were enhanced after delivery of ecDNA but not liDNA. The treatment of 25OHD to cells delivered with ecDNA, significantly counteracted the increase in malignant phenotype, indicating that the effect of 25OHD on the malignant phenotype of cells was ecDNA-dependent. Furthermore, a pair of ecDNA low/high cell lines, MHCC-97L and MHCC-97H, were selected to verify the effect of 25OHD on cell function was ecDNA dependent. MHCC-97L and MHCC-97H are a pair of low/high metastatic HCC homologous cell lines from the same patient^21^, and the ecDNA level was correlated with malignancy. The low metastatic cell line MHCC-97L contained a very low level of ecDNA, whereas the high metastatic cell line MHCC-97H contained a significantly higher level of ecDNA (Figure S2H). Notably, FISH images of *KRAS* showed oncogene amplification on ecDNA in MHCC-97H cells, while in MHCC-97L cells the oncogene amplified in HSR (Figure S2H). Functional experiments were performed separately on the two cell lines. The results showed that 25OHD treatment on MHCC-97L cell line showed no significant changes in malignant behavior such as proliferation and invasion, however, MHCC-97H cells were significantly affected by 25OHD in malignant behaviors (Figure S2I, J). These results suggested that the inhibitory effect of 25OHD on the malignant phenotype of HCC cells is related with ecDNA. To verify the effect of ecDNAs including *BRAF*, *KRAS* and *CENPF* on cellular function, CRISPR-Cas9 plasmids specifically cutting the ecDNA junctions were designed (the effectiveness of plasmid knockout was validated in Figure S10E) and transfected with PLC-PRF-5 cells. Through clone formation and gelatin degradation assay, we observed different degrees of suppression of malignant phenotypes such as cell proliferation or invasion (Figure S2K, L), which suggested that these oncogenes amplified on ecDNA are closely associated with malignant behaviors. Next, we examined the antitumor effect of VD *in vivo* using the H22 orthotopic xenograft model. The results showed that VD3 inhibited the growth and metastasis of H22 xenograft tumor (Figure 2K). FISH was performed to evaluate ecDNA levels at tumor sites in mouse livers, consistent with the *in vitro* results, VD3 treated mice have significantly less ecDNA levels (Figure 2L, M) of the three oncogenes, and immunohistochemistry (IHC) results showed a downward trend in their protein levels (Figure 2N and Figure S2M). The above results showed that VD could inhibit malignant behaviors such as proliferation, migration and invasion in HCC, and the anti-tumor effect of VD was related to ecDNA.

### VD increases the nuclear GC content and facilitates its interaction with DNase1L3

We further investigated the mechanism by which 25OHD reduced ecDNA. Firstly, according to the chemical structure of 25OHD, we applied three target prediction algorithms including SEA^22^, SwissTargetPrediction^23^, and HITPICK^24^ to predict the possible pharmacologically active molecules targets of 25OHD. All three algorithms predicted the VDR and GC proteins with high confidence (Figure 3A), which were supported by some research^9^. Therefore, we knocked out VDR or GC using the CRISPR-Cas9 system (the effectiveness of plasmid knockout was validated in Figure S10A, B) to determine whether 25OHD treatment can still reduce ecDNA. After knocking out GC, but not VDR, 25OHD could not reduce ecDNA (Figure 3B), suggesting that 25OHD reduced ecDNA mainly depends on GC. Interestingly, 1,25(OH)_2_D has been reported primarily activated VDR but could bind extremely weakly to GC^25^, which further supporting the idea that 25OHD reduces ecDNA by GC rather than VDR. After 25OHD treatment, we found the upregulation of GC content in the nucleus (Figure 3C, D). Western blot of whole cell lysate and nuclear lysate also showed an increase in intranuclear GC content after 25OHD treatment (Figure 3E). The mechanism by which 25OHD upregulates GC levels intracellularly has been reported in our previous work^26^. Based on this finding, we considered the degradation of ecDNA was most likely related to the increase of GC protein in the nucleus. We then performed pulldown assay and mass spectrometry (MS) to identify the interacting proteins of the intranuclear GC protein after 25OHD treatment, from which DNase1L3 (Figure 3F and Table. S2) was identified. The coimmunoprecipitation (CoIP) results also confirmed that GC interacted with DNase1L3 intracellularly after the treatment of 25OHD (Figure 3G). Proximity ligation assay (PLA) results further showed a significant increase of intracellular interaction between GC and DNase1L3 after 25OHD treatment (Figure 3H, I). Since DNase1L3 has been reported to be related to apoptosis^19^, to avoid this phenomenon being caused by apoptosis, TUNEL staining was performed. The negative signal for TUNEL staining positive signal of PLA indicated that the PLA signals were not false positives due to apoptosis (Figure S3A). Furthermore, we analyzed the dynamic process, free energy change, and stable conformation of 25OHD-promoted GC-DNase1L3 interactions by molecular dynamics simulations (Figure 3J-L). The Root Mean Square Deviation (RSMD) fluctuations were smaller during the kinetic simulation compared with the solvent control system, indicating that the binding system was more stable during the kinetic simulation (Figure 3J). The parameters of free energy, van der Waals force energy, and binding free energy of GC-DNase1L3 interactions were lower for the system containing 25OHD (Figure 3K), suggesting that the presence of 25OHD in the system led to more stable GC-DNase1L3 interactions. The images of the GC-DNase1L3 stable conformation of the intercalation region obtained from dynamic simulations are shown in Figure 3L. The presence of 25OHD increased the amino acid residues of GC-DNase1L3 intercalation, and the distance between the two α-helices was brought closer by 2.5 Å, indicating that GC-DNase1L3 bound more tightly. To further verify the effect of 25OHD on the interactions between GC and DNase1L3, Surface plasmon resonance (SPR) and microscale thermophoresis (MST) experiments were performed. As shown in Figure S3B, the response value of DNase1L3 was also very low at the highest concentration of GC in the PBS solution system. However, in the 25OHD solution system, the response value of DNase1L3 increased with increasing GC concentration. This finding indicates that GC and DNase1L3 protein interactions increased under 25OHD environment. *In vitro* MST (Figure S3C) also showed that GC did not enhance the thermal surge of DNase1L3 in PBS environment. However, the thermal surge of DNase1L3 in 25OHD environment increased with increasing GC concentration, indicating that 25OHD promoted the interaction between DNase1L3 and GC.

**Figure 3.**
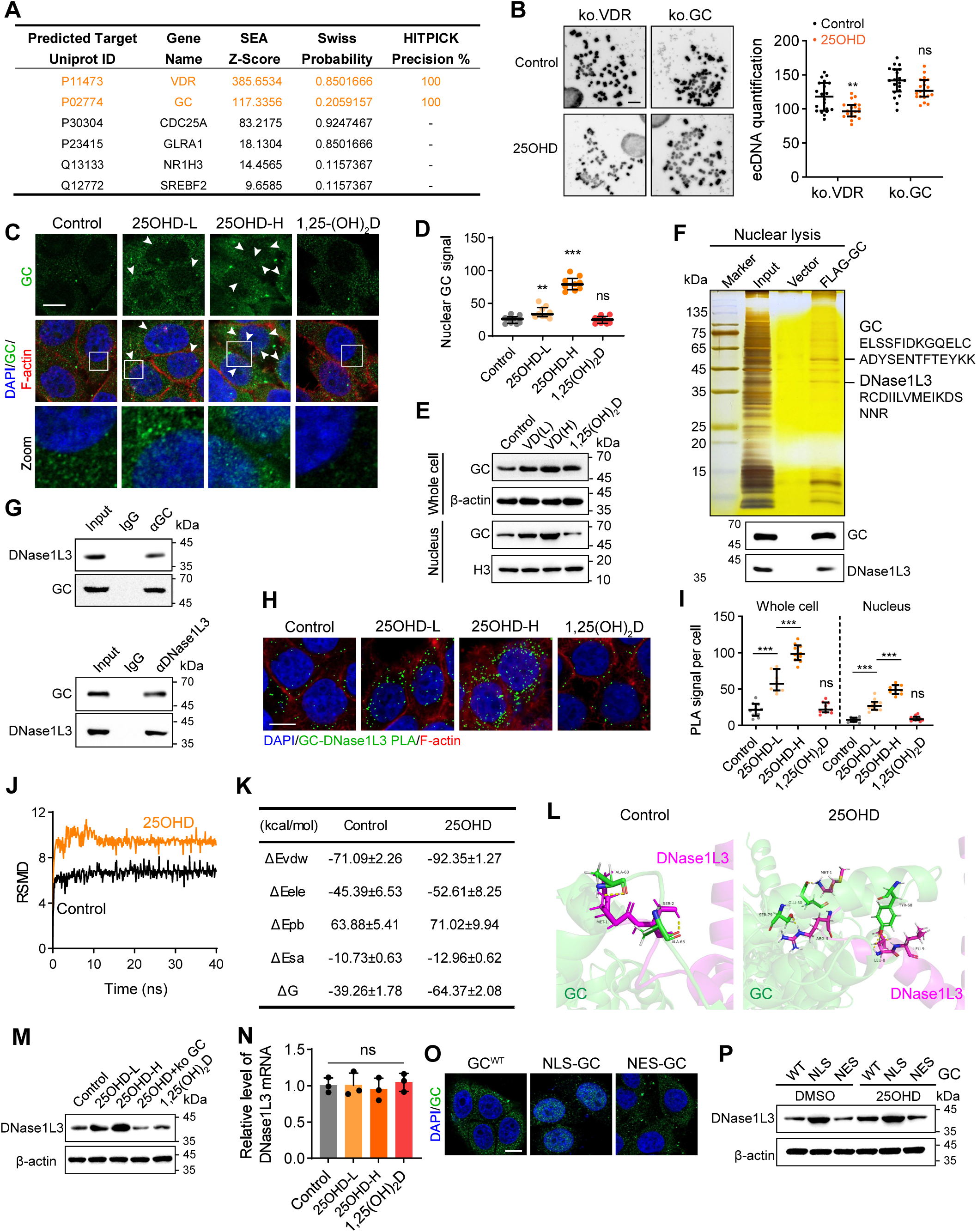
VD increases the nuclear GC content and facilitates its interaction with DNase1L3. **(A)** Prediction of VD targets through SEA, Swiss, and HITPICK. **(B)** Representative image (left) and quantification by ecSeg (right) of metaphase ecDNA in PLC-PRF-5 cells knocking out of VDR or GC and treated with DMSO or 25OHD (500 nM) for 48 h. Scale bar, 10 µm. 20 metaphase spreads from 3 biologically independent samples were counted. Statistics were calculated on biological replicates with unpaired *t-* test with Welch’s correction. \*\*\**P* < 0.001, ns, not significant. Error bars show median, upper and lower quartiles. **(C and D)** Immunostaining of GC signal (**C**) and quantification (**D**) of nuclear GC signal in PLC-PRF-5 cells treated with DMSO or 25OHD (500 nM) for 48 h. Scale bar, 10 µm. *n*=10, biological replicates. Statistics were calculated on biological replicates with two-tailed unpaired *t*-tests. \*\**P* < 0.01, \*\*\**P* < 0.001; ns, not significant. Error bars show median, upper and lower quartiles. **(E)** Western blot analysis of GC expression level in the whole cell lysis and nucleus lysis of PLC-PRF-5 cells treated with DMSO, VD(L) (100 nM), VD(H) (500 nM) or 1,25(OH)_2_D (500 nM) for 48 h. **(F)** Silver staining (top) of proteins acquired by Flag-GC pull-down in nuclear lysis from PLC-PRF-5 cells treated with 25OHD (500 nM) for 48 h and western blot analysis (bottom) of GC and DNase1L3. **(G)** Co-immunoprecipitation assays in PLC-PRF-5 cells treated with 25OHD (500 nM) for 48 h with anti-GC followed by immunoblotting (IB) with antibodies against DNase1L3 and GC or with anti-DNase1L3 followed by IB with anti-GC and DNase1L3. **(H and I)** PLA assays with GC and DNase1L3 in PLC-PRF-5 cells treated with DMSO, VD(L) (100 nM), VD(H) (500 nM) or 1,25(OH)_2_D (500 nM) for 48 h **(H)** Scale bar, 10 µm. Quantification of PLA signal in whole cell and nucleus were shown **(I)** *n*=10, biological replicates. Statistics were calculated on biological replicates with two-tailed unpaired *t*-tests. \*\*\**P* < 0.001; ns, not significant. Error bars show median, upper and lower quartiles. **(J)** The Root Mean Square Deviation (RSMD) during the molecular dynamic simulation shows that the 25OHD (500 nM)-containing system has a more stable binding of the two proteins during the molecular dynamics simulation. **(K)** Changes in Van der Waals potential energy (ΔEvdw), electrostatic potential energy (ΔEele), polar interaction potential energy (ΔEpb), solvation potential energy (ΔEsa) and binding free energy (ΔG) during molecular dynamics simulation, showing that the binding of GC to DNase1L3 protein is more stable in the VD-containing system. **(L)** The stable structure of GC binding to DNase1L3 protein obtained by molecular dynamic simulation with or without VD. **(M)** Western blot analysis of DNase1L3 in ko GC PLC-PRF-5 cells with treatment of VD(H) (500 nM) for 48 h or wild type PLC-PRF-5 cells treated with DMSO, VD(L) (100 nM), VD(H) (500 nM) or 1,25(OH)_2_D (500 nM) for 48 h. **(N)** The level of DNase1L3 mRNA was detected by RT-qPCR in PLC-PRF-5 cells treated with DMSO, VD(L) (100 nM), VD(H) (500 nM) or 1,25(OH)_2_D (500 nM) for 48 h. *n*=3, biological replicates. Statistics were calculated on biological replicates with two-tailed unpaired *t*-tests. ns, not significant. Error bars show mean with SD. **(O)** Representative confocal images after transfection with GC^WT^-EGFP, NLS-GC-EGFP or NES-GC-EGFP for 48 h. Scale bar, 10 µm. NLS: nuclear localization signals (MKRVLVLLLAVAFGHA). NES: nuclear export signals (MNLVDLQKKLEELELDEQQ). **(P)** Western blot analysis of DNase1L3 expression level in PLC-PRF-5 cells treated with DMSO or 25OHD (500 nM) and indicated transfection for 48 h. WT: transfection of empty vector, NLS: transfection of NLS-GC plasmid, NES: transfection of NES-GC plasmid.

On the basis of the above results, we believe that GC was activated by 25OHD, which accelerated the shuttle to the nucleus and increased the chance of interaction with DNase1L3. We further investigated whether mRNA and protein levels of DNase1L3 were affected by 25OHD and GC. The results showed that 25OHD elevated DNase1L3 protein level, but it was unable to induce an increase in the protein level following the knockout of GC (Figure 3M). However, the mRNA level of DNase1L3 was not affected with 25OHD treatment (Figure 3N). We also investigated whether elevated protein levels of DNase1L3 are associated with nuclear localization of GC. Wildtype (WT) GC, nuclear localization signal (NLS)-GC and nuclear export signal (NES)-GC were designed and verified by IF (Figure 3O). We found that the protein level of DNase1L3 in the NLS-GC group were higher than that in the GC^WT^ or NES-GC group. In the overexpression GC group with 25OHD treatment, the protein level of DNase1L3 continued to increase while no increase was observed in the NES-GC group (Figure 3P). This result suggested that nuclear localization of GC may be an important link in 25OHD elevating protein levels of DNase1L3. Furthermore, we found that the increased GC level in the nucleus was related to the reduction of ecDNA (Figure S3D).

### GC interacts with USP25 to promote the deubiquitination of DNase1L3

Previous results have shown that 25OHD treatment increases levels of the DNase1L3 protein, but not mRNA, suggesting that 25OHD did not affect the transcription of DNase1L3. Thus, we further investigated whether 25OHD affects the synthesis or degradation of the DNase1L3 protein. After the addition of protein synthesis inhibitor cycloheximide (CHX), 25OHD and NLS-GC were also able to increase DNase1L3 protein levels, whereas the addition of proteasome inhibitors MG132 did not affect DNase1L3 levels neither in 25OHD treatment group nor NLS-GC group (Figure S4A). We therefore concluded that 25OHD inhibited the degradation of DNase1L3, resulting in the accumulation of DNase1L3. Given that the protein degradation pathway relies primarily on the ubiquitin–proteasome pathway, we investigated whether 25OHD affects the ubiquitination of DNase1L3. The results showed a low level of ubiquitination of DNase1L3 in the 25OHD and NLS-GC groups; after knockout of GC, 25OHD could not reduce the ubiquitination of DNase1L3, which suggested that 25OHD reduced DNase1L3 ubiquitination by GC (Figure S4B). As this effect is associated with nuclear localization of GC, we studied the interaction protein of GC and observed deubiquitinase USP25 in MS data (Figure S4C and Table. S2). We further validated the interaction between GC and USP25 through CoIP (Figure S4D) and PLA (Figure S4E, F). With the suspicion that the GC in the nucleus can recruit USP25 adjacent to DNase1L3 so that the ubiquitination of DNase1L3 was removed, we further investigated whether USP25 can deubiquitinate DNase1L3. As shown in Figure S4G-I, after overexpression of USP25, the ubiquitination of DNase1L3 was reduced and the half-life significantly improved. However, the ubiquitination level of DNase1L3 increased (Figure S4J) and the degradation became faster after knocking out USP25 (Figure S4K, L), indicating USP25 deubiquitinates DNase1L3 thereby stabilizing its protein level.

### DNase1L3 degrades ecDNA *in vitro* and *in vivo*

We found that GC interacted with DNase1L3 upon activation by 25OHD, thereby increasing its content in the nucleus. Thus we examined whether DNase1L3 is closely linked to the degradation of ecDNA. Firstly, we found that transfection of DNase1L3 plasmid or protein reduced the content of ecDNA, whereas an increased level of ecDNA was detected after knocking out DNase1L3 (Figure 4A, B). We further studied the FISH signals of *BRAF*, and the results also revealed that the ecDNA content decreased after transfection of plasmid or protein of DNase1L3 in both metaphase spreads and interphase HCC cells (Figure 4C-E). The results of qPCR of isolated ecDNA were also consistent with the data of FISH (Figure 4F). The rest of the two oncogenes (*KRAS* and *CENPF*) were also verified in Figure 4G-I and Figure S5A and B. To confirm that these results were caused by the direct interaction of DNase1L3 with ecDNA, excluding indirect effects, we also designed *in vitro* experiments for verification. After the extraction of whole DNA, Plasmid-Safe ATP-dependent DNase was added to eliminate linear chromosomal DNA to purify ecDNA. DNase1L3 protein was added to the reaction containing ecDNA with the presence of Ca^2+^ and Mg^2+^ and incubated at 37 °C; the DNA concentration was found to be greatly reduced (Figure 4J). We further studied found that the quantitative efficacy (Figure 4K) and timeliness (Figure 4L) of DNase1L3 were significantly linearly correlated (*R*²>0.75, *P*=0.0001). Nearly 90% of ecDNA was degraded after 1 h of 40 ng/μL DNase1L3 or 2 h of 10 ng/μL DNase1L3, but the degradation efficiency was low when the solution system was alkaline (pH=8) (Figure 4M). We also examined the degradation preference of DNase1L3 on liDNA versus ecDNA. As shown in Figure 4N, DNase1L3 degraded both forms of DNA, and the degradation effect for ecDNA was better for the same incubation time and concentration. Here, we also made a comparison among DNase1 (a human endonuclease), micrococcal Nuclease (MNase, an endonuclease from Staphylococcus aureus), and DNase1L3 to evaluate their respective degradation effects on ecDNA. As shown in Figure 4O, the degradation of ecDNA by DNase1 was weaker than that by DNase1L3, and the degradation of ecDNA by MNase, a widely used enzyme in ChIP experiments, was the strongest. Moreover, the levels of ecDNA exhibited only a slight decrease in response to two treatments, the presence of the mutant DNase1L3 (R206C) and the addition of zinc chloride to the reaction system, both of which effectively inhibited the nuclease activity of DNase1L3.

**Figure 4.**
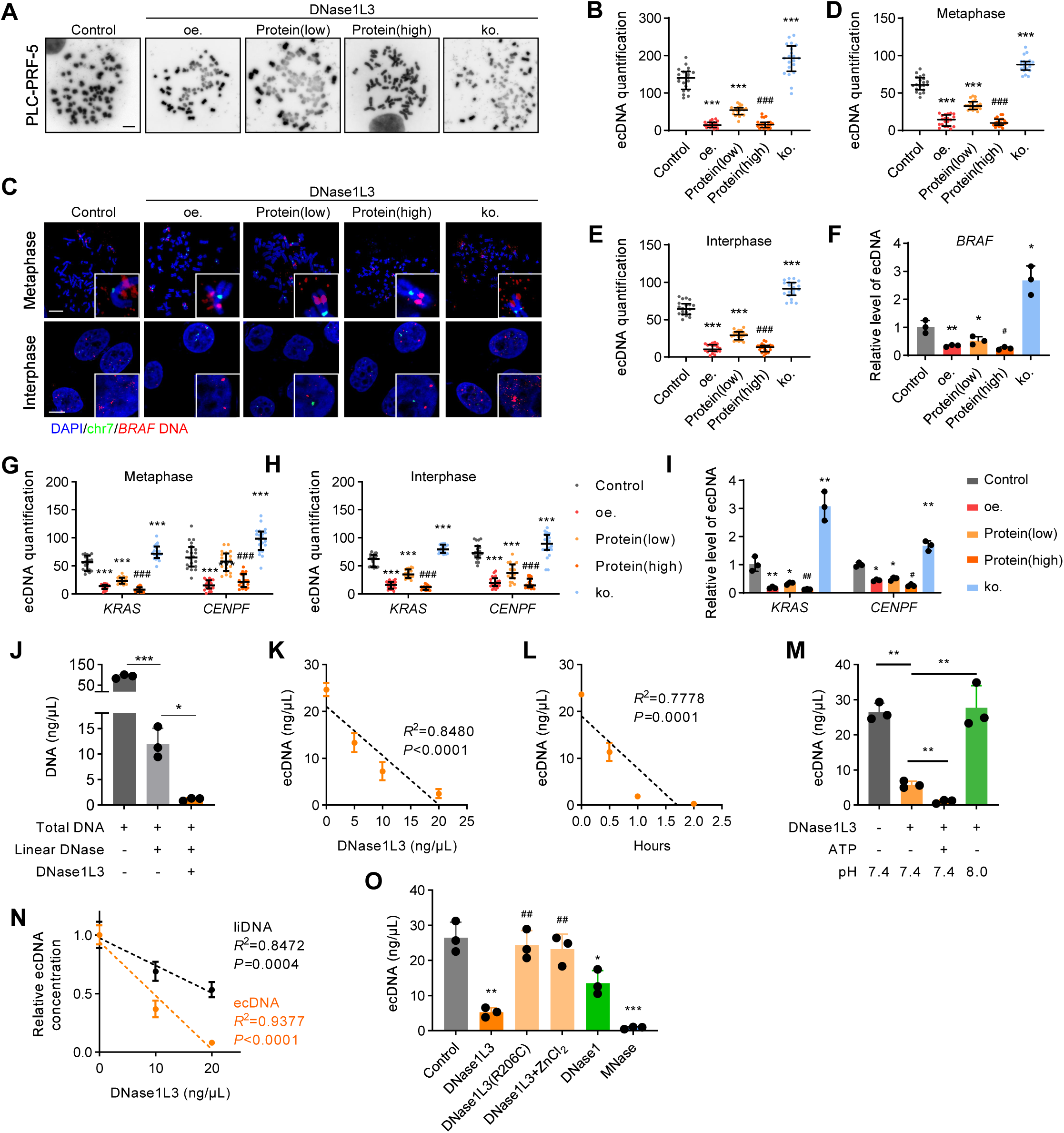
DNase1L3 degrades ecDNA *in vitro* and *in vivo*. **(A and B)** Representative images (**A**) and quantification (**B**) of metaphase ecDNA by ecSeg in PLC-PRF-5 cells after indicated treatment. Control: transfection of empty vector, oe: over expression of DNase1L3 plasmid, Protein(low): transfection of DNase1L3 protein (2 µg), Protein(high): transfection of DNase1L3 protein (10 µg), ko: transfection of DNase1L3 ko plasmid. Scale bar, 10 µm. 20 metaphase spreads from 3 biologically independent samples were counted. Statistics were calculated on biological replicates with Wilcoxon rank-sum test. \*\*\**P* < 0.001, compared with control unless otherwise indicated. #*P* < 0.05, compared with protein (low) unless otherwise indicated. Error bars show median, upper and lower quartiles. **(C to E)** Representative *BRAF* FISH images of metaphase ecDNA (top) and interphase ecDNA (bottom) signal in HCC cells after indicated treatments, Scale bar,10 µm (**C**). FISH signal quantification by ecSeg (**D and E**). 20 metaphase spreads or interphase cells of FISH images from 3 biologically independent samples were counted. Statistics were calculated on biological replicates with Wilcoxon rank-sum test. \*\*\**P* < 0.001, compared with control unless otherwise indicated. ###*P* < 0.001, compared with protein (low) unless otherwise indicated. Error bars show median, upper and lower quartiles. **(F)** QPCR detected of ecDNA levels of *BRAF* purified and isolated from PLC-PRF-5 cells with indicated treatments. *n*=3, biological replicates. Statistics were calculated on biological replicates with two-tailed unpaired t-tests, \**P* < 0.05, \*\**P* < 0.01, compared with control unless otherwise indicated. #*P* < 0.05, compared with protein (low) unless otherwise indicated. Error bars show mean with SD. **(G and H)** Quantification of FISH signal of *KRAS*, *CENPF* in metaphase (**G**) and interphase (**H**) in PLC-PRF-5 cells with indicated treatments. 20 metaphase spreads or interphase cells of FISH images from 3 biologically independent samples were counted. Statistics were calculated on biological replicates with Wilcoxon rank-sum test. \*\*\**P* < 0.001, compared with control unless otherwise indicated. ###*P* < 0.001, compared with protein (low) unless otherwise indicated. Error bars show median, upper and lower quartiles. **(I)** QPCR detected of ecDNA levels of *CENPF, KRAS* purified and isolated from PLC-PRF-5 cells with indicated treatments. *n*=3, biological replicates. Statistics were calculated on biological replicates with two-tailed unpaired t-tests. \**P* < 0.05, \*\**P* < 0.01, compared with control unless otherwise indicated. #*P* < 0.01, compared with protein (low) unless otherwise indicated. Error bars show mean with SD. **(J)** Quantification of ecDNA in PLC-PRF-5 cells after treated with plasmid-safe ATP-dependent DNase or co-incubation with DNase1L3 protein (10 ng/μL) at 37 °C for 1 h. *n*=3, biological replicates. Statistics were calculated on biological replicates with two-tailed unpaired t-tests. \**P* < 0.05, \*\*\**P* < 0.001. Error bars show mean with SD. **(K)** Quantification of purified ecDNA in PLC-PRF-5 cells after incubated with different concentrations of DNase1L3 protein (5 ng/μL, 10 ng/μL,20 ng/μL) at 37 °C for 1 h. *n*=3, biological replicates. Statistics were calculated on biological replicates with simple linear regression. Error bars show mean with SD. **(L)** Quantification of purified ecDNA in PLC-PRF-5 cells after incubated with DNase1L3 protein (10 ng/µl) at 37 °C of indicated time points (0.5 h, 1 h, 2 h). *n*=3, biological replicates. Statistics were calculated on biological replicates with simple linear regression. Error bars show mean with SD. **(M)** Quantification of purified ecDNA incubated with the indicated treatments at 37 °C for 1 h. DNase1L3 (10 ng/μL), ATP (the final concentration is 1 mM). *n*=3, biological replicates. Statistics were calculated on biological replicates with two-tailed unpaired t-tests. \*\**P* < 0.01. Error bars show mean with SD. **(N)** Quantification of ecDNA and linear DNA (liDNA) incubated with DNase1L3 protein (10 ng/μL, 20 ng/μL) at 37 °C for 1 h. *n*=3, biological replicates. Statistics were calculated on biological replicates with simple linear regression. Error bars show mean with SD. **(O)** Quantification of purified ecDNA incubated with the indicated treatments at 37 °C for 1 h. *n*=3, biological replicates. DNase1L3 (10 ng/μL), DNase1L3 (R206C) (10 ng/μL), DNase1 (5 μL), MNase (5 μL), ZnCl_2_ (1 mM), the incubation was performed in a 60 μL reaction volume in the presence of 5 mM MgCl_2_, 2.5 mM CaCl_2_. Statistics were calculated on biological replicates with two-tailed unpaired t-tests. \**P* < 0.05, \*\**P* < 0.01, \*\*\**P* < 0.01, compared with Control. ##*P*<0.01, compared with DNase1L3. Error bars show mean with SD.

### The affinity of DNase1L3 towards lipid droplets is through its lipophilic regions, enabling the degradation of ecDNA surrounded by lipid droplets

We further investigated the spatial properties of DNase1L3 to degrade ecDNA. Imaging with structured light illumination microscopy (SIM, Figure 5A), it showed that *BRAF* ecDNA in PLC-PRF-5 cells was clustered rather than diffuse, which was consistent with the ecDNA distribution characteristics reported previously^27^. DNase1L3 and ecDNA were highly co-localized after 25OHD treatment, with a colocation coefficient of more than 0.7. In particular, large DNase1L3 signals were also found in ecDNA hubs. Fluorescence statistics of the ecDNA-enriched regions showed that the signals of ecDNA and DNase1L3 in the two groups always appeared at the same time. The ecDNA signal observed in the control group exhibited greater intensity compared to DNase1L3, whereas in the 25OHD group, it displayed reduced intensity in comparison to DNase1L3 (Figure 5B). Additionally, the ecDNA signal was greatly reduced after treatment of 25OHD (Figure 5C). The presence of the DNase1L3 signals adjacent to ecDNA suggested that 25OHD may have helped DNase1L3 better recognize and degrade ecDNA. The lipid-soluble nature of 25OHD prompted us to investigate the charge distribution of DNase1L3. As shown in Figure 5D, it can be observed that beyond the enzymatically functional region of DNase1L3, there exists an electrically neutral domain at the N-terminal (NT) that exhibits lipophilic characteristics. Therefore, we further studied whether degrading ecDNA by DNase1L3 is related to lipophilicity.

**Figure 5.**
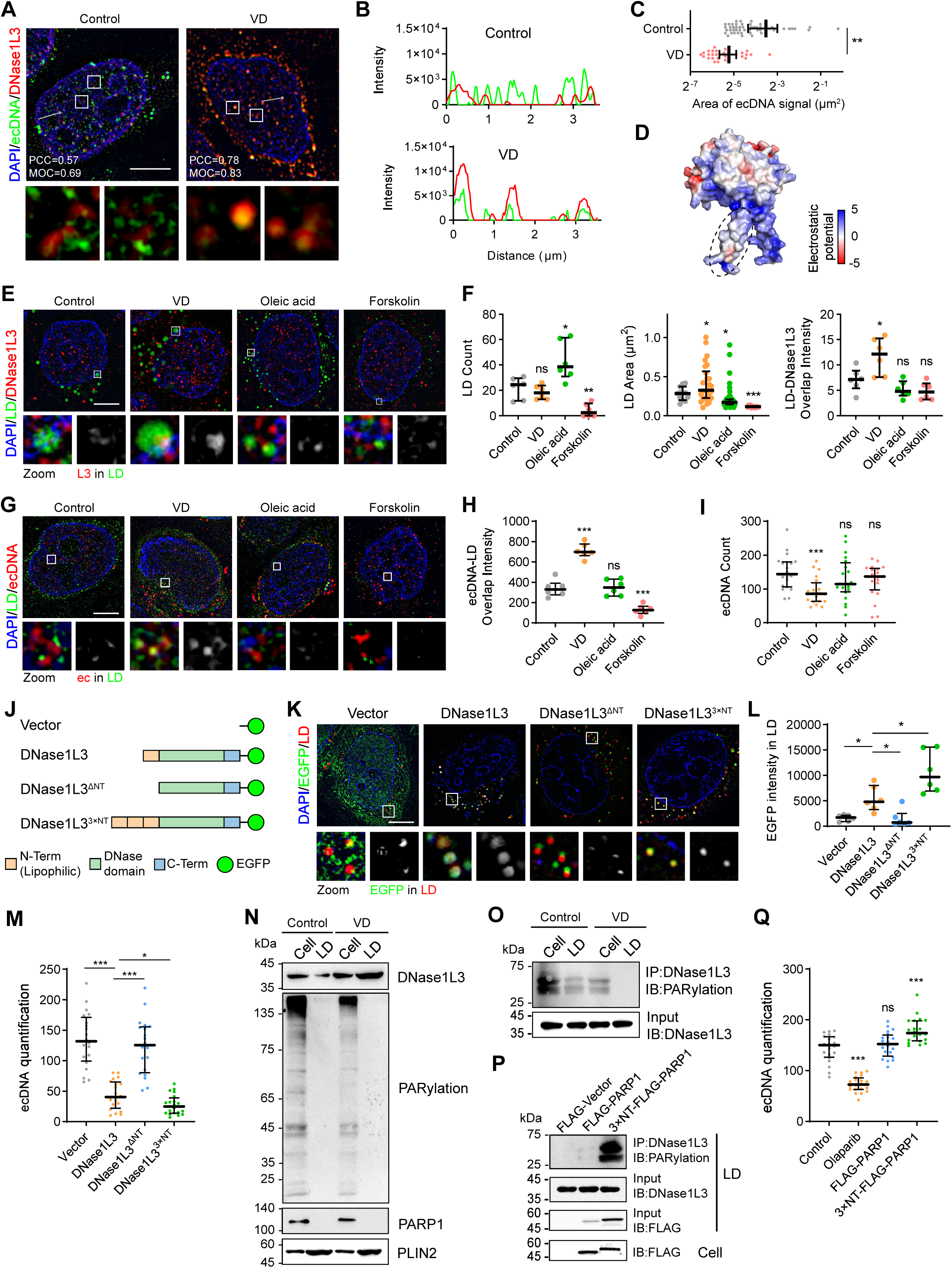
The affinity of DNase1L3 towards lipid droplets is through its lipophilic regions, enabling the degradation of ecDNA surrounded by lipid droplets. **(A)** Representative SIM images of ecDNA (indicated by *BRAF* DNA FISH, green) combined with Immunostaining of DNase1L3 signal (red) in PLC-PRF-5 cells treated with DMSO or 25OHD (500 nM) (top) for 48 h, corresponding zoom views of colocalization analysis were shown (bottom). PCC, Pearson’s correlation coefficient; MOC, Mandas’s overlap coefficient. Scale bar: 5 μm. **(B)** Quantification of the fluorescence intensity of FISH and DNase1L3 signals in PLC-PRF-5 cells treated with DMSO or 25OHD (500 nM) for 48 h. **(C)** Analysis of size of area of ecDNA signal in PLC-PRF-5 cells treated with DMSO or 25OHD (500 nM) for 48 h. At least 29 ecDNA signal from 3 biologically independent samples were counted. Statistics were calculated on biological replicates with Wilcoxon rank-sum test. \*\**P* < 0.01. Error bars show mean with 95% CI. **(D)** Electrostatic potential of DNase1L3 molecular surface generated by APBS Electrostatics Plugin in Pymol software. The elliptical dotted line marks the lipophilic region. **(E)** Representative live-cell SIM images of EGFP-labelled DNase1L3 and BODIPY staining of lipid droplets (top) and corresponding zoom views of the overlap of DNase1L3 and lipid droplets (bottom) in PLC-PRF-5 cells after DMSO, 25OHD (500 nM), oleic acid (200 µM) or forskolin (10 µM) treatments for 48 h. Scale bar: 5 μm. **(F)** Statistics of lipid droplets and DNase1L3 signal in PLC-PRF-5 cells after indicated treatments. *n*=6, biological replicates. Statistics were calculated on biological replicates with two-tailed unpaired t-tests. \**P* < 0.05, \*\**P* < 0.01, \*\*\**P* < 0.001, ns, not significant, compared with control unless otherwise indicated. Error bars show median, upper and lower quartiles. **(G)** Representative SIM images of *BRAF* DNA FISH and Bodipy staining of lipid droplets in PLC-PRF-5 cells after DMSO, 25OHD (500 nM), oleic acid (200 µM) or forskolin (10 µM) treatments for 48 h. Scale bar: 5 μm. **(H and I)** Statistics of overlap of ecDNA and LD, *n*=6, biological replicates (**H**) and ecDNA FISH signal of 20 interphase FISH images in PLC-PRF-5 cells from 3 biologically independent samples were counted after indicated treatments(**I**). Statistics were calculated on biological replicates with Wilcoxon rank-sum test. \*\*\**P* < 0.001, ns, not significant, compared with control. Error bars show median, upper and lower quartiles. **(J)** Schematic of the different DNase constructs used. **(K)** Representative live-cell SIM images of EGFP-labelled different DNase1L3 constructs and BODIPY staining of lipid droplets (top) and corresponding zoom views of the overlap images (bottom) in PLC-PRF-5 cells after transfection of indicated DNase1L3 constructs. Scale bar: 5 μm. **(L)** Quantification of the lipid droplets containing EGFP intensity in PLC-PRF-5 cells after transfection of indicated plasmids. *n*=6, biological replicates. Statistics were calculated on biological replicates with unpaired *t* test with Welch’s correction. \**P* < 0.05. Error bars show median, upper and lower quartiles. **(M)** Quantification of the number of ecDNA in PLC-PRF-5 cells after transfection of indicated plasmids. *n*=6, biological replicates. Statistics were calculated on biological replicates with Wilcoxon rank-sum test. \**P* < 0.05, \*\*\**P* < 0.001. Error bars show median, upper and lower quartiles. **(N)** PARylation of the whole cell proteins and LD binding protein in PLC-PRF-5 cells treated with DMSO or 25OHD (500 nM) for 48 h were monitored by immunoblot analysis with antibodies to PAR. PARP1, DNase1L3 and classical LD binding protein PLIN2 in the whole cell lysis and LD component were measured respectively. **(O)** Western blot analysis of PARylation of DNase1L3 in the whole cell lysis and LD component in PLC-PRF-5 cells treated with DMSO or 25OHD (500 nM) for 48 h. **(P)** Co-immunoprecipitation assays in LD component from PLC-PRF-5 cells after transfection of PARP1 or 3×NT-PARP1 overexpression plasmids with anti-DNase1L3 followed by immunoblotting with antibodies against PAR. **(Q)** Quantification of metaphase ecDNA by ecSeg in PLC-PRF-5 cells after transfection of Flag-PARP1, 3×NT-PARP1 or treated with PARP1 inhibitor Olaparib (20 μM) for 48 h. 20 metaphase spreads from 3 biologically independent samples were counted. Statistics were calculated on biological replicates with Wilcoxon rank-sum test. \*\*\**P* < 0.01, ns, not significant, compared with control unless otherwise indicated. Error bars show median, upper and lower quartiles.

DNase1L3 and lipid droplets (LDs) were co-stained with IF and the chemical dye BODIPY for SIM imaging. As shown in Figure 5E, LDs were observed in the nucleus, where only a few LDs were close in size to the cytoplasmic LDs, and microLDs occupied the majority. To better observe the interaction between LDs and DNase1L3, a magnified image of the LD region (lower left) and the overlapping signals of DNase1L3 and LD (lower right) were provided. In the 25OHD group, the overlapping signal of LDs and DNase1L3 was the strongest. Further statistical analysis was shown in Figure 5F. After 25OHD treatment, there was no significant change in the number of LDs, however, a substantial increase was noted in the size of LDs when compared to the control group. Additionally, a significant enhancement in the signal of DNase1L3 within the LDs was observed. Cells were treated with oleic acid and forskolin which were reported as lipid droplet inducer and inhibitor^28^ respectively, we observed that the number of cytoplasmic LDs and intranuclear LDs increased after oleic acid treatment while a decrease was observed after forskolin treatment. However, neither oleic acid nor forskolin changed the signal intensity of DNase1L3 in the LDs, suggesting that the number of LDs did not affect the interaction with DNase1L3. In conclusion, DNase1L3 is more affinitive for LDs in the present of 25OHD. Since DAPI signals were found in the LD from SIM images, we further investigated correlation of ecDNA and LDs.

We visualized ecDNA containing *BRAF* by FISH while LDs were stained with BODIPY and their signals were observed with SIM. Given that the FISH procedure breaks up large LDs within the cytoplasm, we first confirmed that the microLDs in the nucleus were not affected by the FISH experiment, after the DNA FISH procedure, microLDs within the nucleus were still observed (Figure 5G). *BRAF* FISH results showed that microLDs were adjacent to ecDNA, and there was a strong overlap between microLDs and ecDNA in the 25OHD-treated cells (Figure 5G). However, the oleic acid and forskolin treatments did not significantly affect the overlap between microLDs and the amount of ecDNA (Figure 5H, I), suggesting that the number of LDs did not affect the formation or degradation of ecDNA. The above results implied that LDs might be related to ecDNA degradation. We further investigated whether the action of DNase1L3 is related to LDs, for which we designed three DNase1L3 tracer plasmids shown in Figure 5J: Wild type DNase1L3 (WT), DNase1L3 with lipophilic region deleted (ΔNT), and DNase1L3 with three repeated segments of lipophilic region (3×NT) at the N-terminal. The plasmids were transfected to cells respectively and co-stained with BODIPY to perform live cell imaging by SIM. As shown in Figure 5K and L, DNase1L3^ΔNT^ hardly entered the LDs, whereas DNase1L3^3×NT^ increased the interaction with LDs compared with WT. The results of western blot on DNase1L3 proteins in LDs also coincide with cell imaging (Figure S6A). These results suggested that the NT region of DNase1L3 is crucial for the interaction with LDs, helping DNase1L3 to access ecDNA clusters surrounded by LDs to function as an ecDNA degrader. We also proved that DNase1L3^3×NT^ degraded ecDNA better (Figure 5M).

It has been reported that DNase1L3 plays an important role in the apoptotic process^29^. We conducted additional studies to explore the underlying reasons for the ability of DNase1L3 to cleave DNA while not directly inducing apoptosis in HCC cells. DNase1L3 activity could be inhibited by poly(ADP-ribosyl)ation (PARylation) in DNA degradation^30^. We found that almost no PARylation of DNase1L3 occurred in LDs, but most cellular DNase1L3 was in PARylation state, as determined by western blot and CoIP experiments (Figure 5N-O). PARP1 (the writer of PARylation) with a 3× lipophilic region was transfected into cells, which lead to the induced proximity of PARP1 to DNase1L3 on the LDs, resulting in DNase1L3 on LDs was again modified by PARylation (Figure 5P). Consequently, ecDNA level in the cells was increased (Figure 5Q). This finding suggested that LDs can protect DNase1L3 from PARylation, which may be the key for DNase1L3 to degrade ecDNA around LDs but not genomic DNA under specific spatial and temporal conditions. 25OHD enlarges the size of LDs, increases the amount of DNase1L3 in LD, and also protects DNase1L3 from interacting with PARP1, making DNase1L3 in LDs more susceptible to degrade ecDNA. The results of SIM images (Figure S6B) also support this conclusion.

We further investigated the effect of GC on the interaction of DNase1L3 with LDs. We first observed the localization of GC and DNase1L3 in LDs after 25OHD treatment using IF. The results showed that 25OHD increased the GC and DNase1L3 protein levels in LDs (Figure S6C). We extracted LDs after overexpressing GC in cells and detected the protein level of DNase1L3 in LD. As shown in Figure S6D, the level of DNase1L3 in the equivalent LDs increased after the overexpression of GC. The results indicated that GC interacts with DNase1L3 to increase DNase1L3 binding to LDs. We also constructed the GC-DNase plasmid as shown in Figure S6E, that is, The GC protein was artificially linked to the enzyme-activated domain of DNase1L3 (DNase domain). The SIM results showed that 25OHD enhanced the interaction signal and overlapping area of DNase1L3 with LDs. Additionally, the interaction signal and overlapping area between GC-DNase and LDs were observed to be comparatively stronger than those of DNase1L3 (Figure S6F, G). Therefore, we further performed ecDNA degradation assay (Figure S6H) to simulate LDs environment *in vitro*, i.e., adding liposome to the reaction, the results showed that the simultaneous addition of GC and liposome could substantially enhance the degradation of ecDNA by DNase1L3, consistent with the *in vivo* phenomenon that the GC activation of DNase1L3 is dependent on LDs. The above results showed that 25OHD promoted the recognition of DNase1L3 toward LDs, and this effect was dependent on GC. GC interacted with DNase1L3 after 25OHD treatment, enhancing the lipophilicity of DNase1L3 and facilitating the degradation of ecDNA.

### DNase1L3 inhibits the malignant progression of HCC dependent on lipophilic regions

Given that DNase1L3 has the effect of degrading ecDNA of HCC cells, and this effect depends on its NT lipophilic regions, we further investigated the effect of DNase1L3 in inhibiting the malignant progression of HCC *in vivo* and *in vitro* and determined whether its role is related to the lipophilic region. SEM images showed that the epithelial morphology of HCC cells in the overexpressed DNase1L3, 3×NT group were more pronounced and the adhesion increased compared with the control group. However, ΔNT group did not show obvious changes. The IF images of EMT key markers E-cadherin and Vimentin also showed the same trend (Figure 6A). Thus, DNase1L3 may be able to inhibit malignant behaviors such as EMT in cancer cells, and this effect depends on its NT lipophilic regions. Since DNase1L3 has been reported to be associated with apoptosis^31^, we investigated whether the antitumor effect it exerted in HCC was a consequence of the induction of apoptosis, as indicated by the result of Annexin V-PI flow cytometry in Figure S7A, the cells did not undergo significant apoptosis after over expression or knock out of DNase1L3. Wound healing assay, Transwell assay (Figure 6B, C and Figure S7B), and fluorescent gelatin degradation assay (Figure 6D, E) showed that DNase1L3 could inhibit the migration and invasion ability of HCC cells, but the deletion of NT did not show an obvious effect, indicating that this function was dependent on NT. EMT related marker also showed the same trend of changes (Figure 6F). The pair of ecDNA low/high cell lines were applied to investigate the effect of DNase1L3 on cellular functions including proliferation, migration, and invasion. Figure S7C and D showed that the MHCC-97L cell line with low ecDNA level and with oncogene amplification on HSR after overexpression of DNase1L3 had no significant changes in malignant behavior such as proliferation and invasion. Hence, DNase1L3 is not functionally significant for cancer cells with only HSR amplification in contrast to those with high ecDNA levels. Animal studies of orthotopic liver xenograft tumors also showed that DNase1L3 inhibited the growth of H22 xenograft tumor (Figure 6G, H) and prolonged the survival of tumor-baring mice (Figure 6I), which relied on its NT lipophilic regions. The pathological sections of the tumor site of the mouse liver were subjected to FISH, and the ecDNA level was evaluated. After in situ transplantation of cells overexpressing DNase1L3^3×NT^, the ecDNA levels of *BRAF*, *KRAS*, and *CENPF* were significantly reduced (Figure 6J and Figure S7E), and the corresponding protein levels were also significantly decreased (Figure 6K and Figure S7F). However, no significant differences were found in the overexpression ΔNT group. The above results showed that DNase1L3 degraded ecDNA *in vivo*, exerted antitumor effects, mainly relied on its NT lipophilic regions.

**Figure 6.**
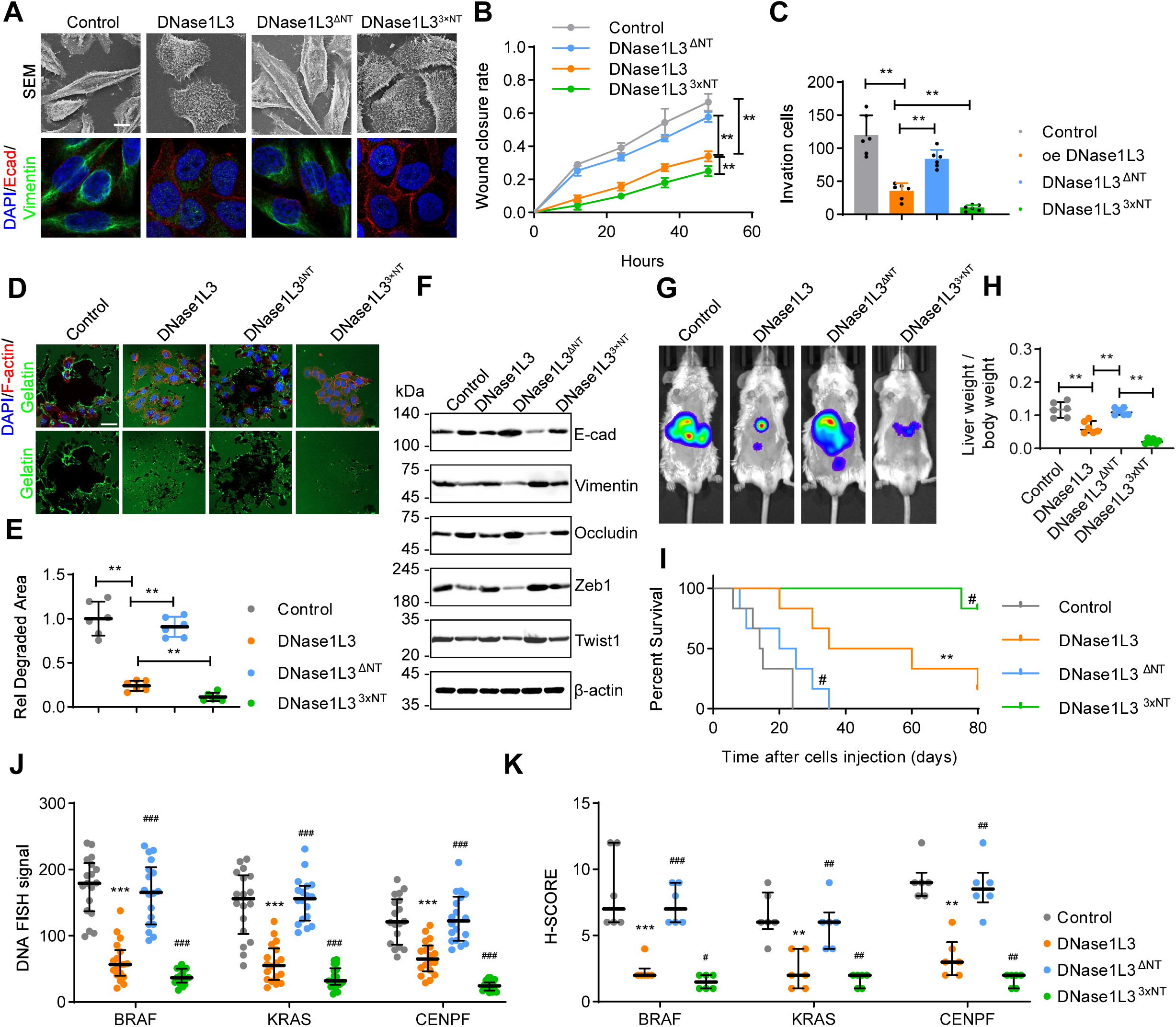
DNase1L3 inhibits the malignant progression of HCC dependent on lipophilic regions. **(A)** The morphologic detect with scanning electron microscopy (top) and Immunostaining (bottom) of E-cadherin and vimentin signals in PLC-PRF-5 cells transfected with DNase1L3, DNase1L3^ΔNT^ or DNase1L3 ^3×NT^ plasmid for 48 h. Scale bar, 10 µm. **(B)** Quantification of wound closure rate in wound healing assay of PLC-PRF-5 cells treated as indicated. *n*=3, biological replicates. Statistics were calculated on biological replicates with two-tailed unpaired *t*-tests, \*\**P* < 0.01. Error bars show mean with SD. **(C)** Quantification of invasion cells in Transwell assay of PLC-PRF-5 cells treated as indicated. *n*=6, biological replicates. Statistics were calculated on biological replicates with two-tailed unpaired *t*-tests, \*\**P* < 0.01. Error bars show mean with SD. **(D and E)** Fluorescent gelatin degradation and phalloidin/DAPI staining of PLC-PRF-5 cells treated as indicated (**D**) and quantification of degradation area (**E**). Scale bar, 40 µm. *n*=6, biological replicates. Statistics were calculated on biological replicates with two-tailed unpaired *t*-tests, \*\**P* < 0.01. Error bars show mean with SD. **(F)** The levels of EMT related proteins were detected by western blot in PLC-PRF-5 cells treated as indicated. **(G and H)** Representative *in vivo* images (**G**) of H22-luc-tumor-bearing-BALB/c mice with indicated treatments and liver weight/body weight ratio was quantified (**H**). *n*=6, biological replicates. Statistics were calculated on biological replicates with two-tailed unpaired *t*-tests, \*\**P* < 0.01. Error bars show median, upper and lower quartiles. **(I)** Kaplan–Meier curves showing percentage of survival of H22-luc-tumor-bearing-BALB/c mice after indicated treatments. *n*=6, biological replicates. Statistics were calculated on biological replicates with Kaplan–Meier analysis. \*\**P* < 0.01, compared with control unless otherwise indicated; #*P* < 0.05, compared with DNase1L3. **(J)** Quantification of FISH for liver tissue in H22-luc-tumor-bearing-BALB/c mice with indicated treatments. Average fluorescence intensity in the nucleus in 18 cells of FISH images from 6 biologically independent samples were measured. Statistics were calculated on biological replicates with Wilcoxon rank-sum test, \*\*\**P* < 0.01, compared with control; ##*P* < 0.01, ###*P* < 0.001, compared with DNase1L3. Error bars show median, upper and lower quartiles. **(K)** Quantification of pathological score of Immunohistochemical staining of BRAF, KRAS, CENPF in liver tissues of H22-luc-tumor-bearing-BALB/c mice. *n*=6, biological replicates. Statistics were calculated on biological replicates with two-tailed unpaired *t*-tests, \*\**P* < 0.01, compared with control; ##*P* < 0.01, compared with DNase1L3. Error bars show median, upper and lower quartiles.

Based on these findings, we further studied the pivotal role of DNase1L3 and GC on the anti-tumor effect of 25OHD. The results of SEM and E-cadherin/Vimentin IF staining showed an increase in adhesion of HCC cells and an enhancement in E-cadherin expression after 25OHD treatment, whereas the function which 25OHD exerted disappeared after knocking out of DNase1L3 or GC in HCC cells (Figure S8A). Wound healing assay (Figure S8B), Transwell assay Figure S8C), and fluorescent gelatin degradation assay (Figure S8D, E) showed that 25OHD could inhibit the migration and invasion ability of HCC cells, but the knockout of DNase1L3 or GC did not show an obvious effect. Animal studies of orthotopic liver xenograft tumors revealed that knock out of DNase1L3 or GC dramatically increase the growth of H22 xenograft tumor (Figure S8F, G) and shorten the survival of tumor-baring mice (Figure S8H). Collectively, these findings suggested the anti-tumor effect of 25OHD depend on DNase1L3 and GC.

### DNase1L3-related mRNA drugs have potential therapeutic effects on the clinical treatment of HCC

mRNA drugs have emerged as a therapeutic strategy such as COVID-19, furthermore, several studies have demonstrated the potential efficacy of employing mRNA drugs for the targeted delivery of tumor suppressor genes, thus presenting a promising avenue for cancer treatment^32^. Given that the DNase1L3 we found has the nature of a tumor suppressor gene, TCGA data (Table. S3) was utilized to perform pathological analysis of DNase1L3 expression in patients with HCC. The expression of DNase1L3 in HCC was extremely low compared with that in adjacent normal tissues (Figure 7A). HCC patients high DNase1L3 expression had a longer overall survival and disease-free survival (Figure 7B). The higher the pathological grade and clinical stage, the lower the expression of DNase1L3 (Figure 7C). The above results indicated that DNase1L3 demonstrated pathological significance in HCC, thus prompting our consideration of developing DNase1L3 as an mRNA-based therapeutic agent and evaluating its efficacy through the utilization of patient-derived xenograft (PDX) models (Figure 7D and Table S4). We also evaluated the efficacy of GC-DNase mRNA drugs. The successful of mRNA delivery was verified in Figure S9A. The PDX models showed that both DNase1L3 and GC-DNase mRNAs were able to inhibit tumor growth (Figure 7E) and prolonged survival in mice (Figure 7F). FISH was performed on the pathological slices of the tumor site, and the ecDNA level was evaluated. As shown in Figure 7G and H, after the treatment of the two mRNAs, the ecDNA levels of *BRAF*, *KRAS*, and *CENPF* were significantly reduced. IHC staining results showed that the expression of malignantly associated proteins including BRAF, KRAS, and CENPF, and Vimentin decreased, whereas the expression of E-cadherin increased (Figure 7I, J).

**Figure 7.**
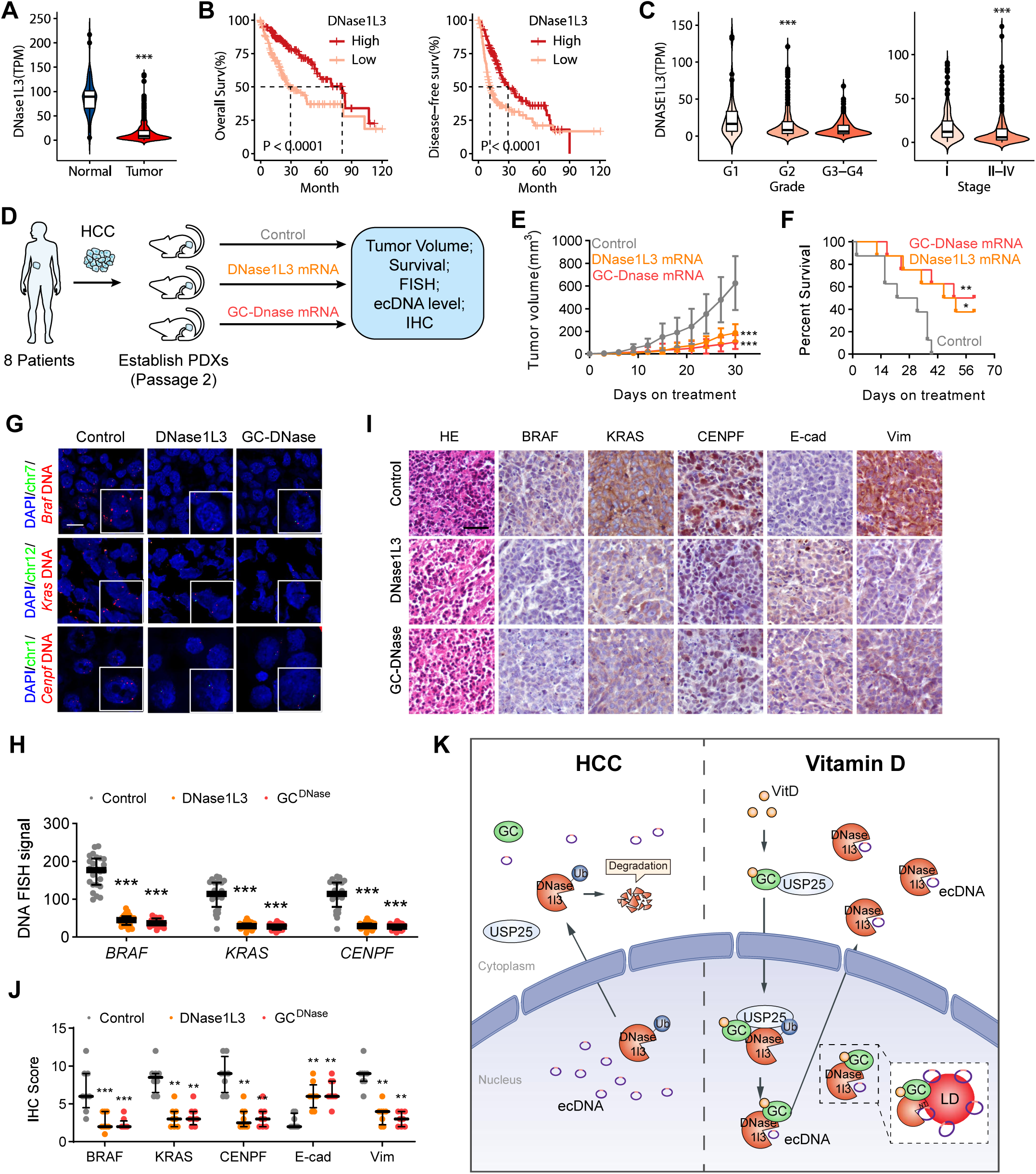
DNase1L3-related mRNA drugs have the potential therapeutic effects on the clinical treatment of HCC. **(A)** mRNA expression level (TPM) of DNase1L3 in normal and tumor tissues of HCC in the TCGA dataset. **(B)** Kaplan–Meier curves showing percentage of the overall survival (left) and disease-free survival (right) of the higher and lower expression of DNase1L3 in the TCGA dataset. **(C)** DNase1L3 mRNA expression level (TPM) in different Grade (left) and Stage (right) in the TCGA dataset. One-way ANOVA and Tukey’s multiple comparisons test was used. **(D)** Schematic of the experimental workflow of PDX. Control, injected intravenously with LNP, DNase1L3, injected intravenously with LNP containing 3 μg DNase1L3 mRNA, GC-DNase mRNA, injected intravenously with LNP containing 3 μg GC-DNase mRNA, GC-DNase mRNA consists of the enzymatically active region of DNaseL3 to the N-terminal of CDS region of GC. **(E)** Tumor growth of mice was measured starting 2 months after established of PDX mouse model with indicated mRNA treatments. *n*=8, biological replicates. Statistics were calculated on biological replicates with two-tailed unpaired *t*-tests, \*\*\**P* < 0.001, compared with control unless otherwise indicated. Error bars show mean with SD. **(F)** Kaplan–Meier curves showing percentage of survival of mice after established of PDX mouse model with indicated mRNA treatments. *n*=8, biological replicates. Statistics were calculated on biological replicates with Kaplan–Meier analysis. \**P*<0.05, \*\**P* < 0.01, compared with control unless otherwise indicated. **(G and H)** Representative images of DNA FISH for tumors of PDX model after indicated treatments (**G**) and the FISH signals were quantified (**H**). Scale bar: 10 μm. Average fluorescence intensity in the nucleus in 24 cells of FISH images from 8 biologically independent samples were measured. Statistics were calculated on biological replicates with Wilcoxon rank-sum test, \*\*\**P* < 0.001, compared with control. Error bars show median, upper and lower quartiles. **(I and J)** Representative images(**I**) of HE staining and Immunohistochemical staining of BRAF, KRAS, CENPF, E-cadherin and vimentin for tumors of PDX mouse model and pathological scores were quantified (**J**). Scale bar: 40 μm. *n*=8, biological replicates. Statistics were calculated on biological replicates with two-tailed unpaired *t*-tests, \*\**P*<0.01, \*\*\**P* < 0.001, compared with control. Error bars show median, upper and lower quartiles. **(K)** Proposed model for the role of VD, GC, DNase1L3 in HCC.

Considering that DNase1L3 could potentially function as a tumor suppressor gene, we evaluated the effect of DNase1L3 deletion on liver organoid formation using human induced pluripotent stem cells (iPSCs)-derived liver organoids. After knocking out of DNase1L3 using lentivirus (the effectiveness of knockout was validated in Figure S10F) three days before hepatic organoid maturation, the morphology of the liver organoid gradually became disordered, the normal hepatic organoids (Figure S9B) could not be formed, the liver-specific marker ALB was no longer expressed, and the liver-specific antibody hepatocyte stain was also negative (Figure S9C). Furthermore, on the basis of the DEN-induced HCC model, we found more tumors in the transfection of AAV8-encapsulated ko DNase1L3 plasmid group, as well as the significantly increase in liver size in mice (the effectiveness of knockout was validated in Extended Data Figure 10G, H) compared to the control group (Figure S9D). Thus, DNase1L3 may be a gene needed for liver development and critical for maintaining a normal state of liver, while the deletion of DNase1L3 may lead to carcinogenesis.

## Discussion

In recent years, there has been a significant increase in research findings pertaining to ecDNA, which have significantly contributed to our comprehension of cancer, DNA, and gene expression. The literature has extensively covered the distribution and content, structure and function of ecDNA, yet there remains a dearth of information regarding the degradation mechanism of ecDNA and drug interventions. In this study, we found that DNase1L3 was a crucial protein for degrading ecDNA. Furthermore, we substantiated the pivotal role of DNase1L3 across various biological systems, including cellular, animal, organoid, and PDX models. The administration of DNase1L3 as a supplementary treatment holds promise as a potential therapeutic approach against tumors. DNase1L3 inhibits proliferation, invasion, and metastasis of HCC^33^. Additionally, this study revealed a robust correlation between DNase1L3 and the clinicopathological characteristics of HCC, a prognostic factor with favorable outcomes. This study elucidated the underlying mechanism through which DNase1L3 inhibits the progression of tumors by degrading ecDNA. Previous studies have demonstrated that the deficiency of DNase1L3 predisposes individuals to systemic lupus erythematosus (SLE)^34^. Hence, our study provided insights into the therapeutic approaches for SLE.

It has been reported that the enzymatic activity of DNase1L3 could be inhibited by PARylation in DNA degradation ^30^. Under normal conditions, a large amount of PARylation writing enzyme PARP1 is present in cells, resulting in a PARylation state of DNase1L3, consequently, DNase1L3 is inhibited, thereby impeding its ability to cleave DNA and precluding the initiation of apoptosis. In the presence of inflammatory factors or other apoptosis inducers, PARP1 undergoes cleavage by Caspase-3, resulting in its inactivation. This process represents the classical apoptotic pathway. Consequently, the inactivation of PARP1 leads to the loss of PARylation state in DNase1L3, thereby activating DNase1L3 to cleave DNA, ultimately leading to the formation of a DNA ladder. In this work, we found that almost no PARylation of DNase1L3 occurred in LDs, and LDs can protect DNase1L3 from being modified by PARP1, which may be the key for DNase1L3 to degrade ecDNA around LDs but not genomic DNA under specific spatial and temporal conditions. VD/GC promoted DNase1L3 enrichment in LDs and probably also protected DNase1L3 from PARylation by PARP1 through spatial site blocking, allowing DNase1L3 in LDs to remain active and more susceptible to the degradation of ecDNA.

Our findings indicated that the lipophilic region of DNase1L3 significantly contributes to the affinity of ecDNA, implying that the protection provided by LDs could be a key factor in the degradation of ecDNA. LDs have received widespread attention from the biological community and have been recognized as intact organelles with key functions in lipid and energy homeostasis^35^. However, research on LDs in the nucleus is limited. The presence of LDs in the nucleus and their biogenesis processes have been observed and studied in the literature^36–38^, but the function of LDs in the nucleus has not been studied in depth. Nuclear LDs have been found to regulate the synthesis of phosphatidylcholine^39^. Our results suggested that intranuclear LDs may be closely related to the behavior of DNA. Specifically, in this study, we found that VD mediated GC increased the binding of DNase1L3 to LDs by interacting with DNase1L3, potentially through a liver-specific pathway. Additionally, there may be other mechanisms with similar effects. Further study is required to understand the association between intranuclear LDs and ecDNA, as well as the potential involvement of LDs in transcriptional regulation.

VD has been reported to reduce the risk of cancer mortality, and low 25OHD serum levels are associated with an increased risk of cancer in humankind^40, 41^. Several Meta-analyses have shown that cancer patients with VD deficiency (defined as 25(OH)D < 30 nmol/L)^42^ significantly improve their prognosis with dietary supplementation of VD3^43–46^. However, the mechanism underlying the anti-tumor effect of VD is not well studied, with the biological function of VD mainly mediated by its receptor VDR^9^. The VD-VDR complex binds to the VD response elements (VDREs) on the promoters of target genes and regulates their expression^47^. VD inhibits the adhesion and proliferation of cancer cells by suppressing THBS1 via VDR/Smad3 competition^48^. We demonstrated experimentally that 25OHD can physically enhance the interaction of DNase1L3 with GC and demonstrated the triple action of GC on DNase1L3. (1) Quantity: GC enters the nucleus and interacts with DNase1L3 under the activation of 25OHD, recruiting USP25, deubiquitinating DNase1L3, inhibiting DNase1L3 degradation, and ensuring the protein level of DNase1L3. (2) Spatial: 25OHD enhances the hydrophobicity of the GC/DNase1L3 interaction structural domain, making it more susceptible to lipid droplet adsorption, drawing it closer to ecDNA, and becoming more easily degraded ecDNA (spatial). (3) Activity: GC makes DNase1L3 less susceptible to PARylation and activation by promoting LDs adsorption and spatial site blocking. 25OHD/GC enhances the degradation of ecDNA by affecting three aspects: quantity, spatial distribution, and DNase1L3 activity. Only 25OHD, a form of VD that can bind to GC can degrade ecDNA, which exhibits a strict GC-dependent feature. Conversely, 1,25(OH)_2_D cannot bind GC, and experiments have shown that it cannot reduce ecDNA. Therefore, the effect of VD on ecDNA, which is not significantly associated with its involvement in calcium metabolism as a vitamin, is a novel ligand-like regulatory molecular mechanism. Given that ecDNA can explain the heterogeneity of tumors, this mechanism can also explain why VD can play a distinguished role of prevention and treatment among many cancer types. VD can promote the degradation of ecDNA, and this role may also explain the clinical conclusion that VD can reduce the incidence of autoimmune diseases^49^. Compared with DNase1L3 and GC-DNase, the effect of VD degradation of ecDNA, although indirect, is more economical and more available, and appropriate supplementation of VD in patients with cancer still has strong clinical significance. Direct vitamin D3 supplementation (800-4000 IU/day) can maintain normal 25(OH)D serum levels in humans^50^. Based on this, in our *in vivo* experiments, the anti-tumor effects of VD were verified by human-mouse dose conversion of VD3, and mice were given to intake the corresponding VD3 supplementation dose. Meanwhile, compared with the direct application of VD, our *in vivo* study showed that mRNA drugs have stronger therapeutic effects on tumors than VD, which may be related to their better targeting of ecDNA, and thus we believe that the developed mRNA therapeutics are more suitable for tumor treatment than VD. Chemotherapy is thought to lead to the accumulation of ecDNA, and this study provided a theoretical basis for the combination therapy strategies. Whether VD can be combined with targeted drugs and immunotherapy in the future also needs to be studied, and this study is believed to provide theoretical reference for future related therapies (Figure 7K).

### Limitations of the Study

Frankly speaking, there are some limitations to this work. Firstly, due to constraints in the research methodologies pertaining to ecDNA, interventions targeting individual ecDNA were limited to junction sequence deletions when assessing their impact on anti-tumor functions. Consequently, the assessment of the overall or collective anti-tumor functions of multiple ecDNA entities remains challenging. Secondly, this study did not delve deeply into exploring the other characteristics and functionalities of nuclear microLDs. Thirdly, it remains unknown whether DNase1L3-mediated degradation of ecDNA triggers DNA double-strand breaks, and whether ecDNA re-formation occurs post-degradation. Fourthly, significant variations exist in the levels of GC, DNase1L3, and lipid droplet numbers across different types of tumors.While this study focused on hepatocellular carcinoma, further investigations are warranted to determine whether VD functions via the GC pathway in other tumor types. In tumors lacking GC, DNase1L3 mRNA-based therapeutics may potentially serve as a treatment modality. Fifthly, we revealed two mechanisms by which 25OHD decrease ecDNA level through DNase1L3, one is to maintain the expression level of DNase1L3 by recruiting USP25 through GC, and the other is to protect DNase1L3 from PARylation and thus maintain its enzyme activity by increasing the size of LDs. However, the simultaneous presence of USP25 in LD, as well as the correlation between ubiquitination and PARylation of DNase1L3 needs to be further verified. Moreover, the spatiotemporal relationship between these two regulatory mechanisms in the cell deserves to be further studied. Finally, while the elucidated mechanism of ecDNA clearance in this work holds potential implications for tumor drug resistance, immunotherapy, and developmental biology, these aspects were not addressed in this manuscript and warrant further investigation.

## Acknowledgments

We appreciate the guidance and inspiration provided by Prof. Xue Yang for this study. We are grateful to Prof. Vineet Bafna, Dr. Jens Luebeck and Dr. Yue Ren for valuable guidance.

## Author Contributions

Heng Zhang conceived experiments, carried out experiments, analyzed data, drafted the manuscript and edited figures. Lu-ning Qin carried out experiments, analyzed data, drafted the manuscript, edited figures and secured funding. Qing-qing Li, Ting Wu, Lei Zhang, Kai-wen Wang, Shan-bin Cheng, Jing-xia Han and Yi-qian Feng carried out experiments and analyzed data. Yi-nan Li, Zhi-yang Li and Hui-juan Liu provided technical support. Tao Sun conceived experiments, edited the manuscript, and secured funding. All authors had final approval of the submitted and published versions of the manuscript.

## Data and materials availability

All data are available in the main text or the supplementary materials.

