## Supplementary material for "Vitamin D promotes DNase1L3 to degrade ecDNA and inhibit the malignant progression of hepatocellular carcinoma": Methods and Suplemental figures

### Supplementary Information

### Methods

#### KEY RESOURCES TABLE

| REAGENT or RESOURCE | SOURCE | IDENTIFIER |
|---|---|---|
| Antibodies | | |
| Mouse monoclonal anti-Vimentin | Proteintech | Cat# 60330-1-Ig; RRID: AB_2881439 |
| Mouse monoclonal anti-ZEB1 | Proteintech | Cat#66279-1-Ig; RRID: AB_2881662 |
| Rabbit polyclonal anti-Twist1 | Proteintech | Cat# 25465-1-AP; RRID: AB_2880093 |
| Rat monoclonal anti-CD324 (E-Cadherin) | Invitrogen | Cat# 53-3249-80; RRID: AB_10671270 |
| Rabbit polyclonal anti-beta Actin | Affinity | Cat# AF7018; RRID: AB_2839420 |
| Mouse monoclonal anti-Vitamin D binding protein (DBP) | Proteintech | Cat# 66175-1-Ig; RRID: AB_2881570 |
| Mouse monoclonal anti-DNaseI3 | Proteintech | Cat# 67041-1-Ig; RRID: AB_2882355 |
| Rabbit polyclonal anti-Histone-H3 | Proteintech | Cat# 17168-1-AP; RRID: AB_2716755 |
| Rabbit polyclonal anti-ubiquitin | Proteintech | Cat# 10201-2-AP; RRID: AB_671515 |
| Rabbit polyclonal anti-USP25 | Proteintech | Cat# 12199-1-AP; RRID: AB_2212771 |
| Rabbit polyclonal anti-DYKDDDDK tag | Proteintech | Cat# 20543-1-AP; RRID: AB_11232216 |
| Rabbit polyclonal anti-BRAF | Proteintech | Cat# 20899-1-AP; RRID: AB_2882047 |
| Rabbit polyclonal anti-KRAS | Proteintech | Cat# 12063-1-AP |
| Rabbit polyclonal anti-CENPF | Proteintech | Cat# 28568-1-AP; RRID: AB_2918176 |
| Rabbit monoclonal anti-PARP1 | ZENBIO | Cat# R25279; |
| Mouse monoclonal anti-ADFP | ZENBIO | Cat# 220889; |
| Mouse monoclonal anti-PADPr | Santa Cruz | Cat# sc-56198; RRID: AB_785249 |
| YF 633-Phalloidin | US Everbright | Cat# YP0053S; |
| YF 647 Goat Anti-Rabbit IgG (H&L) | US Everbright | Cat# Y6109L; |
| CoraLite 488-conjugated Goat Anti-Mouse IgG(H+L) | Proteintech | Cat# SA00013-1; RRID: AB_2810983 |
| Goat Anti-Rabbit IgG (H+L) HRP | Affinity | Cat# S0001; RRID: AB_2839429 |
| Goat Anti-Mouse IgG (H+L) HRP | Affinity | Cat# S0002; RRID: AB_2839430 |
| Biological samples | | |
| HCC samples for patient-derived xenografts | Tianjin Medical University Cancer Institute and Hospital | N/A |
| Chemicals, peptides, and recombinant proteins | | |

| | | |
|---|---|---|
| Cycloheximide (CHX) | Solomen | CAT# IN0194;<br>CAS: 66-81-9 |
| Forskolin | Meilunbio | CAT# MB5959;<br>CAS: 66575-29-9 |
| MG132 | Med Chem Express | CAT# HY-13259C<br>CAS: 1211877-36-9 |
| Oleic acid | HEOWNS | CAT# 0443469;<br>CAS: 112-80-1 |
| 25-hydroxy Vitamin D3 | Med Chem Express | CAT# HY-32351;<br>CAS: 19356-17-3 |
| 1,25-Dihydroxyvitamin D3 | Med Chem Express | CAT# HY-10002;<br>CAS: 32222-06-3 |
| VD3 | Med Chem Express | CAT# HY-15398<br>CAS: 67-97-0 |
| Colchicine | MERYER | CAT# M22777;<br>CAS: 64-86-8 |
| PEI 25K | Polysciences | CAT# 23966100;<br>CAS: 9002-98-6 |
| DYKDDDDK Peptide | GenScript | CAT# RP10586 |
| Plasmid-Safe ATP-Dependent DNase | Biosearch Technologies | CAT# E3101K |
| EnGen Lba Cas12a (Cpf1) | New England Biolabs | CAT# M0653S |
| S1 Nuclease | ThermoFisher | CAT# 18001-016 |
| SYBR Gold nucleic acid gel stain | ThermoFisher | CAT# S11494 |
| GeneRuler 1kb Plus DNA Ladder | ThermoFisher | CAT# SM1331 |
| Color-enhanced prestained protein molecular weight markers (10-245kD) | SparkJade | CAT# EC0020 |
| BODIPY 493/503 | GlpBio | CAT# GC42959;<br>CAS: 121207-31-6 |
| BODIPY 558/568 | Cayman Chemical | CAT# 27014<br>CAS: 158757-84-7 |
| Salidroside | Meilunbio | CAT# MB5843-1;<br>CAS: 10338-51-9 |
| Artemisinin | TCI chemicals | CAT# A2118;<br>CAS: 63968-64-9 |
| Myricetin | Med Chem Express | CAT# HY-15097;<br>CAS: 529-44-2 |
| Hesperetin | PUSH BIOTechnology | CAT# PS0243;<br>CAS: 520-33-2 |
| Baicalein | PUSH BIOTechnology | CAT# PS0711<br>CAS: 491-67-8 |
| Puerarin | PUSH BIOTechnology | CAT# PS0180;<br>CAS: 3681-99-0 |
| Vitamin C | PUSH BIOTechnology | CAT# PS2289<br>CAS: 50-81-7 |
| Lentinan | Aladdin | CAT# L413225;<br>CAS: 37339-90-5 |
| Sodium butyrate | Aladdin | CAT# S102956<br>CAS: 156-54-7 |
| Metformin | MACKLIN | CAT# N886063;<br>CAS: 657-24-9 |
| **Critical commercial assays** | | |
| Cell Counting Kit-8 (CCK-8) | APExBIO | CAT# K1018 |

| *Continued* | | |
|---|---|---|
| Cell Membrane Protein and Cytoplasmic Protein Extraction Kit | Beyotime | CAT#P0033 |
| DNA Native PAGE Electrophoresis Kit | RealTimes | CAT# RTE4101 |
| One-4-All Genomic DNA Miniprep Kit | BIO BASIC | CAT# B618503 |
| Lipid Droplet Isolation Kit | CELL BIOLABS | CAT# MET-5011 |
| Cell Specimen FISH Pretreatment Kit | Exon Biological | CAT# D-0014 |
| Paraffin Pretreatment Kit II | Exon Biological | CAT# D-0002 |
| Fast Silver Stain Kit | Beyotime | CAT# P0017S |
| Duolink In Situ Detection Reagents Orange | Sigma Aldrich | CAT# DUO92007 |
| Duolink In Situ PLA Probe Anti-Rabbit MINUS | Sigma Aldrich | CAT# DUO92005 |
| Duolink In Situ PLA Probe Anti-Mouse PLUS | Sigma Aldrich | CAT# DUO92001 |
| MonoRab Anti-DYKDDDDK Magnetic Beads | GenScript | CAT# L00835 |
| MagAttract HMW DNA Kit | QIAGEN | CAT# 67563 |
| Experimental models: Cell lines | | |
| Hep3B | CELLCOOK | CAT#CC0103 |
| SK-Hep1 | Keygen Biotech | CAT#KG064 |
| PLC-PRF-5 | Keygen Biotech | CAT#KG068 |
| H22-LUC | ZQXZbio | CAT# LZQ0029 |
| MHCC-97L | ZQXZbio | CAT# ZQ0019 |
| MHCC-97H | ZQXZbio | CAT# ZQ0020 |
| Experimental models: Organisms/strains | | |
| BALB/c mice | Beijing Vital River Laboratory Animal Technology | |
| C57BL/6 mice | Beijing Vital River Laboratory Animal Technology | |
| BALB/c nude mice | Beijing Vital River Laboratory Animal Technology | |
| Oligonucleotides | | |
| Primers for DNase1L3<br>Forward: TGCAGCTACGTCCCCAAGA<br>Reverse: TCCCCGATCAGCCAAACAAAC | TsingkeBiotechnology | |
| Primers for GAPDH<br>Forward: ACAACTTTGGTATCGTGGAAGG<br>Reverse: GCCATCACGCCACAGTTTC | TsingkeBiotechnology | |
| Primers for BRAF mRNA<br>Forward: AATACACCAGCAAGCTAGATGC<br>Reverse: AATCAGTTCCGTTCCCCAGAG | TsingkeBiotechnology | |
| Primers for KRAS mRNA<br>Forward: ACAGAGAGTGGAGGATGCTTT<br>Reverse: TTTCACACAGCCAGGAGTCTT | TsingkeBiotechnology | |
| Primers for CENPF mRNA<br>Forward: GCCATTTTGAGCAGTGTGATG<br>Reverse: TCAGGTTCAGAGTGTCAGCC | TsingkeBiotechnology | |
| Primers for Mitochondrial DNA<br>forward: GCCCACTTCCACTATGTCCT<br>Reverse: GATTTTGGCGTAGGTTTGGTCT | TsingkeBiotechnology | |

| *Continued* | | | |
|---|---|---|---|
| Primers for BRAF ecDNA<br>forward: TTTTCTGTCTTCTTCCTCTGTGT<br>Reverse: TCAATGTATGTTCTTGGCACCT | TsingkeBiotechnology | | |
| Primers for KRAS ecDNA<br>forward: AAGAGTTTGGAGTTGGAATTGC<br>Reverse: GCCTCTGCCTACTTGTCTCC | TsingkeBiotechnology | | |
| Primers for CENPF ecDNA<br>forward: ATTTTTCCACAGACCAATGGC<br>Reverse: GCAAGTATTTTCCTCCAGGTT | TsingkeBiotechnology | | |
| Recombinant DNA | | | |
| GC_OHu14110D_pcDNA3.1+/C-(K)-DYK | Genscript | | |
| DNASE1L3_OHu20141D_pcDNA3.1+/C-(K)-DYK | Genscript | | |
| DNase1L3$^{\Delta NT}$-mEGFP | TsingkeBiotechnology | | |
| DNase1L3$^{3\times NT}$-EGFP | TsingkeBiotechnology | | |
| DNase1L3-EGFP | TsingkeBiotechnology | | |
| GC-DNase-mEGFP | TsingkeBiotechnology | | |
| GC-EGFP | TsingkeBiotechnology | | |
| NLS-GC_pcDNA3.1+/C-(K)-DYK | TsingkeBiotechnology | | |
| NES-GC_pcDNA3.1+/C-(K)-DYK | TsingkeBiotechnology | | |
| CRISPR/Cas9-based knock out | | Targeting Sequence (5'-3') | |
| sgRNA: Ko VDR_eSpCas9-2A-Puro (PX459) V2.0 | Genscript | ACTTTGACCGGAACGTGCCC | |
| sgRNA: ko GC_eSpCas9-2A-Puro (PX459) V2.0 | Genscript | ACCCTGACTGCTATGACACC | |
| sgRNA: DNase1L3 ko_eSpCas9-2A-Puro (PX459) V2.0 | Genscript | CTGTGACATCATACTCGTGA | |
| sgRNA: Ko USP25_eSpCas9-2A-Puro (PX459) V2.0 | Genscript | GTTGATTCAAAAACGTCTGC | |
| sgRNA: KRAS ko_eSpCas9-2A-Puro (PX459) V2.0 | Genscript | AAATATAGATGAAGGTACTA | |
| sgRNA: BRAF ko_eSpCas9-2A-Puro (PX459) V2.0 | Genscript | TTCTTGGCACCTTCATAATG | |
| sgRNA: CENPF ko_eSpCas9-2A-Puro (PX459) V2.0 | Genscript | GCATTAGGCAAACAACAACCTGG | |
| DNA FISH probes Targeted Gene | Coordinates of the FISH probes | BAC ID | |
| BRAF | Chr7:140503484-140699716 | RP11-948O19 | |
| KRAS | Chr12:25230469-25426524 | RP11-295I5 | |
| CENPF | Chr1:214745662-214921681 | RP11-262H5 | |
| Software and algorithms | | | |
| ZEN | Zeiss | | https://www.zeiss.com/microscopy/en/products/software/zeiss-zen.html |
| Ecseg | https://doi.org/10.1016/j.isci.2019.10.035. | | https://github.com/UCRajkumar/ecSeg |

| *Continued* | | | |
|---|---|---|---|
| Circle-Map | https://doi.org/10.1186/s12859-019-3160-3 | | https://github.com/iprada/Circle-Map |
| IGV | IGV | | https://igv.org/ |
| R 4.1.3 | R Foundation | RRID:SCR_001905 | https://www.r-project.org/ |
| ImageJ | Open source processing software | RRID:SCR_003070 | https://imagej.nih.gov/ij/ |
| GraphPad Prism 9.0.0 | GraphPad | RRID:SCR_002798 | https://www.graphpad.com/scientific-software/prism/ |
| FlowJo 10.6.2 Software | FlowJo LLC | RRID:SCR_008520 | https://www.flowjo.com/ |
| Pymol | Schrödinger, LLC | RRID:SCR_000305 | https://pymol.org/2/ |
| GROMACS | GROMACS | RRID:SCR_014565 | https://www.gromacs.org/ |

| Other | |
|---|---|
| Zeiss LSM800 with Airyscan | Zeiss |
| Zeiss Elyra7 | Zeiss |

| mRNA Drugs | | |
|---|---|---|
| Name | Vector | DNA template |
| DNase1L3 | pUC57-mini-Kana-BsmBI free-terminator-T7 deleted-100A-BspQI | AAGCTTTAATACGACTCACTATAAGGGGCTAGCGATCTTCTGGTCCCCACA GACTCAGAGAGAACCCGCCACCATGTCTCGTGAGCTGGCCCCTTTGCTACT CCTGCTGTTGAGCATCCATTCTGCATTGGCCATGCGGATCTGTTCCTTTAAT GTGCGTTCATTTGGAGAGTCTAAGCAGGAAGACAAAAATGCAATGGATGTC ATTGTCAAAGTCATTAAAAGATGTGACATAATTCTTGTGATGGAGATCAAGG ACTCTAACAACAGGATATGCCCCATTTTAATGGAAAAGCTCAATAGAAACTC AAGACGAGGCATCACCTACAACTACGTGATTAGTTCACGGCTCGGGAGAAA TACCTATAAGGAGCAGTATGCTTTCCTGTATAAAGAGAAACTGGTGTCTGTG AAACGGTCCTACCACTACCATGACTACCAGGATGGAGATGCTGATGTATTC TCCCGAGAACCCTTCGTGGTCTGGTTCCAGTCTCCTCACACAGCTGTGAAG GATTTTGTCATCATCCCGCTGCACACGACACCTGAAACCAGCGTTAAAGAA ATTGATGAGTTAGTGGAGGTCTACACAGATGTGAAGCACCGCTGGAAAGCA GAGAACTTCATCTTCATGGGTGACTTCAACGCAGGCTGCAGCTATGTTCCC AAGAAAGCCTGGAAGAACATCCGGCTCAGAACTGACCCAAGGTTTGTGTGG CTGATAGGGGACCAAGAAGACACTACTGTTAAAAAGTCCACGAACTGCGCC TATGATCGCATCGTACTACGCGGCCAGGAGATTGTTAGCAGCGTGGTGCCA AAAAGCAATAGTGTGTTTGACTTTCAGAAGGCCTACAAGCTGACCGAAGAG GAAGCTCTGGATGTATCGGACCATTTCCCAGTAGAGTTTAAGCTGCAATCC TCTAGGGCGTTCACAAATAGCAAGAAGTCAGTCACCCTTAGGAAGAAGACC AAGAGTAAAAGGTCCTGATAATAGGCTGGAGCCTCGGTGGCCATGCTTCTT GCCCCTTGGGCCTCCCCCCAGCCCCTCCTCCCCTTCCTGCACCCGTACCC CCGTGGTCTTTGAATAAAGTCTGAGTGGGCGGCAGGATCCCGTCTCTTAAC TAACTAAACTAGTAAAAAAAAAAAAAAAAAAAAAAAAAAAAAAAAAAAAAAAA AAAAAAAAAAAAAAAAAAAAAAAAAAAAAAAAAAAAAAAAAAAAAAAAAAAAA AAAAAAA |

| | | |
|---|---|---|
| GC-DNase | pUC57-mini-Kana-BsmBI free-terminator-T7 deleted-100A-BspQI | AAGCTTTAATACGACTCACTATAAGGGGCTAGCGATCTTCTGGTCCCCACA GACTCAGAGAGAACCCGCCACCATGAAGAGAGTGTTGGTGTTACTGTTGGC AGTGGCATTTGGACACGCCCTGGAGCGAGGCAGAGATTACGAGAAGAACA AAGTATGTAAAGAGTTTAGTCACCTTGGGAAGGAAGACTTTACCAGCCTCA GCCTGGTTCTGTACAGCAGGAAGTTCCCCAGTGGGACGTTTGAACAAGTGT CCCAACTGGTCAAGGAGGTGGTTTCTCTGACAGAAGCCTGTTGTGCCGAAG GCGCCGACCCGGACTGTTATGACACGAGAACTTCAGCCCTGTCCGCAAAAT CCTGCGAGAGCAATTCCCCCTTTCCTGTGCATCCCGGGACCGCAGAATGTT GCACCAAAGAAGGGCTCGAACGCAAATTATGTATGGCAGCTCTAAAACACC AGCCTCAGGAATTCCCGACATATGTCGAGCCCACCAATGACGAGATCTGTG AGGCCTTCCGGAAAGACCCTAAGGAGTATGCCAACCAGTTCATGTGGGAGT ACTCCACTAACTATGGGCAGGCTCCGCTCTCACTTCTGGTATCATACACGA AGAGTTACCTCTCCATGGTCGGGAGCTGCTGCACAAGCGCCAGCCCTACC GTTTGTTTTCTTAAAGAAAGACTGCAGCTCAAACATCTCTCCCTCCTGACTA CTCTGAGTAACCGGGTGTGCTCCCAGTATGCAGCGTATGGAGAAAAGAAGT CTAGGCTTTCCAATCTGATCAAACTAGCTCAAAAGGTTCCCACGGCTGACC TCGAAGATGTCCTTCCATTGGCAGAAGACATCACAAACATTCTCTCTAAGTG TTGTGAGTCCGCGAGTGAAGATTGCATGGCCAAAGAATTACCAGAGCACAC CGTGAAACTATGCGACAACCTCTCTACCAAAAATTCGAAATTCGAGGACTG CTGCCAGGAGAAAACTGCGATGGACGTATTTGTCTGCACATATTTTATGCCT GCTGCCCAGCTGCCCGAACTCCCAGACGTGGAACTTCCAACTAACAAGGA CGTCTGTGACCCAGGTAACACTAAGGTCATGGATAAGTACACCTTCGAGCT CAGCAGGAGGACTCATCTCCCAGAAGTTTTCCTTAGCAAGGTATTGGAGCC TACGCTAAAATCACTGGGCGAATGCTGTGATGTGGAGGATAGCACAACTTG TTTCAATGCTAAGGGCCCTCTGCTCAAGAAGGAGTTGTCGAGCTTCATCGA CAAGGGACAAGAGCTGTGCGCCGACTACAGTGAGAATACTTTCACTGAGTA TAAGAAAAAGCTGGCCGAGCGATTGAAGGCTTGATAATAGGCTGGAGCCTC GGTGGCCATGCTTCTTGCCCCTTGGGCCTCCCCCCAGCCCCTCCTCCCCT TCCTGCACCCGTACCCCCGTGGTCTTTGAATAAAGTCTGAGTGGGCGGCA GGATCCCGTCTCTTAACTAACTAAACTAGTAAAAAAAAAAAAAAAAAAAAAAA AAAAAAAAAAAAAAAAAAAAAAAAAAAAAAAAAAAAAAAAAAAAAAAAAAAAA AAAAAAAAAAAAAAAAAAAAAAAA |

#### Cell culture

The human HCC cell lines Hep3B was obtained from the Guangzhou CELLCOOK, SK-Hep1, PLC-PRF-5 were purchased from Nanjing Keygen Biotech. MHCC97-L, MHCC97-H and H22-Luc were purchased from Shanghai Zhong Qiao Xin Zhou Biotechnology. SK-Hep1 and H22-Luc cells were cultured in RPMI-1640 medium (Keygen Biotech), and Hep3B, PLC-PRF-5, MHCC97-L and MHCC97-H cells were cultured in Dulbecco's modified Eagle's medium (Keygen Biotech) supplemented with 10% (v/v) fetal bovine serum (Thermo Fisher) at 37 °C in humidified atmosphere containing 5% $CO_2$. The cell lines were tested to determine the presence of Mycoplasma before use. Complete cell identification was provided by CELLCOOK or KeyGen Biotech.

#### Plasmids construction and Overexpression and CRISPR knockout

The DNase1L3$^{\Delta NT}$ sequence was obtained by deleting the N-terminal lipophilic region sequence (NT, MSRELAPLLLLLLSI) and the C-terminal DYK sequence of the DNase1L3 protein coding sequence from the DNase1L3_OHu20141D_pcDNA3.1+/C-(K)-DYK plasmid (Genscript). The obtained sequence was then inserted into the pmEGFP-N1 vector to construct DNase1L3$^{\Delta NT}$-EGFP. DNase1L3$^{3\times NT}$-EGFP was constructed by adding three lipophilic region repeats (3×NT, MSRELAPLLLLLLSIMSRELAPLLLLLLSIMSRELAPLLLLLLSI) to the N terminus of the DNase1L3 protein coding region and deleting the C-terminal DYK sequence from the DNase1L3_OHu20141D_pcDNA3.1+/C-(K)-DYK plasmid (Genscript), followed by insertion into the pmEGFP-N1 vector. The GC-DNase plasmid was obtained by deleting the sequence of the NT lipophilic region, as well as the C-terminal SSRAFTNSKKSVT and DYK tag of DNase1L3 from the DNase1L3 _OHu20141D_pcDNA3.1+/C-(K)-DYK plasmid. The obtained sequence was inserted into the GC_ OHu14110D_pcDNA3.1+/C-(K)-DYK plasmid at the C-terminus of the GC protein coding region, with insertion of linker GGGGSGGGGSGGGGS between the two sequences to achieve fusion expression of GC and DNase1L3 proteins. Information about these plasmids is also provided in the Materials table.

PLC-PRF-5 cells were transfected using Lipo8000 (Beyotime) according to the manufacturer's instructions with Flag-DNase1L3, Flag-GC, and Flag-USP25, DNase1L3-EGFP, DNase1L3$^{\Delta NT}$-EGFP, DNase1L3$^{3\times NT}$-EGFP, NLS-GC, NES-GC, Flag-PARP1, GC-DNase-EGFP overexpression plasmid or CRISPR Knockout Plasmids containing Cas9 and guide RNAs of GC, VDR, DNase1L3, and USP25. Transfected cells were incubated for 48 h before puromycin selection. GC, VDR, DNase1L3, and USP25 knockout were confirmed by western blotting (Fig. S10A-D).

**Cell proliferation assay**

At 0, 12, 24 and 48 h after treatment, cell viability was measured by Cell Counting Kit-8 (CCK8, APExBIO). Cell culture medium was removed and 10 μL of CCK8 solution was added to each well of a 96-well plate. Afterward, the cells were incubated for 1 h at 37° C in 5% $CO_2$. Absorbance was measured at a wavelength of 450 nm using an ELISA microplate reader (Thermo Fisher). Assays were repeated at least three times.

**Cell apoptosis**

After 48 hours cells were harvested with 0.25% trypsin without EDTA. They were also washed twice with ice-cold phosphate-buffered saline (PBS), resuspended in 250 mL of binding buffer, adjusted to $1\times10^6$ /mL, and fixed in 1% paraformaldehyde. Subsequently, they were stained with the Annexin V/PI Apoptosis Detection Kit (KeyGen Biotech). They were analyzed with a flow cytometer (Millipore) after incubation in the dark for 30 min.

**Colony formation assay**

Colony formation was performed in 6 cm culture dishes (Corning). Totally 600 cells were seeded per dish and cultured for 14 days. Colonies were fixed in methanol for 10 min and stained with 0.1% crystal violet for 30 min at room temperature. The number of colonies was counted by ImageJ.

**Wound healing assay**

Cells were seeded and cultured in a 6-well plate, and the cell density reached an appropriate level. Use a sterile pipette tip to evenly scrape the cells for a scratch test. Add serum-free medium and use live cell imaging for 12, 24, and 48 h respectively. The cell migration was photographed and measured, and analyzed by ImageJ.

**Transwell**

Matrigel (Corning, 354234) was diluted in DMEM at a ratio of 1:8, $1\times10^5$ cells were seeded into a 24-well Transwell chamber (JET, TCS013024) covered with matrigel and then incubated at 37 C for 48 h. 1% crystal violet was used to stain the invasive cells on the lower surface of the chamber. Images were taken by using a phase-contrast microscope.

**Gelatin degradation assay**

Prepare pork skin gelatin (Thermo Fisher, G13187) diluted with 2% sucrose in PBS. Gelatin was used at a final concentration of 0.2 mg/mL. The working solution is protected from light and heated to 60° C to cover the slides evenly, after drying, add 1 mL of a pre-chilled solution of glutaraldehyde and incubate for 15 min on ice. Wash coverslips three times at room temperature with PBS. Add 1 mL of a freshly prepared solution of Sodium Borohydride and incubate for 3 min at room temperature. Inoculate cells and culture for 24 ho Images were acquired on the ZEISS LSM 800 Inverted Confocal microscope attached with an Airyscan area detector.

**Pull down**

PLC-PRF-5 cells expressed Flag-GC were harvested in lysis buffer (50 mM Tris-HCl, pH7 .5, 100 mM NaCl, 5 mM $MgCl_2$, 0.5% Triton, protease inhibitor cocktail). Anti-Flag tag affinity beads (Beyotime) were used to incubate with the cell lysates extracts for 12 h at 4° C. After binding, wash the beads with lysis buffer (3×10 min. And then elute the Flag protein complex. The eluates were gathered and visualized by Western blotting.

**Silver staining**

Protein eluates were collected by pull-down assay. The 10% SDS-PAGE was used to separate proteins, and the electrophoresis were performed with a constant voltage (80 V for concentrated glue, 120 V for separated glue) in the running buffer until the dye front reached the lower end of the gel (2 ho. According to the manufacturer's manual, the gel was silver stained under shaking with Fast Silver Stain Kit (Beyotime).

**LC-MS/MS**

The target bands on SDS-PAGE were collected and cut into approximately 1 $mm^3$ size, decolorized by adding 200 µL of 50 mM $NH_4HCO_3$ 50% acetonitrile in a centrifuge tube and shaking for 10 min at 37 °C, until the gel becomes transparent. Add 200 µL acetonitrile and wash the gel 2 times. Subsequently dry it at 37 °C for 15 min at 300 rpm. Add 100 µL of 10 mM dithiothreitol and shake at 56 °C for 45 min at 400 rpm. After that, add 200 µL of acetonitrile to wash the gel. Discard the liquid in the tube and immediately incubate the gel by adding 100 µL of iodoacetamide at room temperature and protected from light for 30 min. Sequentially add 200 µL of 50 mM $NH_4HCO_3$/50% acetonitrile, 100% ACN to wash the gels. Dry the gel and incubate it by adding 100 µL of 10 ng/µL trypsin (Thermo Scientific, 90057) at room temperature for 1 min, followed by ice bath for 5 min. The supernatant was removed by centrifugation at 130,00 rpm for 30 s. 200 µL 25 mM $NH_4HCO_3$ was added to incubate the gel at room temperature for 1 min, followed by an ice bath for 10 min. Overnight incubation was conducted at 300 rpm at 37 °C. After completion, 200 µL of 1% formic acid 2% acetonitrile-$H_2O$ solution, 50% acetonitrile-$H_2O$ solution and 50 µL of 1% formic acid-acetonitrile solution were added sequentially to the original tube of gel, shaken thoroughly and sonicated for 10 min. The purified tryptic peptides were analyzed by LC-MS/MS retrieval and protein identification was conducted by MaxQuant software.

**Western blotting**

Cell lysates were collected, and the protein levels were quantified using a standard BCA assay kit (Thermo Fisher, 23227). Proteins were separated by 10% SDS-PAGE gel and transferred onto PVDF membranes (Millipore). After blocked in 5% skim milk for 2 ho the membranes were incubated with primary antibody at 4° C overnight, and then incubated with horseradish peroxidase-conjugated goat anti-rabbit or goat anti-mouse IgG secondary antibody (Beyotime) for 2 h at room temperature. Membranes were washed 3 times with TBST for 10 min. Protein expression was assessed using an enhanced chemiluminescent substrate (Affinity) and exposed to a chemiluminescent film.

**Co-IP assay**

For Immunoprecipitation, 50 μL of Protein A/G agarose (Beyotime, P2012) was incubated with antibodies against GC, DNase1L3, USP25 overnight at 4° C with continuous rotation. The lysates were centrifuged at 120,00 rpm for 10 min at 4° C, and then incubated with the antibody-conjugated beads overnight at 4° C. After incubation, washed the beads three times with cold lysis buffer. The precipitated proteins were separated from the beads by resuspending the beads in 1×SDS-PAGE loading buffer and boiling at 99° C for 10 min. Consequently, the boiled proteins were analyzed by Western blotting.

**Quantitative PCR**

Total RNA of HCC cells was extracted using Trizol reagent (Invitrogen, 15596026) in accordance with the manufacturer's instructions. cDNA was obtained by a FastQuant RT kit (TIANGEN, R6906). Transcript quantification was performed by a SYBR green kit (TIANGEN, FP205). The samples were run in triplicate in each experiment. The $2^{-\Delta\Delta CT}$ method was applied to quantify relative gene expression. The qRT-PCR primers are listed in the Materials table.

**Protein expression and purification**

Flag-DNase1L3, Flag-GC, plasmids were overexpressed in HEK293F cells. Cells were then grown to optical density of $1.2\times10^6$ cells/mL at 37 °C and transfected with Flag-Tagged plasmids using PEI (Polysciences, 23966-2). After 48 h of transfection, cells were harvested by centrifugation at 800g. The collected cells were resuspended lysis buffer [50 mM tris-HCl (pH 7.4), 150 mM NaCl, 1 mM EDTA, 0.5%T rittonX-100, 10% Glycerol, protease inhibitor cocktail (TargetMol)] and were lysed by sonication for 2 h at 4° C. The supernatant was collected by centrifugation at 120,000g for 60 min at 4 °C and then incubated with anti-DYKDDDDK Magnetic Beads (Genscript, L00835) for 1 h at 4° C. Wash the beads for 3 times using lysis buffer, proteins were eluted with competitive DYKDDDDK peptide (Genscript, RP10586).

**Surface plasmon resonance (SPR) assay**

SPR experiments were performed using a BIACORE T200 instrument (GE Healthcare, GER) at 25° C. DNase1L3 protein were immobilized on CM5 sensor chips using the Biacore Amine Coupling Kit in accordance with the manufacturer's instructions. Briefly, the chip was activated using a 1:1 mixture of 0.2 M N-ethyl-N'-(3-dimethylaminopropyl) carbodiimide and 0.05 M N-hydroxysuccimide at 10 μL/min for 7 min. DNase1L3 was coated on the chip at 150 μg/mL in 10 mM sodium acetate buffer (pH 5.0). GC protein was diluted to 20 μg/ml by 1×PBS (pH 7.4). Finally, a series of GC from 0–220 nM was used as kinetic analytes. 1×PBS (pH 7.4, 0.1% DMSO) was used as the running buffer for analyte dilution. All buffers were filtered and degassed prior to use and the data were analyzed using the BIA evaluation software (Version 4.1).

**Microscale thermophoresis**

Proteins were labelled with the cysteine-reactive dye Alexa Fluor 488 C5 maleimide (Thermo Fisher). Purified DNase1L3 protein was incubated with ~20-fold molar excess of the dye in gel-filtration buffer for 30 min. After labelling, the excess dye was removed by applying the sample on Superdex 200 column equilibrated with the gel filtration buffer. Tween 20 (0.05%) was added to the protein buffer for microscale thermophoresis measurements before the experiment. Datasets were collected at a temperature of 25° C. For the Alexa Fluor 488-labelled DNase1L3 interaction with GC, a concentration series of GC was prepared using a 1:1 serial dilution of GC in PBS buffer. The range of GC concentration used was from 28 μM to a final concentration of 0.85 nM, over 16 serial diluted capillaries with 10 μL samples. The interaction was initiated by the addition 10 μL of 400 nM Alexa Fluor 488-labelled DNase1L3 to each reaction mixture, resulting in a 200 nM final concentration of DNase1L3. The LED power was 100% and the microscale thermophoresis power (that is, the power supplied to the infrared laser) was 60%. The pre-microscale thermophoresis period was 5 s, the microscale thermophoresis acquisition period was 20 s, and the post- microscale thermophoresis period was 5 s. Data were analyzed by MO Control software provided by NanoTemper.

**Extraction and isolation of ecDNA**

EcDNA was firstly extracted following the published procedures(*1*). In brief, MagAttract HMW DNA Kit (Qiagen, Hilden, Germany, 67563) was applied for genomic DNA extraction. To eliminate linear chromosomal DNA and purify circular DNA, the genomic DNA was treated with Plasmid-Safe ATP-dependent DNase (Lucigen) at 37 °C, after 6 days of digestion, heat inactivate the exonuclease by incubating at 70 °C for 30 min and then extracted with PCI solution (25:24:1). To confirm elimination of linear chromosomal DNA, the sensitivity of the extracted ecDNA to Plasmid-Safe ATP-dependent DNase was conducted. For the isolation of ecDNA, we recovered the large molecular weight ecDNA by agarose gel electrophoresis using the TIANgel Medi Purification Kit (TIANGEN, cat. #DP209-02), which we subsequently stained with YOYO1 before SIM imaging, to confirm the circular structure of the extracted ecDNA and the integrity of the circular structure. The isolated ecDNAs were used for Cellular delivery.

**Cellular delivery of ecDNA**

The effect of traditional liposome transfection reagents on the delivery of DNA with a molecular weight of over 100 kb is not clear. In recent years, various types of extracellular vesicle technologies, have been widely studied and used for substance delivery including drugs and DNA using cell membrane-coated nanoparticles (CNPs)(*2*). The flexible and variable cavities of CNPs can allow the encapsulation of large molecules such as ecDNA, and their nanoproperties and their homology with the target cell membrane allow them to easily enter the cell by endocytosis. Therefore, we utilized PLC-PRF-5 cell-derived CNPs to encapsulate and deliver ecDNA.

Extraction of tumor cell membranes: The harvested PLC-PRF-5 cells were resuspended using Solution A in the Cell Membrane Protein and Cytoplasmic Protein Extraction Kit (P0033, Beyotime), and incubate at 4 °C for 15 min, and then repeatedly freeze-thawed in liquid nitrogen and 37 °C water bath, cycling three times. The suspension was centrifuged at 4 °C, 800 g, for 10 min, the supernatant was collected, and the precipitate in the tube was discarded, which consisted of nuclei and unbroken cells. The supernatant was centrifuged in a centrifuge at 4 °C, 14000 g for 30 min, the supernatant was discarded; the resulting precipitate was PLC-PRF-5 cell membrane. The ecDNAs were dissolved in PBS and were added to the PBS solution containing PLC-PRF-5 cell membrane vesicles (CMVs) in a certain proportion and incubated at 4 °C for 1 h.

Extracellular vesicles (EV) preparation: The liposome extruder and polycarbonate filter membrane were soaked in 75% ethanol for 30 min, and then irradiated with UV light on an ultra-clean bench for 30 min. The CMVs-coated ecDNA suspension was transferred into a liposome extruder and extruded from different pore sizes of polycarbonate filter membrane (0.8 μm, 0.4 μm, 0.2 μm) in ascending order, and each pore size was extruded three times. The vesicles obtained were collected and added to cultured PLC-PRF-5 cells.

**ecDNA linearization**

ecDNA linearization was performed following the reported method (*3*). ecDNA was linearized by using the nickase fnCpf1 (NEB, M0653), 50 ng of ecDNA were nicked by incubating in a 20 μL reaction that contained 10× fnCpf1 linearization buffer and 1 μL fnCpf1 at 37 °C for 1 h, Nicked ecDNAs were linearized in 10 μL reaction that include 2 μL 5× buffer, 1 μL S1 Nuclease (Thermo Fisher, 18001016) at 37 °C for 5 min. The successful linearization of ecDNA was verified by verifying its sensitivity to exonuclease ATP-dependent DNase (Lucigen). After treatment with exonuclease, the concentration decreased in a time-dependent manner, indicating that the sample was already linear.

**Assay of DNase1L3 activity for degrading ecDNA in vitro**

Proteins were diluted immediately before use to a final concentration of 10 mg/mL and various combinations and concentrations of cations. Purified ecDNA or linear DNA were co-incubation with DNase1L3 protein in a solution containing 5 mM $MgCl_2$, 2.5 mM $CaCl_2$ at 37 °C for 1 h. $ZnCl_2$ was added to the reaction to inhibit the nuclease activity of DNase1L3. When conducting the liposome containing reaction, ecDNA was pre-incubated with liposomes for 10 min at room temperature, then proteins were added, pipe gently and incubate the reaction at 37 °C for 30 min or 2 h. The DNA concentration was measured by Qubit.

**Immunofluorescence staining**

Cells were cultured on coverslips in 24-well plates, and then were washed three times with PBS, fixed with 4% paraformaldehyde for 20 min and permeabilized with blocking buffer (QuickBlock Blocking Buffer for Immunol Staining, Beyotime) for 30 min at room temperature. Subsequently, the cells were incubated with primary antibodies for 2 h at room temperature. The cells were washed with PBS (3×5 min) and incubated with fluorochrome-labeled secondary antibody for 2 h and phalloidin-594 for 30 min at room temperature. Finally, coverslips were stained with DAPI and the analysis of immunofluorescence using ZEISS LSM 800 Inverted Confocal microscope attached with an Airyscan area detector.

**BODIPY staining of lipid droplets**

Cells were co-incubated in PBS containing Hoechst and 20 μg/mL BODIPY-558/568 for 45 min at 37 °C, and then were washed three times with PBS. For fluorescence imaging, cells were visualized using live imaging by SIM. For lipid droplets in fixed samples, cells were stained with 1 µg/mL BODIPY-493/503 and DAPI for 10 min at room temperature, and then visualized using SIM.

**Scanning electron microscope (SEM)**

Soak the sample with 2.5% glutaraldehyde, overnight at 4 °C, dehydration with gradient ethanol twice, 15 min each time. Gradient tert-butanol was dried for 15 min each time, after which the sample was condensed at four degrees and dried in vacuum. Spray gold, the sample was imaged with scanning electron microscope.

**Duolink in situ proximity ligation assay (PLA)**

The Duolink in situ PLA was performed using Duolink In Situ Red Starter Kit Mouse/Rabbit (Sigma-Aldrich) according to the manufacturer's protocol. In brief, PLC-PRF-5 cells were plated on glass coverslips, rinsed three times with PBS and fixed in 4% formaldehyde in PBS for 10 min. The cells were permeabilized in 0.5% TritonX-100 for 5 minutes and blocked with 3% BSA in PBS for 60 min at 37 °C. After blocking, cells were then incubated with primary antibodies in PBS containing 1% BSA overnight at 4 °C, followed by incubation with corresponding secondary antibodies conjugated with PLA probes for 60 min at 37 °C in the dark. Cells were washed three times in wash buffer. Finally, the cells were stained with DAPI, and Duolink signals were detected using ZEISS LSM 800 Inverted Confocal microscope attached with an Airyscan area detector.

**Metaphase chromosome spread**

After 2 h treatment with 0.1 µg/mL colchicine, cells in metaphase were obtained. Cells were collected, washed with PBS, and suspended in 0.075 M KCl for 15-30 min. Carnoy's fixative (3:1, methanol: glacial acetic acid) were added to stop the reaction. The cells were washed additional 3 times with Carnoy's fixative, and then dropped on a moistened glass coverslip. DAPI was added to the slides for ecSeg analysis. Images were acquired on the ZEISS LSM 800 Inverted Confocal microscope attached with an Airyscan area detector.

**DNA FISH for cells**

FISH was performed by applying Cell specimen FISH pretreatment kit (EXONBIO, Guangzhou, D-0014). DNA FISH probes targeting *BRAF* (Chr7: 126063877-149379609), *KRAS* (Chr12: 22403560-26647340), *CENPF* (Chr1:214497072-215108050) ecDNA were ordered from EXONBIO, Guangzhou. In brief, FISH was performed by adding 10 µL DNA FISH probes (EXONBIO, Guangzhou) onto slides containing fixed cells in interphase or metaphase, followed by applying a coverslip and seal with rubber cement.

DNA denaturation was performed at 85 °C for 5 min, then hybridization in the hybridizer overnight at 42 °C. Wash the slides in 1×wash buffer for 2 min at 72 °C, followed by a final wash in 1×wash buffer at room temperature for 5 min. Metaphase cells and interphase nuclei were stained with DAPI, a coverslip was applied and images were captured on the ZEISS LSM 800 Inverted Confocal microscope attached with an Airyscan area detector.

**DNA FISH for tissues**

FISH was performed by applying Paraffin Pretreatment Kit (EXONBIO, Guangzhou, D-0002). In brief, specimens were baked at 70 °C for more than 2 h and then immersed in xylene substitute for dewaxing. Subsequently, soak the specimens in 100% alcohol for dehydration and air dry. Put the slices into the boiled pretreatment solution (100 ± 5 °C) for 25 min. Soak in PBS for 2×5 min. Followed by protein digestion process at 37 °C for 20 min, wash in PBS for 2×5 min. Dehydration step was performed in 75%, 85% and 100% alcohol for 1 min each. DNA FISH was performed as described previously and images were captured on the ZEISS LSM 800 Inverted Confocal microscope attached with an Airyscan area detector.

**Immunofluorescence combined with DNA FISH**

Immunofluorescence was performed as previously described. After incubating the cells with the secondary antibodies, cells were washed three times in PBS for 5 min at RT and fixed with 4% PFA in PBS for 10 min, washed three times in PBS. Cells were incubated in 70%, 85% and then 100% ethanol for 1 min at RT. 10 µL of probe hybridization mixture was added on a slide and coverslip was placed on top. The following steps of DNA FISH were as previously described and images were acquired on the ZEISS LSM 800 Inverted Confocal microscope attached with an Airyscan area detector.

**Live-cell-imaging microscopy**

To monitor DNase1L3 and lipid droplets dynamics within the cells, we transiently expressed DNase1L3-EGFP, DNase1L3$^{\Delta NT}$-EGFP, DNase1L3$^{3\times NT}$-EGFP, and GC-DNase-EGFP for 48 h and co-stained the cells with Hoechst and 20 µg/mL BODIPY-558/568 for 45 min followed by washing 3 times in PBS, and then performed imaging experiments. Cells were imaged at the Structured light illumination microscope (Zeiss Elyra7) pre-stabilized at 37 °C for 2 h. We illuminated the sample with 488-, 561- and 405-nm laser 60× Oil lens, Z-stack images were acquired with a 0.3 µm z-step size with 3 second intervals between each volumetric imaging. For DNase1L3 and lipid droplets colocalization analysis, a straight line was drawn across the center of the objects in a 2D plane and the fluorescent intensity was profiled along the line path.

**Image analysis**

For fluorescence intensity statistics, measurements were made using the Profile and Measure functions of the ZEN Blue software. For the number and area statistics of fluorescent signal points, as well as the statistics of wound healing area and number of clones were measured using the Analyze Particles function of ImageJ software. For the analysis of overlapping signals of two channels, after importing the original image using ImageJ software and splitting it by channel, selecting the Min algorithm in the image calculation module, following by selecting the images of the two channels to be analyzed, confirming and generating the images of the two overlapping channels, and then measuring the overall intensity.

**Immunohistochemical (IHC) staining**

The tissue samples were dissected in PBS, the tissues were fixed overnight in 4% formalin, embedded in paraffin, 4 μm sectioned, and stained with hematoxylin and eosin (H&E) for pathological examination. Tissue slides were deparaffinized in xylene and rehydrated in a graded series of ethanol. The slides were immersed in sodium citrate buffer and boiled for 25 min for antigen retrieval. After blocking endogenous peroxidase activity with 0.3% hydrogen peroxide and blocking nonspecific protein binding with goat serum. Incubate the section with the primary antibody overnight at 4 °C. Next, incubate the section with the peroxidase-conjugated secondary antibody, rinse with PBS Three times for 5 min each, add DAB solution dropwise for 6 min, and terminate in water color rendering. Tissue slides are photographed to obtain images. Images were performed for IHC scoring, to be specific, the degree of staining (0-3 points) and the rate of positivity (0-4 points) in the immunohistochemical sections were scored separately and multiplied to obtain a comprehensive score (0-12 points). The staining intensity was scored as follows: 0 for no staining, 1 for light yellow, 2 for brownish yellow, and 3 for brownish brown; 0-5% was scored as 0, 6%-25% as 1, 26%-50% as 2, 51%-75% as 3, and 75% as 4.

**RNA-seq**

Total RNA was extracted by RNeasy mini kit (Qiagen) for sequencing with TruSeq RNA Library Prep Kit v2 (Illumina) according to the manufacturer’s instruction. In brief, total

RNA was firstly processed with poly-A selection and fragmentation, and then first-and second-strand cDNA was synthesized and ligated with sequencing adaptor. The library was finally sequenced in high throughput on the Illumina Hiseq platform in 2×150 bp pair-end sequencing mode to obtain FastQ data. Data were processed following the TCGA mRNA analysis pipeline. Expression level of mRNA was computed as FPKM for cell line samples and TCGA samples.

**Circle-Seq library preparation and sequencing**

EcDNA sequencing Service was provided by Shanghai Jiayin Biotechnology (Shanghai, China) by following the published procedures (10.1038/protex.2019.006). In brief, MagAttract HMW DNA Kit (Qiagen, Hilden, Germany, 67563) was applied for genomic DNA extraction. To eliminate linear chromosomal DNA and purify circular DNA, the genomic DNA was treated with Plasmid-Safe ATP-dependent DNase (Lucigen) at 37 °C, after 6 days of digestion, heat inactivate the exonuclease by incubating at 70 °C for 30 min and then extracted with PCI solution (25:24:1). To confirm elimination of linear chromosomal DNA, the sensitivity of the extracted ecDNA to Plasmid-Safe ATP-dependent DNase was conducted. Amplify the remaining circular DNA in the reaction by MDA using φ29 DNA polymerase and random hexamer primers (REPLI-g Mini Kit (Qiagen, Hilden, Germany, 150025)), followed by sample clean up (Agencourt AMPure XP magnetic beads (Beckman Coulter, Brea, CA, USA, A63880)). The samples were used for DNA Library Preparation and Next Generation Paired-End Sequencing using Illumina MiSeq Technology.

**Identification of ecDNA by combined WGS and Circle-seq analysis**

ecDNA was identified following the published literature(*4, 5*) Specifically, we applied Circle-Map software to detect ecDNA from sequencing data to obtain the bed file. Firstly, sequencing reads were aligned to the human reference genome (hg38 genome download from UCSC) using BWA-MEM(RRID:SCR_017619). Then, to detect the coordinates of each ecDNA, two BAM files which were sorted according to read names and coordinates for the extraction of circular reads were used. Several filtering steps were conducted to improve the accuracy of ecDNA detection. The specific settings were as follows: (1) split reads ≥ 2, (2) circle score ≥ 20, (3) coverage increase in the start coordinate ≥ .33, (4) coverage increase in the end coordinate ≥ .33, (5) coverage continuity ≤ .1 and (6) The SD of coverage smaller than the mean coverage over the whole ecDNA region. The matched WGS data were firstly compared with the reference genome using the BWA software package to obtain the BAM file, and then the copy number was calculated using the Cnvkit software. the genomic region of CN ≥ 5 was extracted, and using the intersect function of Bedtools software was used to take the intersection with the bed file obtained by Circle-seq to get the sequence of ecDNA.

**Molecular Dynamics Simulations**

Vitamin D binding protein was downloaded from RCSB PDB data bank (http://www.rcsb.org, PDB ID:1J78, RRID:SCR_012820). After obtaining the 3D structures of the protein and Vitamin D molecular, the molecular docking process was performed with Rosetta8. Based on a maximum number of 100 conformers extracted from precise docking, we obtained the final conformer owing the lowest binding energy.

For simulations of the protein and aptamer systems, the AMBER FF14SB and AMBER PARM99 force fields were used respectively. Each system (the protein−protein complex) was solvated in TIP3P water and NaCl to neutralize the systems. All-atom MD simulations were carried out by using the GROMACS 2019.03 package. The systems were restrained by a harmonic potential of the form $k\ (\Delta x)^2$ where the force constant k was 100 $kcal/mol^{-1}Å^{-2}$. Then, we optimized the water molecules and counter ions using the steepest descent and conjugate gradient for 2500 steps separately. After that, we optimized the entire systems without any constraint using the first step method, following by an annealing simulation process under a weak restraint ($k=100\ kcal/mol^{-1}Å^{-2}$). The complex systems were heated from 0 to 298 K gradually over 500 ps in the NVT ensemble (i.e., N represents the number of particles, V represents the volume and T represents the temperature of the systems, and their product was a constant). After this heating phase, we performed a 40 ns MD simulation under 1 atm with a constant temperature at 298 K and a constant pressure maintained by isotropic position scaling algorithm with a relaxation time of 2 ps.

**Animal studies**

Animal experiments were carried out in accordance with the National Institutes of Health Animal Use Guidelines. All of the experimental protocols have been approved by the Institutional Animal Care and Use Committee.

**In vivo orthotopic implantation models:** Female BALB/c mice aged 4–6 weeks were used in the experiments. In brief, $1\times10^6$ H22-luc cells, or H22-Luc cells transfected with plasmid of DNase1L3-pcDNA3.1+/C-(K)-DYK, DNase1L3$^{\Delta NT}$-pcDNA3.1+/C-(K)-DYK, DNase1L3$^{3\times NT}$-pcDNA3.1+/C-(K)-DYK, GC-pcDNA3.1+/C-(K)-DYK, DNASE1L3 ko_eSpCas9-2A-Puro(PX459)V2.0, ko GC_eSpCas9-2A- Puro (PX459)V2.0 were suspended in 100 µL of Sanitary saline respectively and injected orthotopically into the left liver lobe of BALB/c mice. Tumor growth was detected in vivo after injection of 150 mg/kg D-Luciferin and potassium salt substrate (Yeasen) in PBS into anesthetized mice with NightOWLIILB983 (Berthold Technologies). The mice were killed and the liver were preserved in 10% formalin solution for further study. For VD treatment experiments, mice transplanted with H22-luc cells were randomly divided into three groups, which were control group, VD low-dose group and VD high-dose group. mice in the control group had anhydrous ethanol added to their daily drinking water, and in the VD low-dose group, anhydrous ethanol-dissolved VD3, 600 IU/kg was added to their daily drinking water. in the VD high-dose group, anhydrous ethanol-dissolved VD3, 1500 IU/kg was added to their daily drinking water. the drug was administered continuously for 2 months. For other experiments involving VD administration, VD was treated in the same way and at the same dose as the VD high-dose group, while the other groups of mice were given anhydrous ethanol as a control.

**Diethylnitrosamine (DEN)-induced mouse models:** 2-week-old C57BL/6 mice were given a single intraperitoneal injection of DEN (25 mg/kg body weight). After one week, the DEN-challenged mice were given Carbon tetrachloride (0.5 mg/kg body weight) by intraperitoneal injections once a week for 5 months. Mice were randomly divided into two groups of 6 mice each. The mice in the experimental group were injected with $1\times10^{11}$ virus particles of AAV8-encapsulated ko DNase1L3 plasmid in 200 µL PBS in the tail vein, followed by two booster injections at an interval of less than 1 week, the control mice were injected with 200 µL PBS in the tail vein. After 10 months, the livers of mice were analyzed for number and size of hepatic tumors.

**PDX mouse models:** 8 fresh surgical tumor tissues (F0, hepatocellular carcinoma) were collected immediately after surgery without any other treatment from 8 patients in Tianjin Medical University Cancer Institute and Hospital. The information of the patients is provided in Table S4. A written informed consent was obtained from each patient. According to the Declaration of Helsinki, studies were performed after approval of the ethics committee of Nankai University, including the use of animal experiments and tumor specimens. Tumor fragments were implanted subcutaneously into the right axilla of 4-6-week-old BALB/c nude mice. By palpation of the skin at the tumor site, we selected mice that bore tumor nodules and began to measure the tumor volumes. When the tumor size reached 100 to 200 $mm^3$, the samples (F1) were divided into pieces for in vivo passaging to construct F2 and then F3 tumors as described above. When the F3 tumor size reached 100 to 200 $mm^3$, the mice bearing different tumor types were randomly divided into control, DNase1L3 mRNA and GC-DNase mRNA groups consisting of eight mice per group. mRNA of DNase1L3 and GC-DNase were synthesized by GenScript, the mRNA is encapsulated by LNP and contains the 5' end-modified base AAGCTT, the 3' end-modified base ACTAGT, and a Poly(A) tail modification at the 3' end consisting of 100 adenosine nucleotides. LNP encapsulation of mRNA is performed by Nuohai Life Science (Shanghai) Co, LTd. 100 μL LNP containing 3 μg mRNA was injected intravenously to BALB/c nude mice every 5 days for 10 times. Mice in the control group were injected intravenously with 100 μL LNP solution.

**Bioinformatics analysis**

The clinical information and gene expression data of hepatocellular carcinoma in The Cancer Genome Atlas (TCGA) were downloaded using the R package "TCGAbiolinks" and analyzed by the R package "ggplot2", "survival" and "survminer". Gene set enrichment analysis (GSEA, RRID:SCR_003199) was carried out using R package "clusterProfiler".

**Statistical analysis**

Statistical analyses were performed using GraphPad Prism version 9 for Windows or R 4.1.3. Statistically significant differences were calculated using two-tailed unpaired t-tests, or unpaired t test with Welch's correction, Pearson's correlation, and Kaplan–Meier as needed. $P<0.05$ was considered significant.

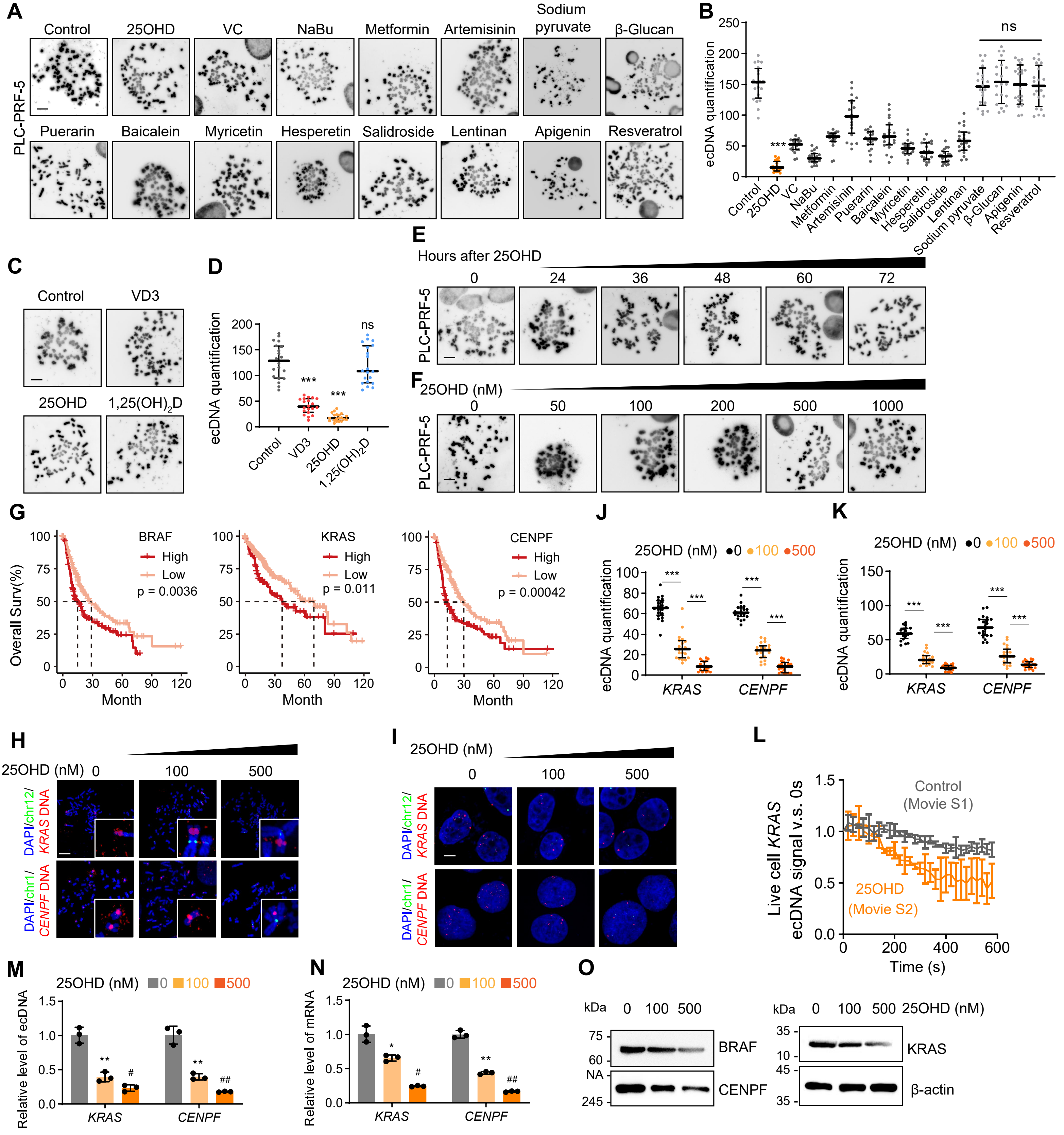

A
Control
25OHD
VC
NaBu
Metformin
Artemisinin
Sodium pyruvate
β-Glucan
PLC-PRF-5
Puerarin
Baicalein
Myricetin
Hesperetin
Salidroside
Lentinan
Apigenin
Resveratrol
B
ecDNA quantification
ns
***
C
Control
VD3
25OHD
1,25(OH)2D
D
ecDNA quantification
ns
E
Hours after 25OHD
0
24
36
48
60
72
F
25OHD (nM)
0
50
100
200
500
1000
G
BRAF
High
Low
p = 0.0036
KRAS
p = 0.011
CENPF
p = 0.00042
Overall Surv(%)
Month
J
25OHD (nM) ●0 ●100 ●500
ecDNA quantification
KRAS
CENPF
K
H
25OHD (nM) 0 100 500
DAPI/chr12/KRAS DNA
DAPI/chr1/CENPF DNA
I
L
Control
(Movie S1)
25OHD
(Movie S2)
Live cell KRAS ecDNA signal v.s. 0s
Time (s)
M
25OHD (nM) 0 100 500
Relative level of ecDNA
N
Relative level of mRNA
O
kDa
0 100 500
25OHD (nM)
BRAF
CENPF
KRAS
β-actin
NA

**Figure S1. VD has prominent effects on reducing ecDNA level in HCC cells, related to Figure 1**

**(A)** Representative images of metaphase ecDNA in PLC-PRF-5 cells after treatment of DMSO, 25OHD (500 nM), Vitamin C (150 μM), Sodium butyrate (1 mM), Metformin (5 mM), Artemisinin (15 μM), Puerarin (25 μM), Baicalein (3 μM), Myricetin (15 μM), Hesperetin (5 μM), Salidroside (3 mM), Lentinan (1 mM), Sodium pyruvate (20 mM), β-Glucan (400 μM), Apigenin (30 μM), and Resveratrol (100 μM) for 48 h. Scale bar, 10 μm.

**(B)** Quantification of metaphase ecDNA by ecSeg in PLC-PRF-5 cells. 20 metaphase spreads from 3 biologically independent samples were counted. Statistics were calculated on biological replicates with Wilcoxon rank-sum test. ***P < 0.001, compared with Control. Error bars show median, upper and lower quartiles.

**(C)** Representative images of metaphase ecDNA in PLC-PRF-5 cells after treatment of different forms of VD including VD3 (500 nM), 25OHD (500 nM), 1,25$(OH)_2$D (500 nM) for 48 h.

**(D)** Quantification of metaphase ecDNA by ecSeg in PLC-PRF-5 cells. 20 metaphase spreads from 3 biologically independent samples were counted. Statistics were calculated on biological replicates with Wilcoxon rank-sum test. ***P < 0.001, ns, not significant, compared with Control. Error bars show median, upper and lower quartiles.

**(E)** Raw images of metaphase ecDNA in PLC-PRF-5 cells after treatment of DMSO or 25OHD (500 nM) for different time. Scale bar, 10 μm.

**(F)** Raw images of metaphase ecDNA in PLC-PRF-5 cells after treatment of DMSO or 25OHD for different concentrations for 48 h. Scale bar, 10 μm.

**(G)** Kaplan–Meier curves showing percentage of the overall survival of the high and low expression of BRAF, KRAS, CENPF in the TCGA dataset. (n=371).

**(H-K)** Representative FISH images of metaphase ecDNA (**H**) and interphase ecDNA (**J**) signal in PLC-PRF-5 cells after treatment of DMSO or 25OHD (100 nM or 500 nM) for 48 h. Scale bar, 10 μm. FISH signal quantification by ecSeg were shown in **(I)** and **(K)** respectively. 20 metaphase spreads or interphase cells of FISH images from 3 biologically independent samples were counted. Statistics were calculated on biological replicates with Wilcoxon rank-sum test. ***P < 0.001. Error bars show median, upper and lower quartiles.

**(L)** The average signal intensity of KRAS ecDNA in live cells by using CRISPR live FISH after treatment of DMSO (movie S1) or 25OHD (movie S2).

**(M)** QPCR detected of ecDNA levels of *KRAS*, *CENPF* purified and isolated from PLC-PRF-5 cells treated with DMSO or 25OHD (100 nM or 500 nM) for 48 h, n=3, biological replicates. Statistics were calculated on biological replicates with two-tailed unpaired t-tests. $^*P < 0.05$, $^{**}P < 0.01$, compared with control; #$P < 0.05$, ##$P < 0.01$, compared with VD (100 nM). Error bars show mean with SD.

**(N)** Quantification of mRNA levels of *KRAS*, *CENPF* by RT-qPCR in PLC-PRF-5 cells treated with DMSO or 25OHD (100 nM or 500 nM) for 48 h. Student's t-test, n=3, biological replicates. Statistics were calculated on biological replicates with two-tailed unpaired t-tests. $^*P < 0.05$, compared with control; #$P < 0.05$, ##$P < 0.01$, compared with VD (100 nM). Error bars show mean with SD.

**(O)** The levels of BRAF, KRAS, CENPF protein in PLC-PRF-5 cells treated with DMSO or 25OHD (100 nM or 500 nM) for 48 h were detected by western blot.

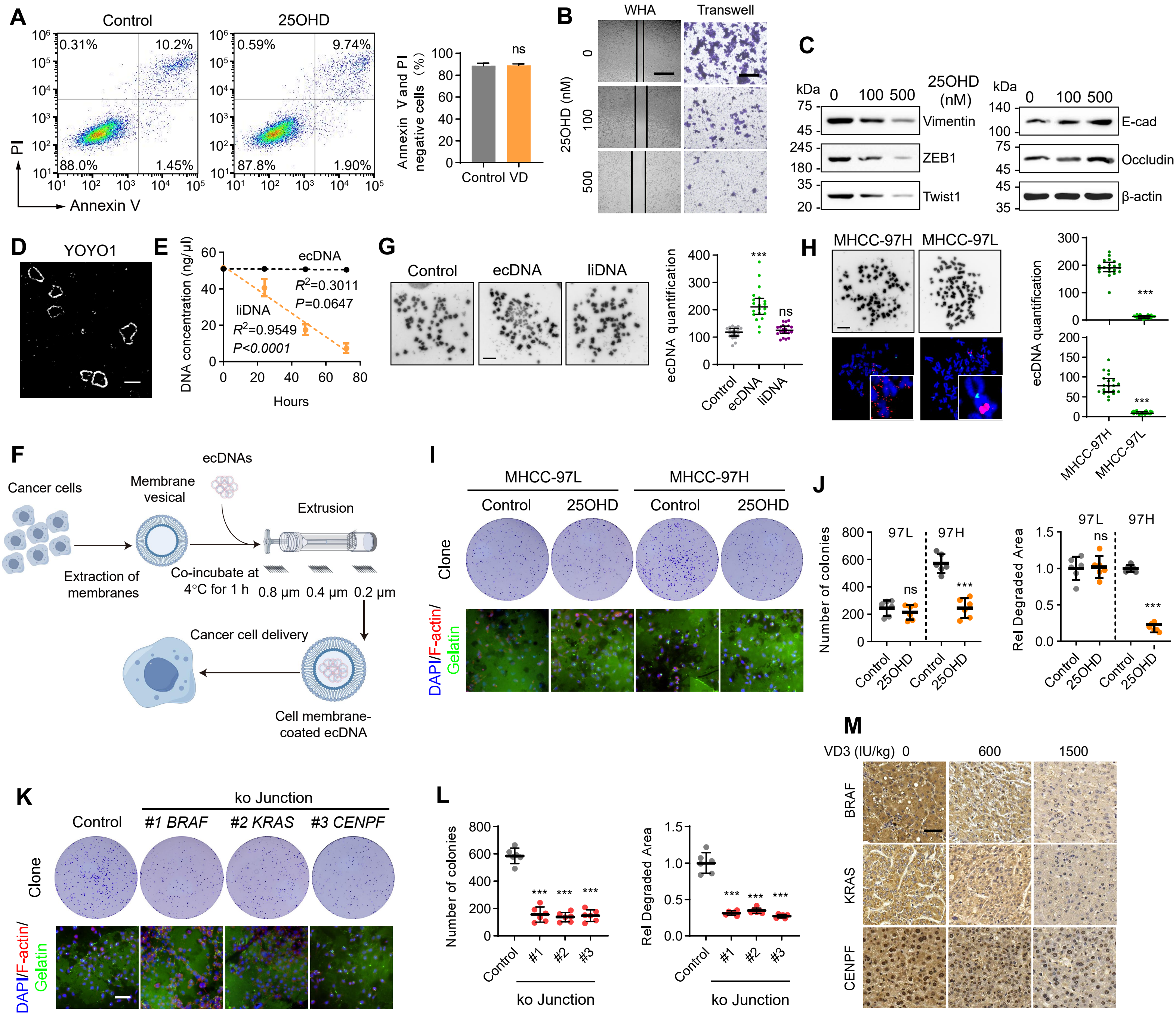

A
Control
25OHD
0.31%
10.2%
88.0%
1.45%
0.59%
9.74%
87.8%
1.90%
PI
Annexin V
Annexin V and PI negative cells (%)
ns
Control VD
B
WHA
Transwell
25OHD (nM)
0
100
500
C
kDa
0 100 500
25OHD (nM)
Vimentin
ZEB1
Twist1
E-cad
Occludin
β-actin
D
YOYO1
E
DNA concentration (ng/μl)
ecDNA
$R^2$=0.3011
P=0.0647
liDNA
$R^2$=0.9549
P<0.0001
Hours
G
Control
ecDNA
liDNA
ecDNA quantification
***
ns
H
MHCC-97H
MHCC-97L
ecDNA quantification
***
***
F
ecDNAs
Cancer cells
Membrane vesical
Extrusion
Extraction of membranes
Co-incubate at 4°C for 1 h
0.8 μm 0.4 μm 0.2 μm
Cancer cell delivery
Cell membrane-coated ecDNA
I
MHCC-97L
MHCC-97H
Control
25OHD
Control
25OHD
Clone
DAPI/F-actin/Gelatin
J
97L
97H
Number of colonies
ns
***
Rel Degraded Area
ns
***
Control
25OHD
K
ko Junction
Control
#1 BRAF
#2 KRAS
#3 CENPF
Clone
DAPI/F-actin/Gelatin
L
Number of colonies
Rel Degraded Area
***
Control
#1
#2
#3
ko Junction
M
VD3 (IU/kg) 0 600 1500
BRAF
KRAS
CENPF

**Figure S2. VD inhibiting the malignant progression of HCC depends on the degradation of ecDNA, related to Figure 2**

**(A)** The PLC-PRF-5 cells did not undergo significant apoptosis after 25OHD (500 nM,48 h) treatment by Annexin V-PI flow cytometry, n=3, biological replicates. Statistics were calculated on biological replicates with two-tailed unpaired t-tests, ns, not significant.

**(B)** Raw images of wound healing assay and Transwell assay of PLC-PRF-5 cells treated with DMSO or 25OHD (100 nM or 500 nM) for 48 h related to Figs. 2b-c. respectively. Scale bar, 200 μm (top), 500 μm (bottom).

**(C)** The levels of EMT related proteins were detected by western blot in PLC-PRF-5 cells treated with DMSO or 25OHD for 48 h.

**(D)** Reresentative SIM image of ecDNA staining with YOYO1, Scale bar, 2 μm.

**(E)** ecDNA purified from PLC-PRF-5 cells or linearized ecDNA (liDNA), treated with ATP-dependent DNase at 37 °C for 0, 24, 48 and 72 h. The concentration of DNA samples showed that ecDNA was insensitive to ATP-dependent DNase, while the concentration of liDNA decreased in a time-dependent manner after ATP-dependent DNase treatment. n=3, biological replicates. Statistics were calculated on biological replicates with simple linear regression. Error bars show mean with SD.

**(F)** Schematic diagram of ecDNA delivery to cancer cells.

**(G)** Representative images (left) and quantification (right) of metaphase ecDNA by ecSeg in PLC-PRF-5 cells after delivery with ecDNA or liDNA for 48 h. Scale bar, 10 μm. 20 metaphase spreads from 3 biologically independent samples were counted. Statistics were calculated on biological replicates with Wilcoxon rank-sum test. ***$P < 0.001$, ns, not significant, compared with Control. Error bars show median, upper and lower quartiles.

**(H)** Representative images of DAPI staining (top) and FISH (bottom) for *KRAS* of metaphase ecDNA in MHCC-97H/L cells, where *KRAS* in MHCC-97L was amplified on HSR, while in MHCC-97H it was amplified in the form of ecDNA. Corresponding ecDNA quantifications by ecSeg were also given (right). 20 metaphase spreads stained with DAPI or FISH images from 3 biologically independent samples were counted. Statistics were calculated on biological replicates with Wilcoxon rank-sum test. ***$P < 0.001$. Error bars show median, upper and lower quartiles.

**(I and J)** (**I**) Representative images of colony formation (top) and fluorescent gelatin degradation and phalloidin/DAPI staining (bottom) of MHCC-97H/L cells treated with

DMSO or 25OHD (500nM) for 48 h and (**J**) corresponding quantification. Scale bar, 40 µm. n=6, biological replicates. Statistics were calculated on biological replicates with two-tailed unpaired t-tests, ***P < 0.001, compared with MHCC-97H Control; ns, not significant, compared with MHCC-97L Control. Error bars show mean with SD.

**(K and L)** (**K**) Representative images of colony formation (top) and fluorescent gelatin degradation and phalloidin/DAPI staining (bottom) of PLC-PRF-5 cells transfected with CRISPR/Cas9-sgRNA plasmids targeting *BRAF*, *KRAS*, *CENPF* respectively, For Colony formation assay, after 48 h of transfection, cells were cultured in a new medium for a total of 14 days. For fluorescent gelatin degradation assay, phalloidin/DAPI staining was performed after 48 h of transfection. The corresponding quantifications were shown in (**L**). n=6, biological replicates. Scale bar, 40 µm. n=6, biological replicates. Statistics were calculated on biological replicates with two-tailed unpaired t-tests, *P < 0.05, ***P < 0.001, compared with Control. Error bars show mean with SD.

**(M)** Raw images of immunohistochemical staining of BRAF, KRAS, CENPF related to fig 2N. Scale bar, 50 µm.

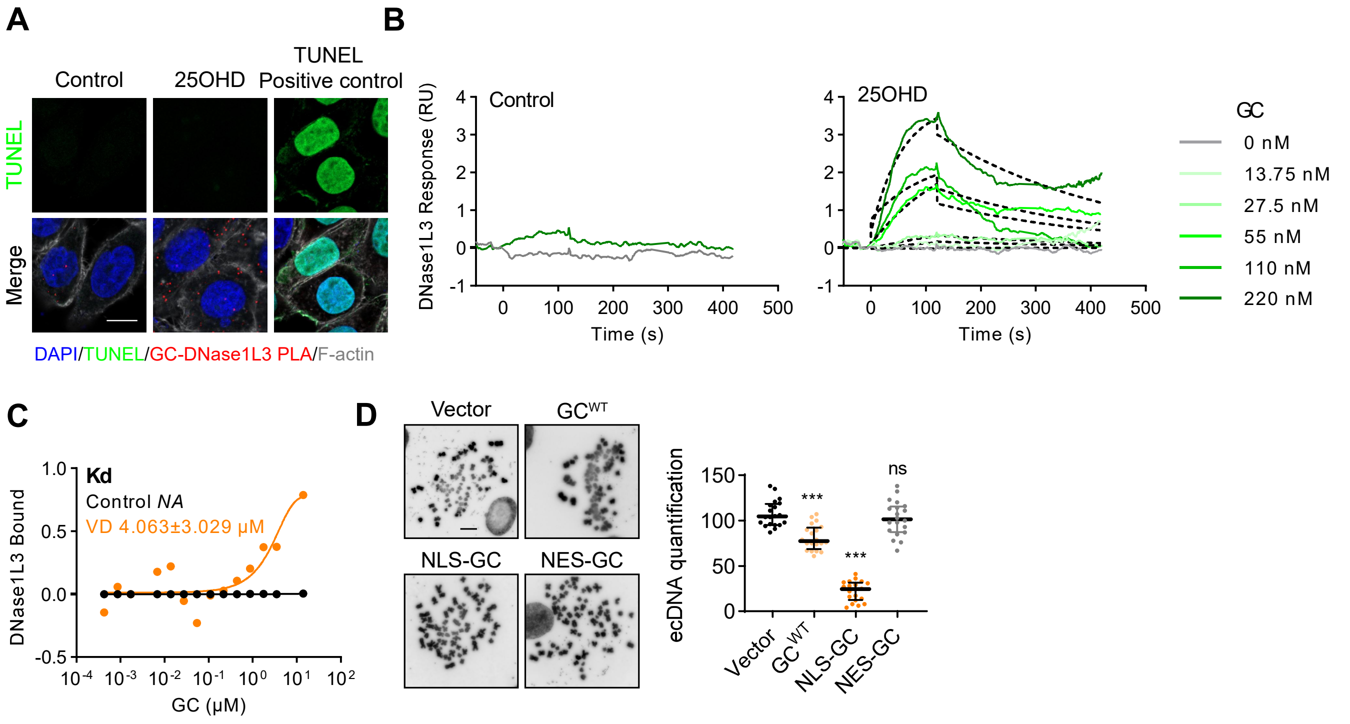

A
Control
25OHD
TUNEL
Positive control
TUNEL
Merge
DAPI/TUNEL/GC-DNase1L3 PLA/F-actin
B
Control
25OHD
DNase1L3 Response (RU)
Time (s)
GC
0 nM
13.75 nM
27.5 nM
55 nM
110 nM
220 nM
C
Kd
Control NA
VD 4.063±3.029 μM
DNase1L3 Bound
GC (μM)
D
Vector
GCWT
NLS-GC
NES-GC
ecDNA quantification
***
***
ns
Vector
GC WT
NLS-GC
NES-GC

**Figure S3. VD enhance the interaction of GC and DNase1L3, related to Figure 3**

**(A)** Representative images of PLA assays in HCC cells with GC and DNase1L3, cells were stained with TUNEL to detect apoptosis. Scale bar, 10 μm.

**(B)** Surface plasmon resonance (SPR) showed that 25OHD (500 nM) facilitates the interaction of GC with DNase1L3.

**(C)** Microscale thermophoresis (MST) experiments showed that 25OHD (500 nM) facilitates the interaction of GC with DNase1L3.

**(D)** Representative images of metaphase ecDNA in PLC-PRF-5 cells after transfected with $GC^{WT}$, NLS-GC or NES-GC for 48 h. **The** Quantification of ecDNA was shown on the right penal. 20 metaphase spreads from 3 biologically independent samples were counted. Statistics were calculated on biological replicates with Wilcoxon rank-sum test. ***P < 0.001, ns, not significant, compared with vector. Error bars show median, upper and lower quartiles.

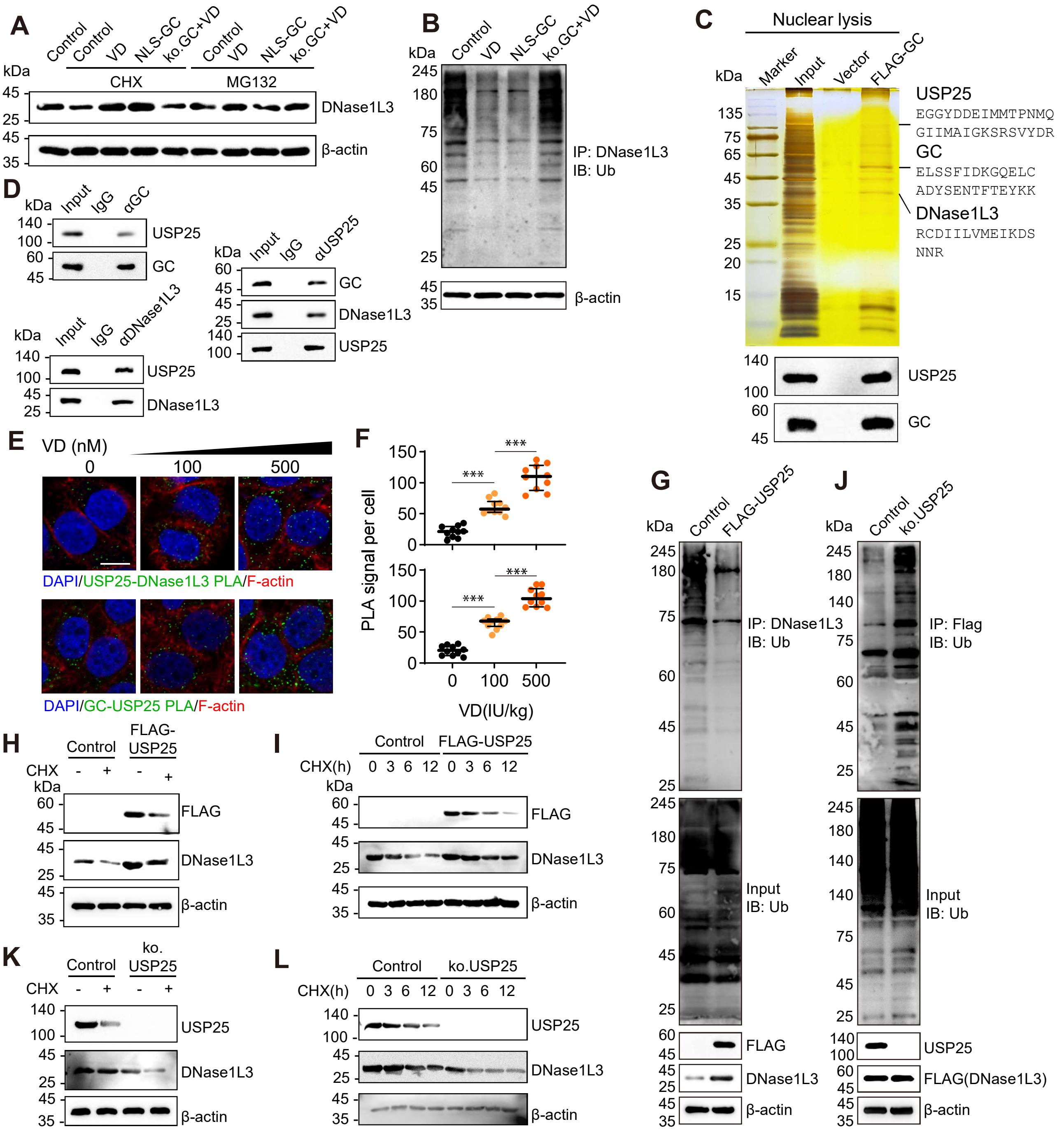
A
Control
VD
NLS-GC
ko.GC+VD
CHX
MG132
kDa
DNase1L3
β-actin
B
IP: DNase1L3
IB: Ub
C
Nuclear lysis
Marker
Input
Vector
FLAG-GC
USP25
EGGYDDEIMMTPNMQ
GIIMAIGKSRSVYDR
GC
ELSSFIDKGQELC
ADYSENTFTEYKK
DNase1L3
RCDIILVMEIKDS
NNR
D
IgG
αGC
αUSP25
αDNase1L3
E
VD (nM)
0
100
500
DAPI/USP25-DNase1L3 PLA/F-actin
DAPI/GC-USP25 PLA/F-actin
F
PLA signal per cell
VD(IU/kg)
***
G
FLAG-USP25
IP: DNase1L3
IB: Ub
Input
IB: Ub
FLAG
J
ko.USP25
IP: Flag
IB: Ub
FLAG(DNase1L3)
H
CHX
I
CHX(h)
0 3 6 12 0 3 6 12
K
L

**Figure S4. GC interacts with USP25 to promote DNase1L3 deubiquitination, related to Figure 3**

**(A)** Effects of proteasome inhibitor MG132 (5 µM, 48 h) or cycloheximide (CHX) (50 µg/mL, 48 h) and indicated treatments on the expression level of DNase1L3 protein by Western blot assay.

**(B)** Effects of indicated treatment on ubiquitination of DNase1L3 in PLC-PRF-5 cells analyzed by in vivo ubiquitination assays.

**(C)** Silver staining (top) of nuclear lysis and western blot analysis (bottom) of USP25 and GC expression level in GC overexpression PLC-PRF-5 cells.

**(D)** Co-IP analysis of interaction of USP25 and GC or USP25 and DNase1L3 in PLC-PRF-5 cells.

**(E)** PLA assays in PLC-PRF-5 cells with USP25 and DNase1L3 (top), USP25 and GC (bottom). Scale bar,10 µm.

**(F)** Quantification of PLA signals. n=10, biological replicates. Statistics were calculated on biological replicates with two-tailed unpaired t-tests. ***$P < 0.001$. Error bars show median, upper and lower quartiles.

**(G)** Overexpression of FLAG-USP25 impaired ubiquitination of DNase1L3 in PLC-PRF-5 cells by in vivo ubiquitination assays.

**(H)** Western blot showed that overexpression of FLAG-USP25 increased DNase1L3 protein levels in PLC-PRF-5 Cells treated with CHX (50 µg/mL) or DMSO.

**(I)** FLAG-USP25 expression increased DNase1L3 protein half-life in PLC-PRF-5 cells. Cells were treated with CHX (50 µg/mL) or DMSO for different hours before Western blot assays.

**(J)** Knocking out of endogenous USP25 increase ubiquitination of DNase1L3 in PLC-PRF-5 cells by in vivo ubiquitination assays.

**(K)** Western blot showed that knock out of USP25 decreased DNase1L3 protein levels in PLC-PRF-5 Cells treated with CHX (50 µg/mL) or DMSO.

**(L)** Knock out of endogenous USP25 decreased DNase1L3 protein half-life (right) in PLC-PRF-5 cells. Cells were treated with CHX (50 µg/mL) or DMSO for different hours before Western blot assays.

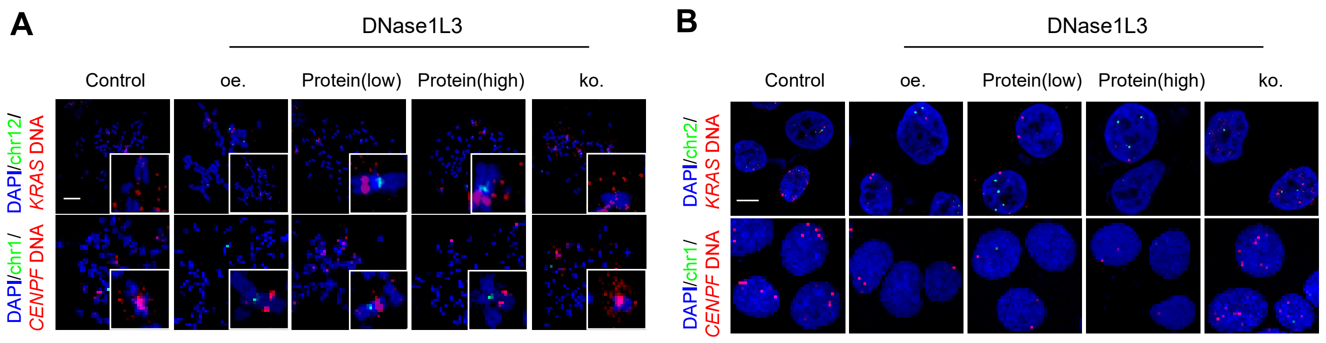

A
DNase1L3
Control
oe.
Protein(low)
Protein(high)
ko.
DAPI/chr12/
KRAS DNA
DAPI/chr1/
CENPF DNA
B
DNase1L3
Control
oe.
Protein(low)
Protein(high)
ko.
DAPI/chr2/
KRAS DNA
DAPI/chr1/
CENPF DNA

**Figure S5. Representative FISH images in Fig. 4G-H, related to Figure 4**

**(A)** Representative FISH images of metaphase ecDNA signal in PLC-PRF-5 cells after indicated treatments in Fig. 4G, Scale bar,10 μm. oe: over expression of DNase1L3 plasmid, Protein(low): transfection of DNase1L3 protein (2 μg), Protein(high): transfection of DNase1L3 protein (10 μg), ko: transfection of DNase1L3 ko plasmid.

**(B)** Representative FISH images of interphase ecDNA signal in PLC-PRF-5 cells after indicated treatments in Fig. 4H, Scale bar,10 μm.

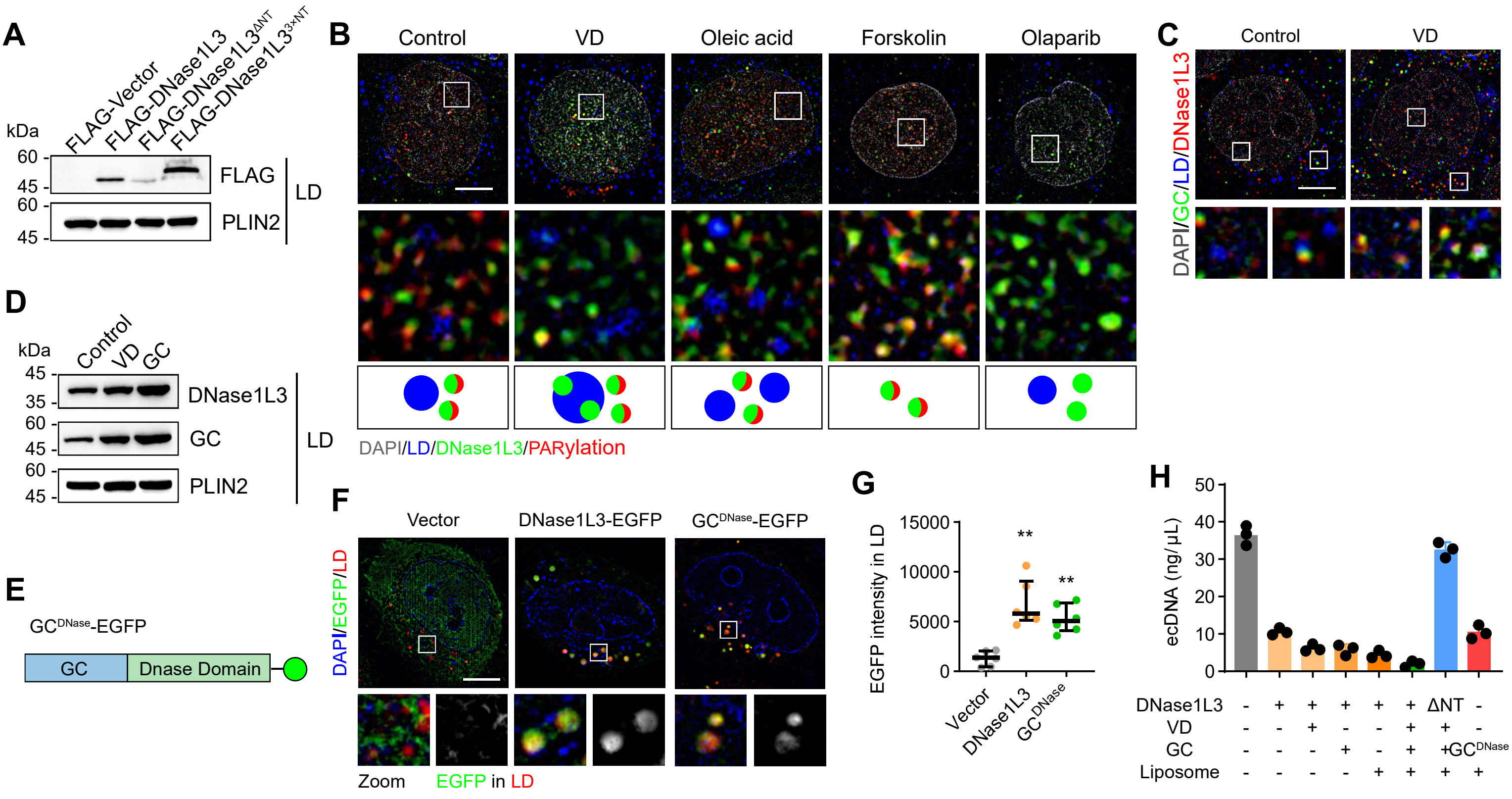

A
FLAG-Vector
FLAG-DNase1L3
FLAG-DNase1L3ΔNT
FLAG-DNase1L33×NT
kDa
60
45
60
45
FLAG
PLIN2
LD
B
Control
VD
Oleic acid
Forskolin
Olaparib
DAPI/LD/DNase1L3/PARylation
C
Control
VD
DAPI/GC/LD/DNase1L3
D
Control
VD
GC
kDa
45
35
60
45
60
45
DNase1L3
GC
PLIN2
LD
E
GCDNase-EGFP
GC
Dnase Domain
F
Vector
DNase1L3-EGFP
GCDNase-EGFP
DAPI/EGFP/LD
Zoom
EGFP in LD
G
EGFP intensity in LD
15000
10000
5000
0
**
**
Vector
DNase1L3
GCDNase
H
ecDNA (ng/μL)
50
40
30
20
10
0
DNase1L3 - + + + + + ΔNT -
VD - - + - - + + -
GC - - - + - + +GCDNase
Liposome - - - - + + + +

**Figure S6. VD promotes GC to induce DNase1L3 targeting LD to escape poly (ADP-ribosyl) ation, related to Figure 5**

**(A)** Western blot analysis of DNase1L3 protein level in LD component after transfection of different constructs of DNase1L3.

**(B)** Representative SIM images of Immunostaining of DNase1L3 (green), PARylation (red) and BODIPY (blue) in PLC-PRF-5 cells in PLC-PRF-5 cells after DMSO, 25OHD (500 nM), oleic acid (200 μM), forskolin (10 μM) or PARP1 inhibitor Olaparib (20 μM, 48 h) treatments for 48 h. after indicated treatments, corresponding zoom views of the overlap of DNase1L3, PARylation and lipid droplets were shown in the bottom panel. Scale bar: 5 μm.

**(C)** Representative SIM images of Immunostaining of DNase1L3 (red), GC (green) and BODIPY (blue) in PLC-PRF-5 cells treated with DMSO or 25OHD (500 nM) for 48 h, corresponding zoom views of the overlap of DNase1L3, GC and lipid droplets were shown in the bottom panel. Scale bar: 5 μm.

**(D)** Western blot analysis of DNase1L3 and GC protein level in LD component after transfection of vector or GC plasmid in PLC-PRF-5 cells.

**(E)** Schematic of the $GC^{DNase}$ construct used.

**(F)** Representative live-cell SIM images of EGFP-labelled DNase1L3, EGFP-labelled $GC^{DNase}$, and Bodipy staining of lipid droplets (top) and corresponding zoom views of the overlap images (bottom) in PLC-PRF-5 cells after indicated treatments.

**(G)** Quantification of the lipid droplets containing EGFP intensity in PLC-PRF-5 cells after transfected with indicated plasmids. n=6, biological replicates. Statistics were calculated on biological replicates with unpaired t test with two-tailed unpaired t-tests. $^{**}P < 0.01$, compared with control. Error bars show median, upper and lower quartiles.

**(H)** Quantification of purified ecDNA incubated with the indicated treatments for 2 h. DNase1L3 (10 ng/μL), $DNase1L3^{\Delta NT}$ (10 ng/μL), $DNase1L3^{3\times NT}$ (10 ng/μL), GC (10 ng/μL), 25OHD (500 nM), liposome (5 μL), the incubation was performed in a 60 μL reaction volume in the presence of 5 mM $MgCl_2$, 2.5 mM $CaCl_2$ at 37°C. n=3, biological replicates.

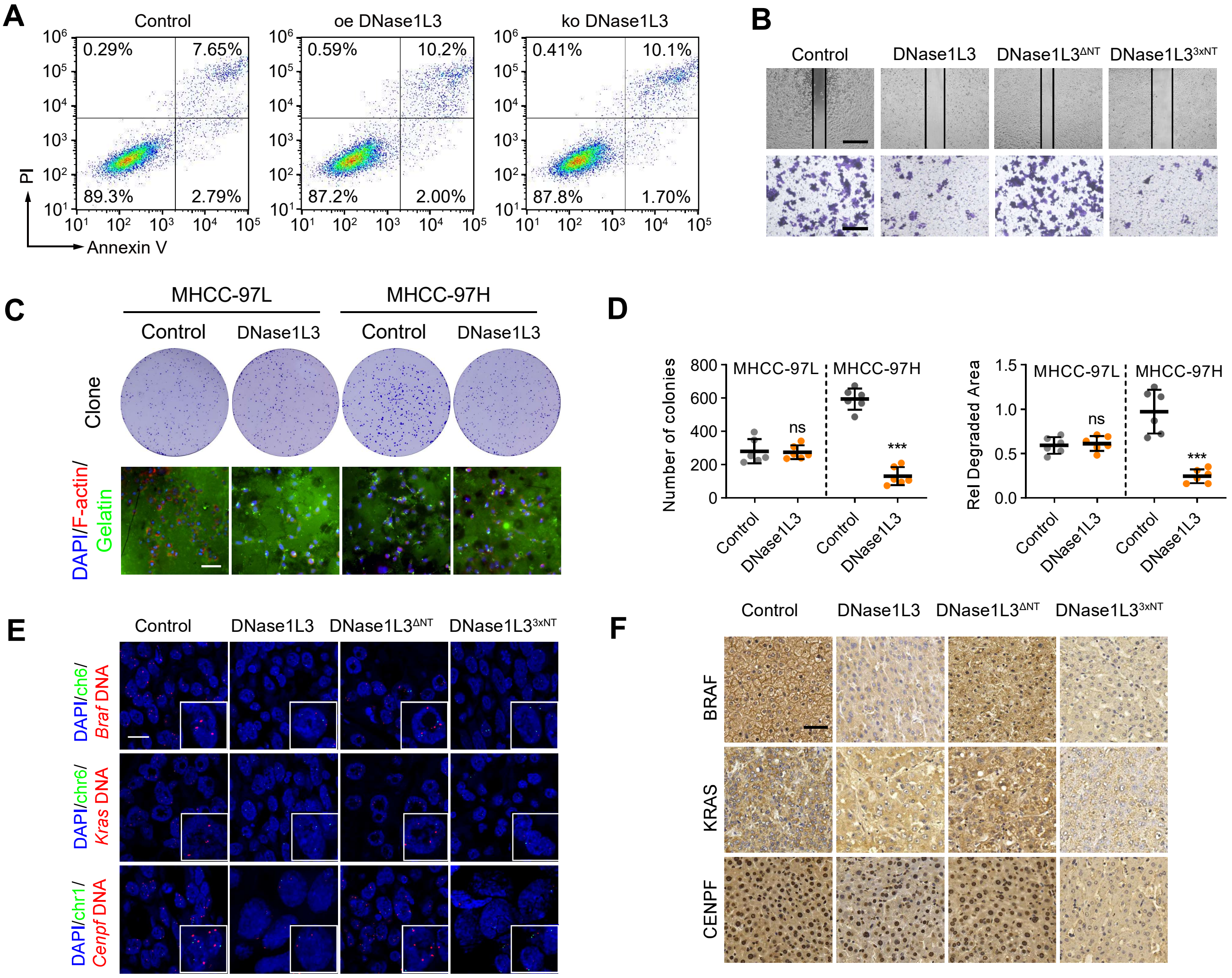
A
Control
oe DNase1L3
ko DNase1L3
0.29%
7.65%
89.3%
2.79%
0.59%
10.2%
87.2%
2.00%
0.41%
10.1%
87.8%
1.70%
PI
Annexin V
B
Control
DNase1L3
DNase1L3ΔNT
DNase1L33xNT
C
MHCC-97L
MHCC-97H
Control
DNase1L3
Control
DNase1L3
Clone
DAPI/F-actin/
Gelatin
D
Number of colonies
MHCC-97L
MHCC-97H
ns
***
Control
DNase1L3
Control
DNase1L3
Rel Degraded Area
MHCC-97L
MHCC-97H
ns
***
Control
DNase1L3
Control
DNase1L3
E
Control
DNase1L3
DNase1L3ΔNT
DNase1L33xNT
DAPI/ch6/
Braf DNA
DAPI/chr6/
Kras DNA
DAPI/chr1/
Cenpf DNA
F
Control
DNase1L3
DNase1L3ΔNT
DNase1L33xNT
BRAF
KRAS
CENPF

**Figure S7. DNase1L3 degrades ecDNA to inhibit the malignant progression of HCC and raw images, related to Figure 6**

**(A)** After overexpression of DNase1L3 in PLC-PRF-5 cells, the cells were still able to proliferate, Annexin V-PI flow cytometry showed that the overexpression of DNase1L3 did not lead to cell apoptosis.

**(B)** Raw images of wound healing assay (top) and Transwell assay (bottom) of PLC-PRF-5 cells treated as indicated related to Figs. 6b-c respectively. Scale bar, 500 μm (top). 200 μm (bottom).

**(C and D)** (**C**) Representative images of colony formation (top) and fluorescent gelatin degradation and phalloidin/DAPI staining (bottom) of MHCC-97H/L cells treated with DMSO or 25OHD (500 nM, 48 h) and (**D**) corresponding quantification. Scale bar, 40 μm. n=6, biological replicates. Statistics were calculated on biological replicates with two-tailed unpaired t-tests, ***$P < 0.001$, compared with MHCC-97H Control; ns, not significant, compared with MHCC-97L Control. Error bars show mean with SD.

**(E)** Representative images of DNA FISH for liver tissues in H22-luc-tumor-bearing-BALB/c mice treated as indicated. Scale bar, 10 μm.

**(F)** Representative images of immunohistochemical staining of BRAF, KRAS, CENPF. Scale bar, 50 μm.

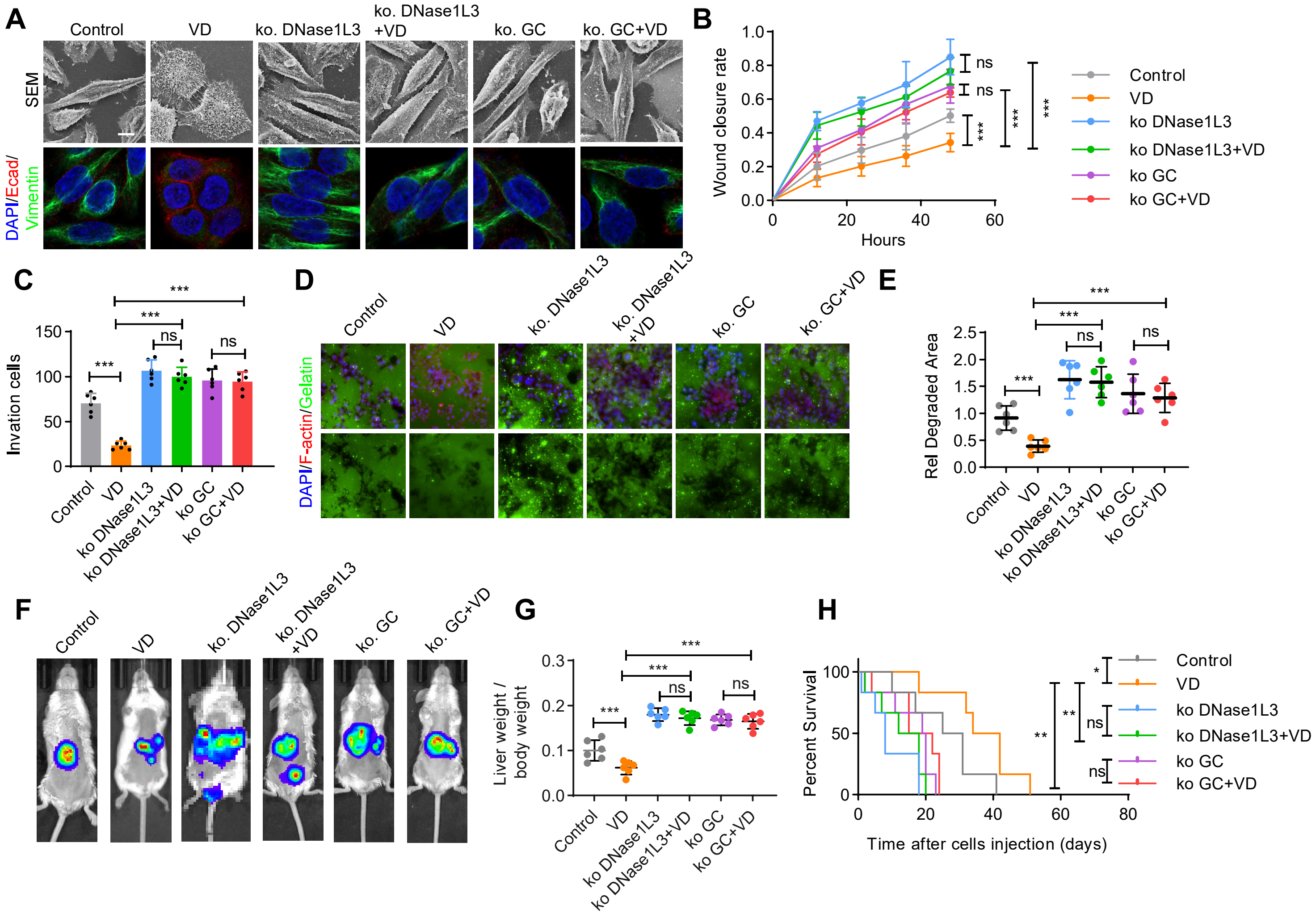

A
Control
VD
ko. DNase1L3
ko. DNase1L3 +VD
ko. GC
ko. GC+VD
SEM
DAPI/Ecad/Vimentin
B
Wound closure rate
Hours
Control
VD
ko DNase1L3
ko DNase1L3+VD
ko GC
ko GC+VD
ns
***
C
Invation cells
D
DAPI/F-actin/Gelatin
E
Rel Degraded Area
F
G
Liver weight / body weight
H
Percent Survival
Time after cells injection (days)

**Figure S8. The anti-tumor effect of VD depends on DNase1L3 and GC, related to Figure 6**

**(A)** The images of scanning electron microscopy (top) and immunostaining (bottom) of E-cadherin and vimentin signals in PLC-PRF-5 cells treated as indicated. Scale bar, 10 µm.

**(B and C)** Quantification of wound healing assay (**B**) and Transwell assay (**C**) of PLC-PRF-5 cells treated as indicated. n=3, biological replicates. Statistics were calculated on biological replicates with two-tailed unpaired t-tests, ***P < 0.001, ns, not significant. Error bars show mean with SD.

**(D and E)** Fluorescent gelatin degradation and phalloidin/DAPI staining of PLC-PRF-5 cells treated as indicated (**D**) and quantification of degradation area (**E**). Scale bar, 40 µm. *n*=6, biological replicates. Statistics were calculated on biological replicates with two-tailed unpaired t-tests, ***P < 0.001, ns, not significant. Error bars show mean with SD.

**(F and G)** Representative in vivo images (**F**) of H22-luc-tumor-bearing-BALB/c mice with transplanted H22-luc cells transfection of indicated plasmids and liver weight/body weight ratio was quantified (**G**). *n*=6, biological replicates. Statistics were calculated on biological replicates with two-tailed unpaired t-tests, ***P < 0.001. Error bars show median, upper and lower quartiles.

**(H)** Kaplan–Meier curves showing percentage of survival of H22-luc-tumor-bearing-BALB/c mice in f. *n*=6, biological replicates. Statistics were calculated on biological replicates with Kaplan–Meier analysis. *P<0.05, **P < 0.01, ns, not significant.

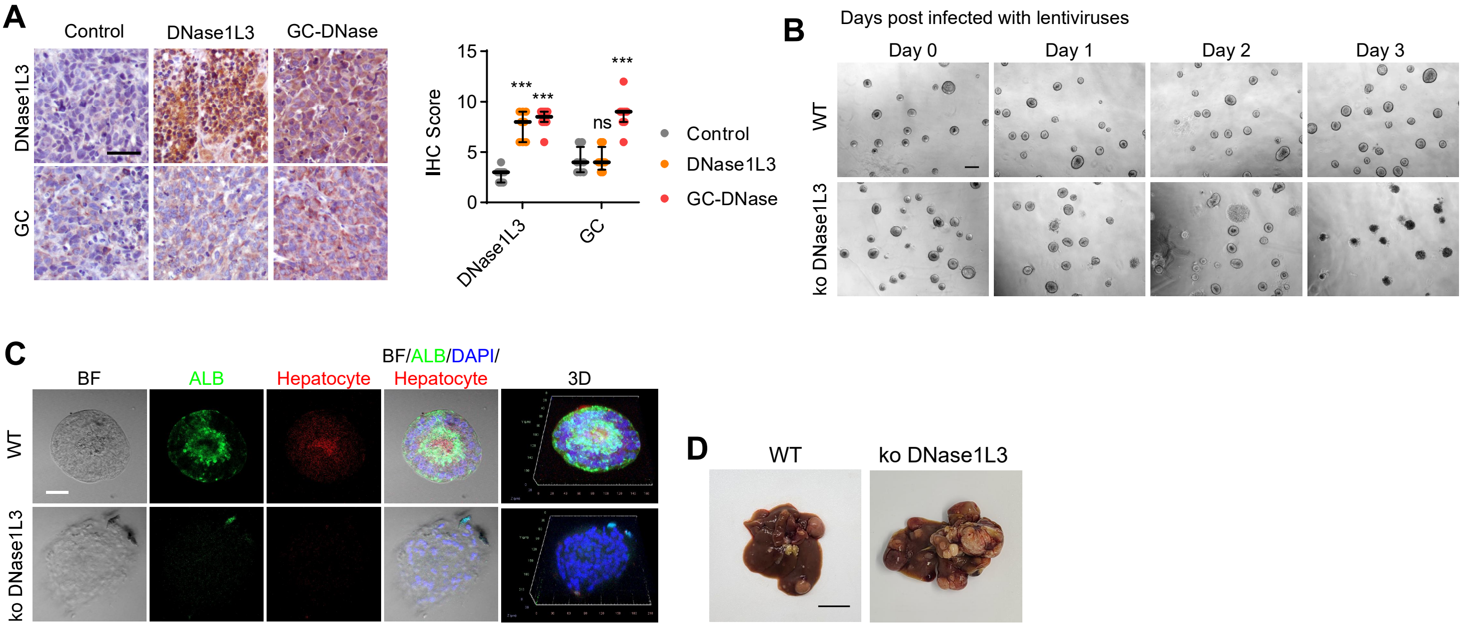

A
Control
DNase1L3
GC-DNase
DNase1L3
GC
IHC Score
15
10
5
0
***
***
ns
***
DNase1L3
GC
Control
DNase1L3
GC-DNase
B
Days post infected with lentiviruses
Day 0
Day 1
Day 2
Day 3
WT
ko DNase1L3
C
BF
ALB
Hepatocyte
BF/ALB/DAPI/
Hepatocyte
3D
WT
ko DNase1L3
D
WT
ko DNase1L3

**Figure S9. Raw images of iPSC-derived liver organoid and liver tissues of DEN-induced mouse model, related to Figure 7**

**(A)** Immunohistochemical staining of DNase1L3 and GC in liver tissues of PDX mice to verify to successful of mRNA delivery. n=8, biological replicates. Statistics were calculated on biological replicates with unpaired t-test with Welch's correction, ***$P < 0.001$, ns, not significant, compared with Control. Error bars show median, upper and lower quartiles.

**(B)** Brightfield microscopy images of iPSC-Derived Liver Organoids after infected with lentiviruses of vector or sgRNA targeting DNase1L3 based on CRISPR/cas9 system. Scale bar, 200 μm.

**(C)** Representative confocal images of ALB (green) and hepatocyte (red) in iPSC-Derived Liver Organoids after infected with lentiviruses of vector or sgRNA targeting DNase1L3 based on CRISPR/cas9 system. Scale bar, 40 μm.

**(D)** Representative images of the visible liver nodules of diethylnitrosamine (DEN)-induced mouse model of HCC. Scale bar, 0.5 cm.

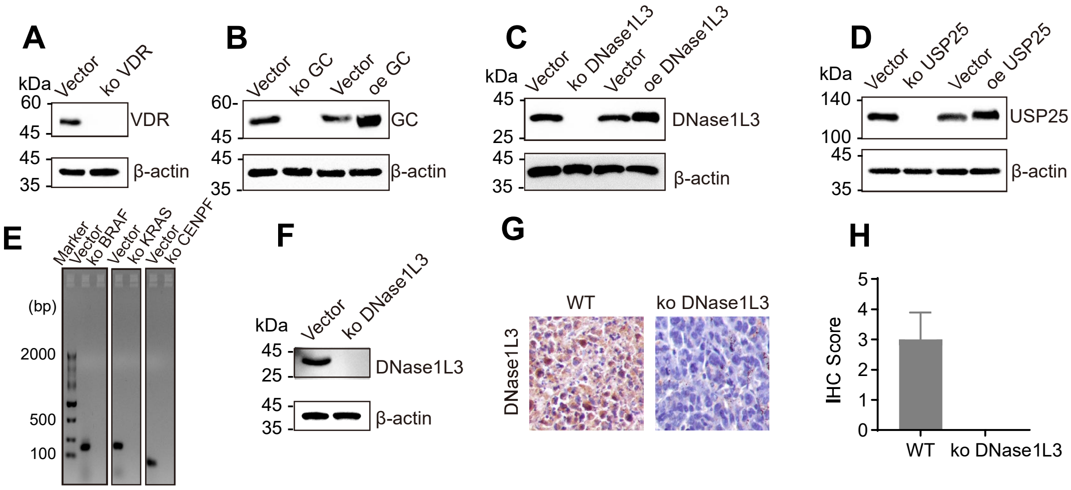

A
kDa
Vector
ko VDR
60
45
VDR
45
35
β-actin
B
Vector
ko GC
Vector
oe GC
60
45
GC
45
35
β-actin
C
kDa
Vector
ko DNase1L3
Vector
oe DNase1L3
45
25
DNase1L3
45
35
β-actin
D
kDa
Vector
ko USP25
Vector
oe USP25
140
100
USP25
45
35
β-actin
E
Marker
Vector
ko BRAF
Vector
ko KRAS
Vector
ko CENPF
(bp)
2000
500
100
F
kDa
Vector
ko DNase1L3
45
25
DNase1L3
45
35
β-actin
G
WT
ko DNase1L3
DNase1L3
H
IHC Score
5
4
3
2
1
0
WT
ko DNase1L3

**Figure S10. Western blotting for verifying knockout of VDR, GC, DNase1L3, and USP25 and overexpression of GC, DNase1L3, and USP25，related to Figure 7 and STAR Method.**

**(A)** CRISPR Knockout Plasmid containing Cas9 and guide RNAs of VDR was transfected into PLC-PRF-5 cells for 48 h followed by puromycin selection, the successful knockout of VDR protein was validated by western blot.

**(B)** CRISPR Knockout Plasmid containing Cas9 and guide RNAs of GC and the GC over expression plasmid were transfected into PLC-PRF-5 cells for 48 h respectively, followed by puromycin selection of cells with GC CRISPR Knockout Plasmid transfection, the successful knockout or over expression of GC was validated by western blot.

**(C)** CRISPR Knockout Plasmid containing Cas9 and guide RNAs of DNase1L3 and the DNase1L3 over expression plasmid were transfected into PLC-PRF-5 cells for 48 h respectively, followed by puromycin selection of cells with DNase1L3 CRISPR Knockout Plasmid transfection, the successful knockout or over expression of DNase1L3 was validated by western blot.

**(D)** CRISPR Knockout Plasmid containing Cas9 and guide RNAs of USP25 and the USP25 over expression plasmid were transfected into PLC-PRF-5 cells for 48 h respectively, followed by puromycin selection of cells with USP25 CRISPR Knockout Plasmid transfection, the successful knockout or over expression of USP25 was validated by western blot.

**(E)** Successful knockout of *BRAF*, *KRAS*, *CENPF* ecDNA verified by PCR of ecDNA junction followed by agarose gel electrophoresis.

**(F)** CRISPR Knockout Plasmid containing Cas9 and guide RNAs of DNase1L3 was transfected into liver organoid for 48 h followed by puromycin selection, the successful knockout of DNase1L3 protein was validated by western blot.

**(G-H)** Knockout of DNase1L3 in diethylnitrosamine (DEN)-induced mouse model of HCC was validated by IHC and corresponding IHC score statistics, n=6, biological replicates.

#### Supplementary tables and Movies

**Table S1. ecDNA junction sequence**

| | Gene | Junction sequence (5'-3') |
|---|---|---|
| **#1** | BRAF | TTCTTGGCACCTTCATAATG |
| **#2** | KRAS | AAATATAGATGAAGGTACTA |
| **#3** | CENPF | GCATTAGGCAAACAACAACCTGG |

**Table S2. The interacting proteins of the intranuclear GC protein after VD treatment identified by MS (Excel File)**

**Table S3. Patient information in TCGA LIHC dataset (Excel File)**

**Table S4. Patient information of PDX mouse model**

| Patient ID | Age | Sex | Pathology diagnosis | Grade | Stage |
|---|---|---|---|---|---|
| **1** | 39 | M | Hepatocellular carcinoma | II-III | II |
| **2** | 33 | M | Hepatocellular carcinoma | II-III | II |
| **3** | 47 | M | Hepatocellular carcinoma | III | II |
| **4** | 65 | M | Hepatocellular carcinoma | III | II |
| **5** | 52 | M | Hepatocellular carcinoma | II | II |
| **6** | 46 | F | Hepatocellular carcinoma | III | IIIB |
| **7** | 52 | F | Hepatocellular carcinoma | III | I |
| **8** | 47 | M | Hepatocellular carcinoma | II | IIIA |

**Movie S1. KRAS ecDNA live FISH on PLC-PRF-5 cells, related to Figure S1L**
Live FISH imaging was performed on PLC-PRF-5 cells for about 30 min. No significant change was detected in the signal of KRAS ecDNA in the cells

**Movie S2. KRAS ecDNA live FISH after adding VD to PLC-PRF-5 cells, related to Figure S1L**

Live FISH imaging was performed after adding VD to PLC-PRF-5 cells for about 30 min. A gradual decrease in the size and magnitude of the signal of the intracellular KRAS ecDNA was found, related to Figure S1I.
